# Fibroblast-Enhanced Tumour Microenvironment Signalling Promotes Adaptive Doxorubicin Tolerance in Heterotypic Melanoma Spheroids

**DOI:** 10.64898/2026.06.30.735445

**Authors:** Ilie Ovidiu Pavel, Giorgiana-Gabriela Negrea, Szilvia Meszaros, Valentin-Florian Rauca, Bogdan-Razvan Dume, Emilia Licarete, Laura Patras, Stefan Dragan, Vlad Alexandru Toma, Alina Sesarman, Manuela Banciu

## Abstract

Melanoma is an aggressive malignancy that rapidly adapts to therapy. While chemotherapy resistance has traditionally been attributed to tumour-intrinsic mechanisms, growing evidence implicates the tumour microenvironment in shaping drug tolerance. However, few *in vitro* models capture the stromal complexity needed to study this interaction.

We developed two multicellular melanoma spheroid models of increasing stromal complexity: a baseline model of melanoma, endothelial, and macrophage cells (*BEM*), and a fibroblast-containing counterpart (*BEMF*), and compared their transcriptional response to doxorubicin. Fibroblast inclusion increased the doxorubicin concentration required to achieve comparable growth inhibition.

While untreated *BEMF* spheroids exhibited only modest baseline transcriptional differences, they showed a profoundly reshaped transcriptional response after doxorubicin exposure, displaying broader and higher-magnitude changes. These responses were characterized by suppression of proliferative and cell-cycle programmes, together with activation of inflammatory, immune-associated, metabolic, and stress-adaptive pathways. Higher-resolution pathway analyses further revealed coordinated attenuation of mitotic progression, checkpoint regulation, homologous recombination repair, and Rho GTPase signalling, consistent with a shift toward stress-adaptive and phenotypically plastic states, rather than classical resistance mechanisms. Transcriptome-derived transcription factor activity inference supported this regulatory rewiring.

Integration with curated resistance-associated genes and external transcriptomic datasets demonstrated strong conservation of core transcriptional features across heterogeneous experimental systems, including consistent suppression of proliferation-associated genes and induction of inflammatory signalling programmes.

Together, these findings indicate that fibroblasts redirect chemotherapy responses toward a stress-adaptive, persister-like phenotype and establish fibroblast-containing 3D melanoma spheroids as a physiologically relevant platform for studying tumour microenvironment-mediated chemotherapy tolerance and stromal-tumour interactions.

## 1. Introduction

Melanoma is among the most aggressive solid tumours, driven by its high metastatic capacity and rapid adaptive responses to therapy [1]. While targeted therapies and immune checkpoint inhibitors have markedly improved patient outcomes, intrinsic and acquired resistance to chemotherapy remains a fundamental limitation, particularly in therapy-refractory tumours, where agents such as doxorubicin (DOX) continue to be explored experimentally or as components of combination therapies [2,3]. Recently, increasing evidence has indicated that drug resistance in melanoma extends beyond the tumour-intrinsic genetic alterations, and is strongly influenced by the tumour microenvironment (TME), which aids in regulating survival signalling, transcriptional programs and therapeutic responses [4,5].

Thus, TME as a complex system composed of immune cells, endothelial cells, perivascular cells, and stromal fibroblasts, communicates with tumour cells through soluble factors, extracellular vesicles, extracellular matrix (ECM) remodelling, and direct cell–cell contacts, collectively contributing to the attenuation of drug-induced cytotoxicity [6,7] and activation of adaptive stress-response pathways in cancer cells [8,9].

Among stromal constituents, fibroblasts have emerged as key regulators of therapy resistance across tumours, including melanoma, where tumour-secreted factors promote the activation of fibroblasts into cancer-associated fibroblast (CAF) phenotypes. Activated fibroblasts promote chemoresistance through several mechanisms, including the induction of drug efflux transporters belonging to the ATP-binding cassette family [10]. Fibroblast-derived cytokines and chemokines, including IL-6, CXCL8 (IL-8), and TNF, activate pro-survival signalling cascades such as STAT3, NF-κB, and MAPK pathways, thereby enhancing resistance to therapy [11]. Moreover, excessive deposition and remodelling of ECM components by fibroblasts increase tissue density and stiffness, forming a physical barrier that impairs drug penetration, and establishes biochemical gradients that favour tumour cell adaptation [12,13]. Fibroblasts can also promote epithelial-mesenchymal transition (EMT) and programs associated with stemness, enhancing survival capacity and plasticity under therapy [14,15].

Given this central role of fibroblasts in regulating cancer cell survival and therapy response the aim of this study was to investigate the contribution of fibroblasts to tumour microenvironment–driven signalling networks that confer resistance to DOX in melanoma, using a heterocellular 3D model (spheroids) that closely mimics TME intercellular crosstalk.

Despite recent advances, most preclinical *in vitro* models fail to fully capture the complexity of tumour drug resistance [16]. Conventional two-dimensional (2D) cultures lack the spatial architecture and heterotypic cellular interactions required to capture physiologically relevant drug responses [17]. Tumour spheroids partially model diffusion gradients and cell–cell interactions, but often lack key stromal components present *in vivo*, limiting their ability to model TME driven resistance mechanisms [18,19]. Therefore, there is a relevant need for advanced 3D models that address stromal complexity and more accurately model therapeutic resistance.

Thus, endothelial cells and macrophages were included in the spheroids alongside fibroblasts because they represent key stromal regulators of vascular function, immune signalling, and chemotherapy response [20,21]. Specifically, endothelial cells influence drug delivery, oxygen and nutrient gradients, and paracrine survival signalling [20], while macrophages contribute to inflammatory and immunosuppressive signals that can reinforce non-mutational drug tolerance [21]. Importantly, fibroblasts modulate the recruitment, activation, and functional polarization of both endothelial cells and macrophages, creating integrated stromal networks that shape intrinsic therapy resistance [21,22]. Inclusion of these cells within the spheroids enables a more physiologically relevant model that captures how fibroblast-driven resistance mechanisms to DOX are amplified or stabilized through stromal cooperation, which are absent in fibroblast–tumour co-cultures alone.

By systematically comparing this sophisticated fibroblast-containing model with a simpler model composed of melanoma, endothelial, and macrophage cells, we evaluated the differential capacity conferred by fibroblast inclusion to activate transcriptional, regulatory, and functional resistance programs upon DOX exposure, using transcriptomic profiling and regulatory network analysis. Fibroblast inclusion broadened and intensified the transcriptional response to DOX, marked by suppression of proliferative and cell-cycle programmes alongside activation of inflammatory and stress-adaptive pathways — a pattern conserved across independent transcriptomic datasets and consistent with a shift toward a stress-adaptive, persister-like state rather than classical drug resistance.

## 2. Results

### 2.1. Fibroblast effects on gene expression in melanoma spheroids

To assess the baseline effect of fibroblast inclusion in melanoma spheroids, *DESeq2* was used to compare the differential expression of genes in *BEMF* and *BEM* spheroids under control conditions. Only a small number of genes exhibited large expression differences: nine genes met the combined threshold of FDR < 0.05 and absolute log₂FC > 1 (Table 1), corresponding to at least a twofold change between models. Although a larger number of genes reached statistical significance overall (790 genes at FDR < 0.05), the vast majority exhibited small effect sizes. This pattern indicates that fibroblast inclusion primarily induces subtle shifts in the basal transcriptome, with only a limited subset of genes showing strong baseline divergence.

**Table 1.** Top differentially expressed genes between *BEMF* and *BEM* spheroids under control conditions.

| Gene Symbol | log<sub>2</sub>FC | FDR-adjusted p-value | Biotype | Description |
| --- | --- | --- | --- | --- |
| Mmp3 | 3.73 | $1.5 \times 10^{-23}$ | Protein coding | Matrix metalloproteinase 3 |
| Gm5537 | 2.33 | $2 \times 10^{-4}$ | Processed pseudogene | Predicted gene 5537 |
| Gm49759 | 1.17 | $2 \times 10^{-53}$ | Uncharacterized | Predicted gene 49759 |
| Malat1 | 1.14 | $5.6 \times 10^{-10}$ | lncRNA | Metastasis-associated lung adenocarcinoma transcript 1 |
| Ier2 | -1 | $2.1 \times 10^{-16}$ | Protein coding | Immediate early response 2 |
| Fos | -1.16 | $2.9 \times 10^{-22}$ | Protein coding | FBJ osteosarcoma oncogene |
| mt-Rnr2 | -1.17 | $1 \times 10^{-45}$ | Mitochondrial rRNA | Mitochondrial 16S rRNA |
| Btg2 | -1.56 | $8.9 \times 10^{-27}$ | Protein coding | BTG anti-proliferation factor 2 |
| Egr1 | -1.78 | $9.5 \times 10^{-07}$ | Protein coding | Early growth response 1 |

The gene showing the largest increase in expression was *Mmp3* (log₂FC = 3.73), a matrix metalloprotease associated with epithelial–mesenchymal transition, extracellular matrix remodelling, and malignant transformation [23]. Additional expression changes were observed in genes with oncogenic regulatory potential, including the lncRNA *Malat1* (log₂FC = 1.14), a well-established oncogenic transcript reported to promote pro-tumorigenic processes (e.g., proliferation, survival, invasion, and metastasis) by modulating tumour-suppressive miRNA networks and enhancing oncogenic signalling [24].

Among the most strongly downregulated genes, *Btg2* (log₂FC = –1.56) and *Egr1* (log₂FC = –1.78) showed reduced expression in untreated *BEMF* compared with untreated *BEM* spheroids. *Btg2* functions as an anti-proliferative tumour suppressor involved in cell cycle control [25], whereas *Egr1* is an early growth response transcription factor that regulates proliferation, differentiation, and stress responses [26].

### 2.2. Fibroblasts increase tolerance of melanoma spheroids to DOX

Quantitative dose–response analysis indicated reduced sensitivity to DOX in fibroblast-containing spheroids compared with spheroids lacking fibroblasts. Spheroids composed of melanoma cells, endothelial cells, and macrophages (*BEM*) exhibited an IC₅₀ of 0.83 µM and an IC₃₀ of 0.56 µM, whereas inclusion of fibroblasts (*BEMF*) increased the IC₅₀ to 1.36 µM and the IC₃₀ to 0.79 µM (Table 2). However, comparison of the dose–response curves using an extra sum-of-squares F-test revealed that this apparent shift did not reach statistical significance (p = 0.53), indicating that the observed differences in IC₃₀ and IC₅₀ were not statistically supported at the curve level (Fig. 1C).

**Figure 1.**
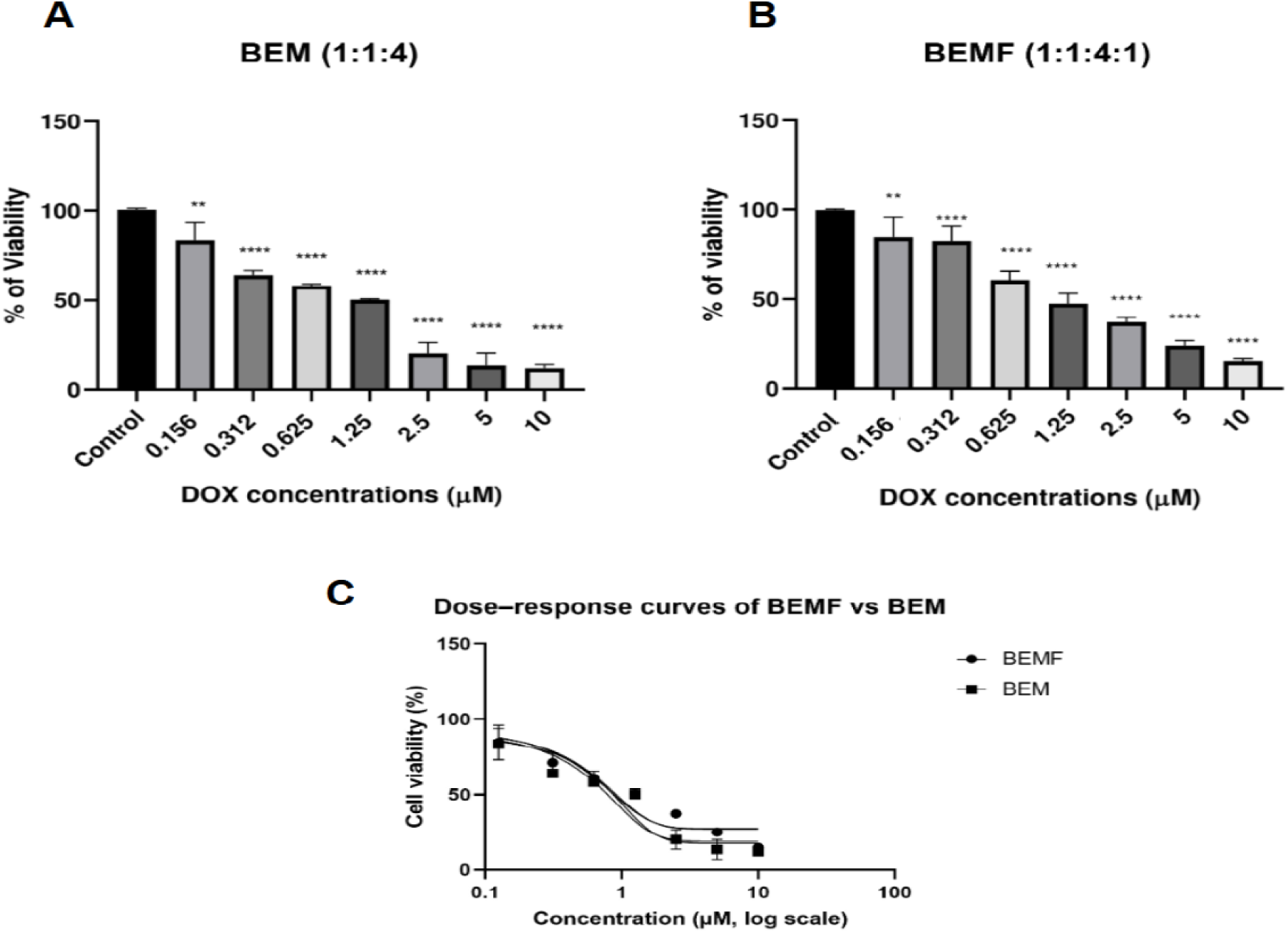
Spheroids viability results. **(A)** Cell viability (%) of melanoma spheroids composed of melanoma cells, endothelial cells, and macrophages (BEM; 1:1:4) following treatment with increasing concentrations of doxorubicin (DOX; 0.156–10 μM), measured after 48 h using the acid phosphatase (APH) assay. **(B)** Cell viability (%) of melanoma spheroids additionally containing fibroblasts (BEMF; 1:1:4:1) under identical treatment conditions. Statistical significance relative to untreated controls was assessed using one-way Anova, followed by Dunnett’s multiple comparisons test, with *p < 0.05, **p < 0.01, ****p < 0.0001. **(C)** Nonlinear regression (four-parameter logistic model) of dose–response curves for BEM and BEMF spheroids. Comparison of fitted curves using an extra sum-of-squares F-test indicated no significant difference between models.

**Table 2.** Inhibitory concentrations of DOX in *BEM* and *BEMF* spheroid models.

| Type of spheroids | DOX IC<sub>50</sub> μM | DOX IC<sub>30</sub> μM |
| --- | --- | --- |
| Spheroids containing murine melanoma cells, endothelial cells and macrophages (*BEM*) | 0.83 | 0.56 |
| Spheroids containing murine melanoma cells, endothelial cells, macrophages and fibroblasts (*BEMF*) | 1.36 | 0.79 |

Notably, fibroblast-containing spheroids (*BEMF*) consistently maintained higher cell viability than fibroblast-free spheroids (*BEM*) across a broad range of DOX concentrations (0.156–10 µM) (Fig. 1A–B). Consistent with this observation, area under the curve (AUC) analysis across the full dose range demonstrated significantly higher overall viability in *BEMF* spheroids compared with *BEM* spheroids (mean AUC: 87.25 vs. 76.59, respectively; Welch’s t-test p = 0.020). This pattern supports a biologically meaningful increase in chemotherapy tolerance associated with fibroblast inclusion, even though variability across replicates limited statistical significance in the global curve comparison. To investigate whether fibroblast inclusion modulated the transcriptional response to DOX, we compared differential expression profiles between *BEMF* and *BEM* spheroids. Using thresholds of FDR < 0.05 and absolute log₂FC > 1, the *BEMF* model displayed a larger number of DOX-responsive genes than the *BEM* model (3,801 vs. 3,281; Fig. 2B).

**Figure 2.**
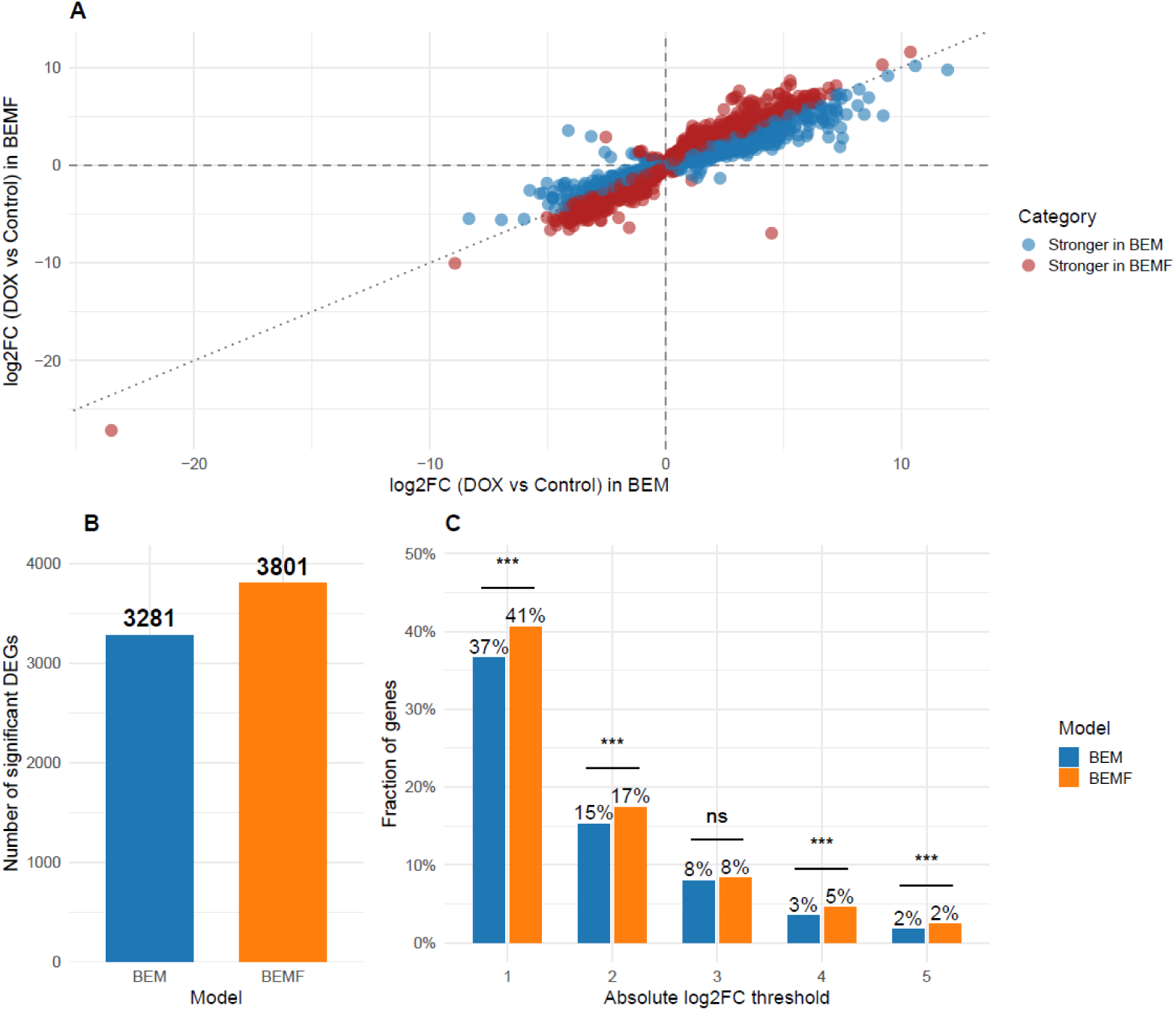
Fibroblast inclusion enhances the magnitude of transcriptional responses to DOX. **(A)** Scatter plot of log₂ fold changes (DOX vs control) in *BEM* (x-axis) versus *BEMF* (y-axis) for genes significantly regulated in both spheroids (FDR < 0.05). Points are coloured according to the model exhibiting the larger absolute effect size, with genes showing stronger regulation in BEM shown in blue and those stronger in *BEMF* shown in red. The diagonal dashed line indicates identical responses between models. A paired t-test on absolute log₂FC values revealed a modest but statistically significant shift toward stronger regulation in *BEMF* (mean |log₂FC|: *BEM* = 1.17, *BEMF* = 1.25; mean paired difference = 0.09; *t* = 13.98, *df* = 7,452, *p* < 2.2 × 10⁻¹⁶). **(B)** Number of significantly DOX-regulated genes (FDR < 0.05 and |log₂FC| > 1) in each model. *BEM* spheroids exhibited 3,281 differentially expressed genes (DEGs), whereas *BEMF* spheroids exhibited 3,801 DEGs. **(C)** Fraction of shared DOX-responsive genes (FDR < 0.05 in both models) exceeding increasing absolute log₂FC thresholds (thresholds of |log₂FC| ≥ 1–5 correspond to ≥2-, 4-, 8-, 16-, and 32-fold expression changes). Bars represent the percentage of genes above each threshold in *BEM* (blue) and *BEMF* (red). Statistical significance of paired differences in proportions was assessed using McNemar’s test (*p* < 0.05; \**p* < 0.01; \*\**p* < 0.001).

Effect-size analysis further revealed that the amplitude of the transcriptional response was increased in *BEMF* spheroids. Among genes significantly regulated in both models, fold-change magnitudes were consistently higher in *BEMF* spheroids, as reflected by the global distribution of log₂FC values (Fig. 2A). A paired t-test confirmed that this shift, while modest in absolute magnitude, was highly significant (mean |log₂FC|: *BEM* ≈ 1.17 vs. *BEMF* ≈ 1.25; p < 2.2 × 10⁻¹⁶). This pattern was observed across a broad range of response magnitudes. Across increasing absolute log₂FC thresholds (1–5), in *BEMF* spheroids, a greater fraction of high-magnitude responders was shown, with differences reaching statistical significance by McNemar’s test for four of five thresholds examined (Fig. 2C).

To independently validate RNA-Seq–derived transcriptional changes in fibroblast-containing spheroids, RT-qPCR was performed for 25 genes spanning inflammatory signalling, drug transport, and proliferation-associated programs (Table S1; Fig. 3). Directional concordance between RT-qPCR and DESeq2 log₂ fold-change estimates was observed for 22 of 25 genes, including key mediator responses to DOX such as *Il6*, *Ptgs2*, *Abcb1a*, *Birc5*, *Tk1*, *Tyms*, and *Pbk* (Table 3). These results confirm robust activation of inflammatory and chemoresistance-associated pathways alongside suppression of proliferative programs in *BEMF* spheroids exposed to DOX.

**Figure 3.**
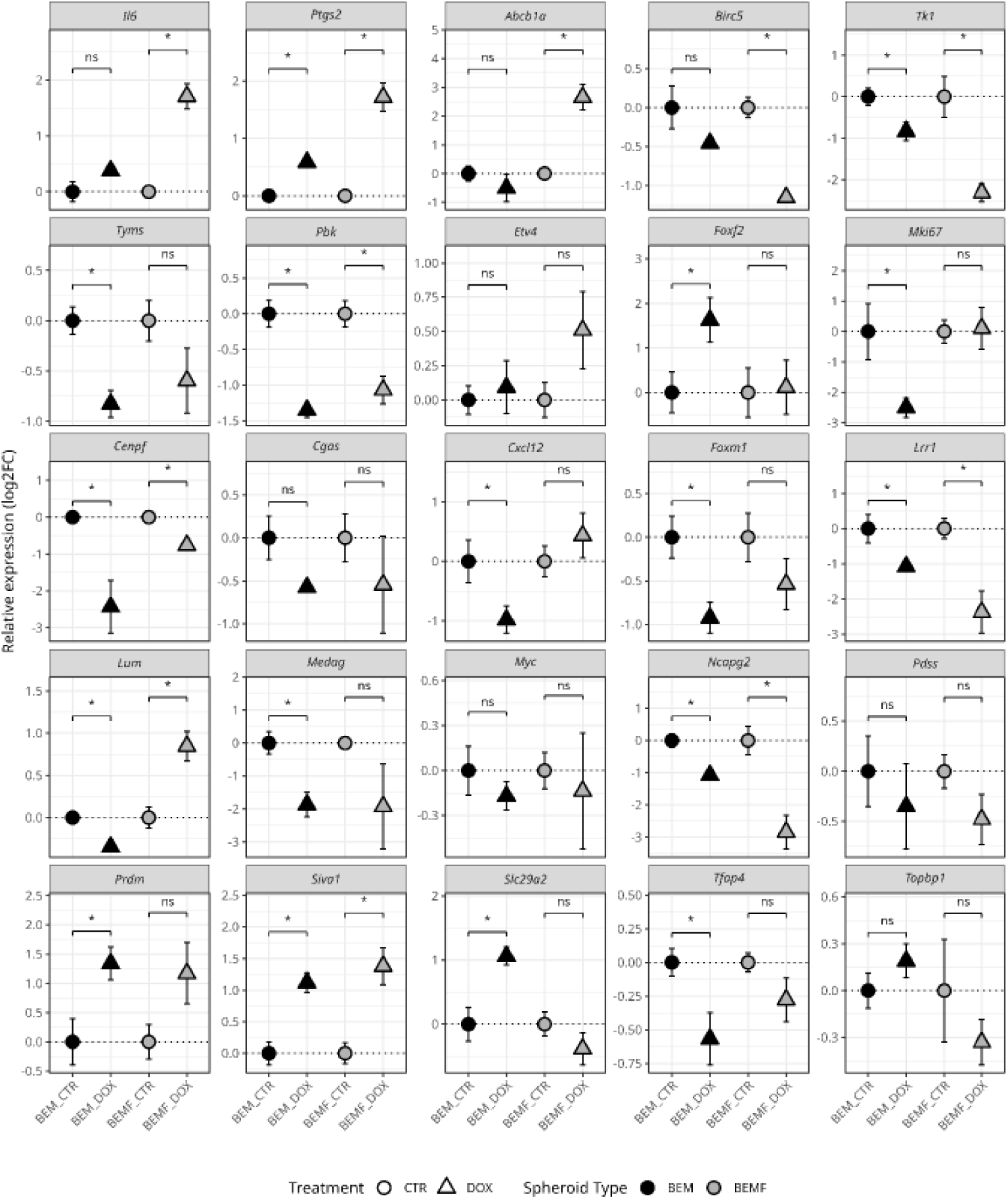
RT–qPCR validation of DOX-responsive gene expression changes in BEM and BEMF spheroids. Relative gene expression changes measured by RT–qPCR in BEM and BEMF spheroids following DOX treatment. Expression values are shown as log2 fold changes relative to the corresponding control group within each cell composition, calculated using the comparative Ct (ΔΔCt) method. For each gene, symbols represent mean values and error bars indicate standard deviation across biological replicates. Comparisons were performed separately within BEM and BEMF spheroids by two-sided Welch’s t-tests on ΔCt values, with p-values adjusted for multiple testing using the Benjamini–Hochberg method. Adjusted significance is indicated as ns, not significant; * adjusted p < 0.05; ** adjusted p < 0.01; *** adjusted p < 0.001; **** adjusted p < 0.0001.

**Table 3.** RT-qPCR Results for DOX-Treated *BEMF* vs. Control.

| Gene | Mean Log2FC | Standard deviation<br>of Log2FC | Direction Match with<br>DESeq2 |
| --- | --- | --- | --- |
| Abcb1a | 2.66 | 0.44 | Yes |
| Birc5 | -1.15 | 0.05 | Yes |
| Cenpf | -0.76 | 0.14 | Yes |
| Cgas | -0.54 | 0.56 | Yes |
| Cxcl12 | 0.43 | 0.37 | Yes |
| Etv4 | 0.51 | 0.28 | No |
| Foxf2 | 0.12 | 0.61 | No |
| Foxm1 | -0.54 | 0.29 | Yes |
| Il6 | 1.71 | 0.22 | Yes |
| Lrr1 | -2.37 | 0.6 | Yes |
| Lum | 0.85 | 0.18 | Yes |
| Medag | -1.93 | 1.29 | Yes |
| Mki67 | 0.1 | 0.68 | No |
| Myc | -0.14 | 0.39 | Yes |
| Ncapg2 | -2.84 | 0.53 | Yes |
| Pdss | -0.48 | 0.25 | Yes |
| Pbk | -1.07 | 0.19 | Yes |
| Prdm1 | 1.17 | 0.52 | Yes |
| Ptgs2 | 1.72 | 0.25 | Yes |
| Siva1 | 1.38 | 0.29 | Yes |
| Slc29a2 | -0.38 | 0.25 | Yes |
| Tfap4 | -0.28 | 0.16 | Yes |
| Tk1 | -2.31 | 0.22 | Yes |
| Topbp1 | -0.33 | 0.14 | Yes |
| Tyms | -0.6 | 0.32 | Yes |

A limited subset of genes (*Etv4*, *Foxf2*, *Mki67*) displayed discordant directionality between RT-qPCR and bulk RNA-Seq, despite exhibiting large-magnitude DESeq2 log₂ fold changes. These discrepancies were confined to transcription factors or cell-cycle–associated genes and did not affect the overall concordance observed for pathway-level responses.

Altogether, these analyses showed that fibroblast-containing spheroids displayed a broader and higher-magnitude transcriptional response to DOX than spheroids without fibroblasts.

### 2.3. Differential responses of *BEMF* and *BEM* spheroids to DOX

To identify transcriptional changes associated with the response of fibroblast-containing melanoma spheroids to DOX, we analysed differential gene expression in both DOX-treated spheroid types. A total of 2,173 genes were significantly affected by the interaction between drug exposure and cell composition (FDR < 0.05), indicating widespread differences in DOX response between *BEM* and *BEMF* spheroids (Fig. 4).

**Figure 4.**
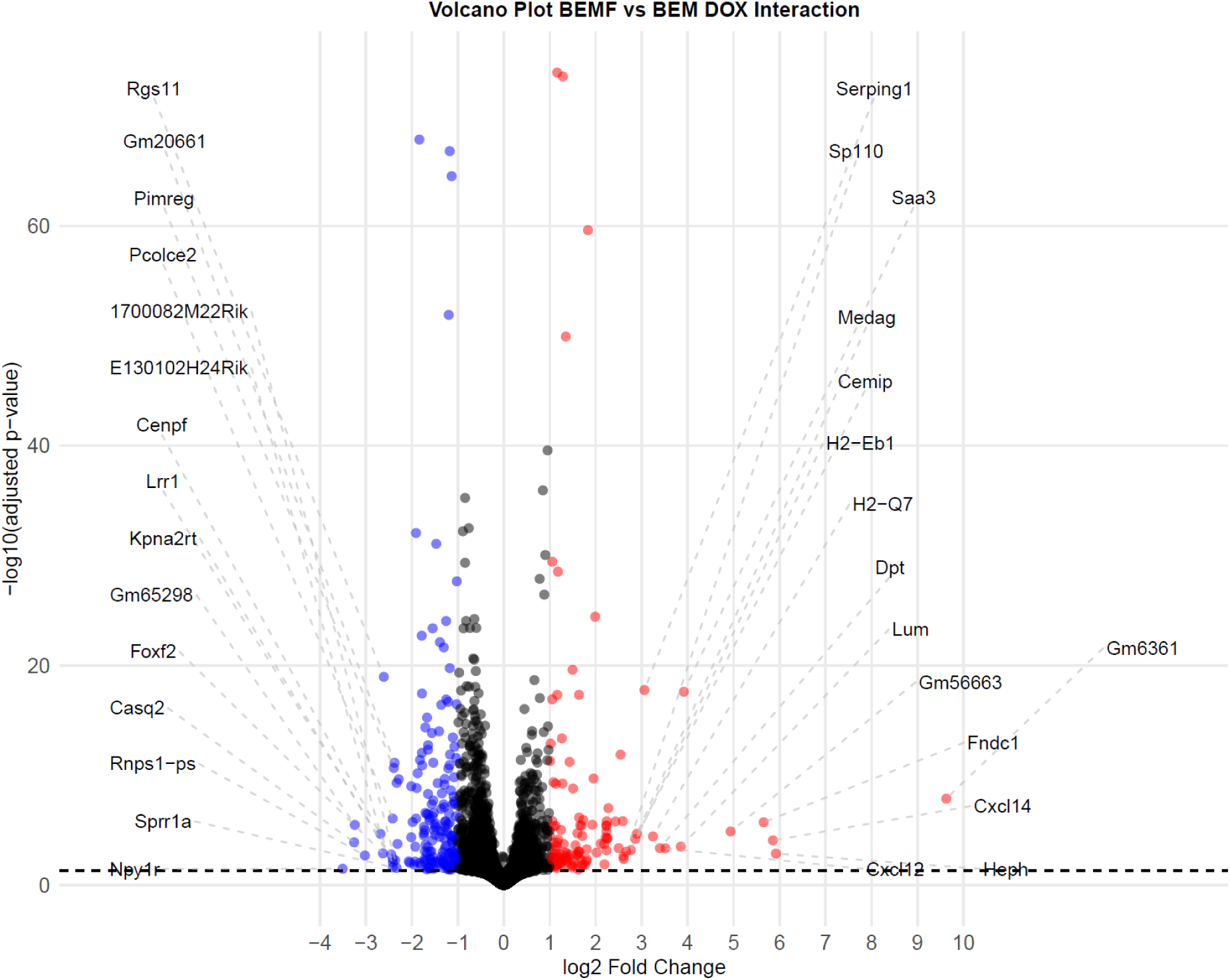
Volcano plot of differential expression of genes between *BEMF* and *BEM* spheroids following DOX treatment. Volcano plot shows differential expression of genes between *BEMF* and *BEM* spheroids in response to DOX. Each point represents one gene, plotted by log₂ fold change (x-axis) and –log₁₀(FDR-adjusted p-value) (y-axis). The horizontal dashed line marks the significance threshold (FDR < 0.05). Genes up-regulated in *BEMF* appear on the right, while down-regulated genes appear on the left. Points are coloured according to gene-level significance classification. The top 15 most strongly up-regulated and top 15 most strongly down-regulated genes (based on log₂ fold change among significant genes) are annotated with their MGI symbols. This visualization highlights the genes most strongly contributing to the differential DOX response between the two spheroid models.

Among these genes, 120 were upregulated and 179 were downregulated in *BEMF* compared to *BEM* spheroids, with at least a twofold change in expression (absolute log₂FC > 1), while the majority (1,874 genes) exhibited more modest effect sizes (absolute log₂FC ≤ 1). The top 50 genes ranked by statistical significance displayed distinct expression patterns associated with the response of spheroids containing fibroblasts to DOX administration (Table 4, Fig. 5).

**Figure 5.**
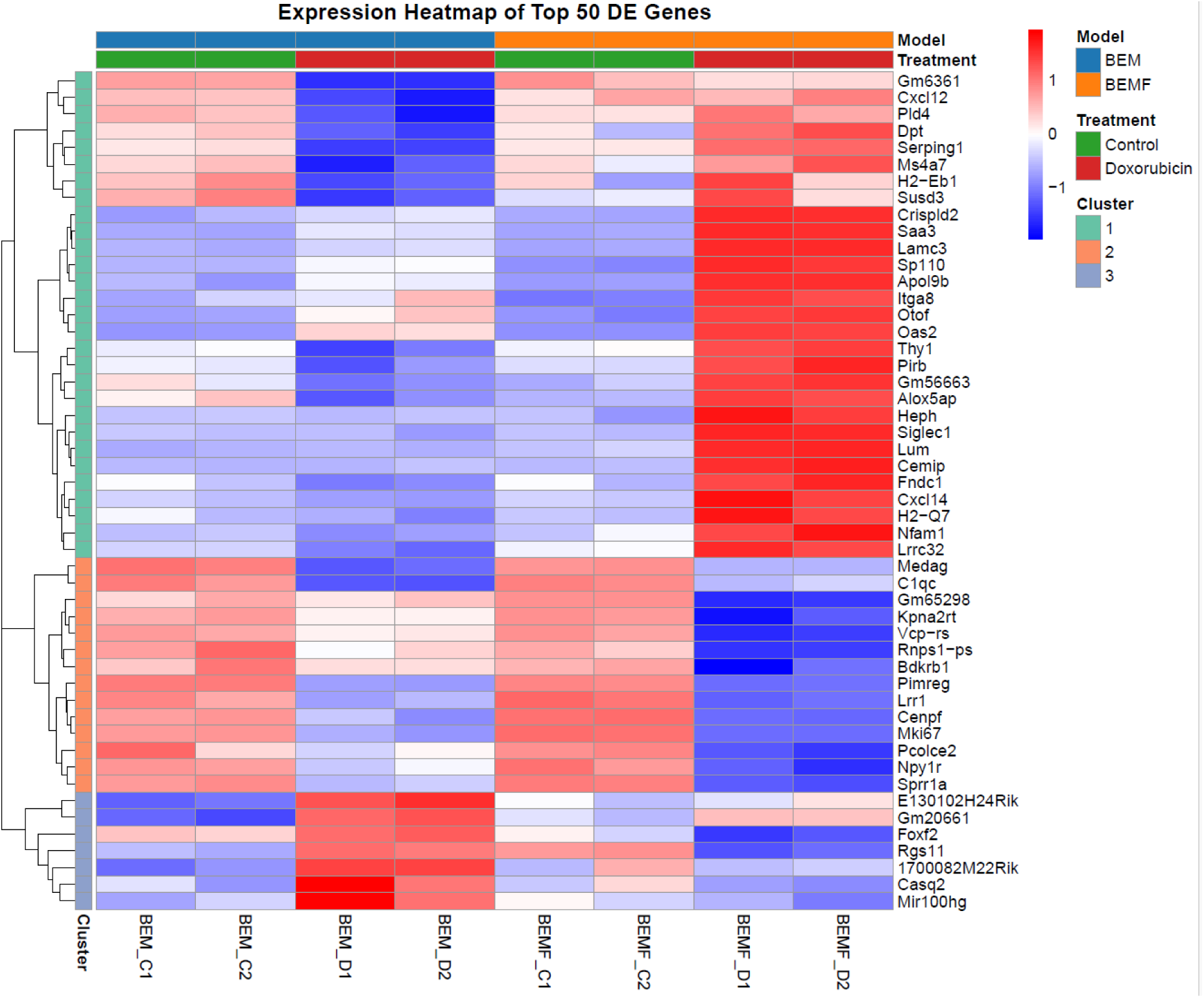
Expression heatmap of the top 50 differentially expressed between *BEMF* and *BEM* spheroids under DOX treatment. Rows represent the gene expression lever for the 50 genes with the largest absolute log₂ fold changes (FDR < 0.05), scaled by Z-score to emphasize relative expression patterns. Columns correspond to individual samples, annotated by model (*BEM* vs. *BEMF*) and treatment condition (control vs. doxorubicin). Hierarchical clustering was applied to rows using Euclidean distance and complete linkage, resulting in three gene clusters (Clusters 1–3). Warmer colours indicate higher relative expression, whereas cooler colours indicate lower relative expression. The heatmap highlights distinct expression patterns across models and treatment conditions.

**Table 4.** Top differentially expressed genes between *BEMF* and *BEM* spheroids under DOX treatment.

| Gene Symbol | log<sub>2</sub>FC | FDR-adjusted p-value | Biotype | Description | Cluster |
| --- | --- | --- | --- | --- | --- |
| Gm6361 | 9.63 | 1.4×10<sup>-8</sup> | Processed pseudogene | Predicted gene 6361 | 1 |
| Heph | 5.92 | 1.4×10<sup>-3</sup> | Protein coding | hephaestin | 1 |
| Cxcl14 | 5.85 | 8.6×10<sup>-5</sup> | Protein coding | C-X-C motif chemokine ligand 14 | 1 |
| Fndc1 | 5.65 | 1.9×10<sup>-6</sup> | Protein coding | fibronectin type III domain containing 1 | 1 |
| Gm56663 | 4.93 | 1.3×10<sup>-5</sup> | lncRNA | Predicted gene 56663 | 1 |
| Saa3 | 3.92 | 2.5×10<sup>-18</sup> | Protein coding | serum amyloid A 3 | 1 |
| Lum | 3.85 | 3.2×10<sup>-4</sup> | Protein coding | lumican | 1 |
| Cxcl12 | 3.52 | 4.6×10<sup>-4</sup> | Protein coding | C-X-C motif chemokine ligand 12 | 1 |
| Dpt | 3.4 | 4.4×10<sup>-4</sup> | Protein coding | dermatopontin | 1 |
| H2-Q7 | 3.24 | 3.8×10<sup>-5</sup> | Protein coding | histocompatibility 2, Q region locus 7 | 1 |
| Serping1 | 3.06 | 1.8×10<sup>-18</sup> | Protein coding | serine (or cysteine) peptidase inhibitor, clade G, member 1 | 1 |
| Cemip | 2.86 | 6.4×10<sup>-5</sup> | Protein coding | cell migration inducing protein, hyaluronan binding | 1 |
| H2-Eb1 | 2.76 | 6.6×10<sup>-4</sup> | Protein coding | histocompatibility 2, class II antigen E beta | 1 |
| Sp110 | 2.66 | 9.3×10<sup>-4</sup> | Protein coding | Sp110 nuclear body protein | 1 |
| Otof | 2.61 | 4.2×10<sup>-3</sup> | Protein coding | otoferlin | 1 |
| Susd3 | 2.6 | 2.5×10<sup>-3</sup> | Protein coding | sushi domain containing 3 | 1 |
| Pld4 | 2.59 | 1.5×10<sup>-6</sup> | Protein coding | phospholipase D family member 4 | 1 |
| Thy1 | 2.54 | 1.3×10<sup>-12</sup> | Protein coding | thymus cell antigen 1, theta | 1 |
| Crispld2 | 2.5 | 4.3×10<sup>-4</sup> | Protein coding | cysteine-rich secretory protein LCCL domain containing 2 | 1 |
| Siglec1 | 2.43 | 1.7×10<sup>-6</sup> | Protein coding | sialic acid binding Ig-like lectin 1, sialoadhesin | 1 |
| Oas2 | 2.27 | 1×10<sup>-8</sup> | Protein coding | 2'-5' oligoadenylate synthetase 2 | 1 |
| Alox5ap | 2.25 | 5.4×10<sup>-5</sup> | Protein coding | arachidonate 5-lipoxygenase activating protein | 1 |
| Ms4a7 | 2.24 | 6.3×10<sup>-5</sup> | Protein coding | membrane-spanning 4-domains, subfamily A, member 7 | 1 |
| Itga8 | 2.24 | 1.6×10<sup>-6</sup> | Protein coding | integrin alpha 8 | 1 |
| Pirb | 2.23 | 1.4×10<sup>-5</sup> | Protein coding | paired Ig-like receptor B | 1 |
| Apol9b | 2.23 | 3.6×10<sup>-6</sup> | Protein coding | apolipoprotein L 9b | 1 |
| Nfam1 | 2.23 | 7×10<sup>-5</sup> | Protein coding | Nfat activating molecule with ITAM motif 1 | 1 |
| Lrrc32 | 2.22 | 4.6×10<sup>-5</sup> | Protein coding | leucine rich repeat containing 32 | 1 |
| Lamc3 | 2.19 | 1.3×10<sup>-2</sup> | Protein coding | laminin gamma 3 | 1 |
| Npy1r | -3.51 | 3.3×10<sup>-2</sup> | Protein coding | neuropeptide Y receptor Y1 | 2 |
| Gm65298 | -3.24 | 3.4×10<sup>-6</sup> | lncRNA | Predicted gene 65298 | 2 |
| Rnps1-ps | -3.02 | 2×10<sup>-3</sup> | Processed pseudogene | RNA binding protein with serine rich domain 1, pseudogene | 2 |
| Medag | 2.89 | 2.1×10<sup>-5</sup> | Protein coding | mesenteric estrogen dependent adipogenesis | 2 |
| Cenpf | -2.45 | 1.7×10<sup>-3</sup> | Protein coding | centromere protein F | 2 |
| Pcolce2 | -2.42 | 8.9×10<sup>-7</sup> | Protein coding | procollagen C-endopeptidase enhancer 2 | 2 |
| Lrr1 | -2.42 | 4.2×10<sup>-3</sup> | Protein coding | leucine rich repeat protein 1 | 2 |
| Kpna2rt | -2.42 | 8.5×10<sup>-3</sup> | Protein coding | karyopherin subunit alpha 2, retrotransposed | 2 |
| Sprr1a | -2.41 | 2.4×10<sup>-2</sup> | Protein coding | small proline-rich protein 1A | 2 |
| Pimreg | -2.39 | 2.2×10<sup>-11</sup> | Protein coding | PICALM interacting mitotic regulator | 2 |
| Bdkrb1 | -2.33 | 2.7×10<sup>-2</sup> | Protein coding | bradykinin receptor, beta 1 | 2 |
| Vcp-rs | -2.33 | 5×10<sup>-10</sup> | Processed pseudogene | valosin containing protein, related sequence | 2 |
| Mki67 | -2.29 | 2.3×10<sup>-10</sup> | Protein coding | antigen identified by monoclonal antibody Ki 67 | 2 |
| C1qc | 2.23 | 4.2×10<sup>-6</sup> | Protein coding | complement component 1, q subcomponent, C chain | 2 |
| Casq2 | -3.26 | $1.3 \times 10^{-4}$ | Protein coding | calsequestrin 2 | 3 |
| E130102H24Rik | -2.68 | $2.2 \times 10^{-5}$ | lncRNA | RIKEN cDNA E130102H24 gene | 3 |
| Foxf2 | -2.63 | $1.3 \times 10^{-13}$ | Protein coding | forkhead box F2 | 3 |
| Rgs11 | -2.61 | $1.1 \times 10^{-19}$ | Protein coding | regulator of G-protein signaling 11 | 3 |
| Gm20661 | -2.38 | $7.4 \times 10^{-12}$ | Protein coding | Predicted gene 20661 | 3 |
| 1700082M22Rik | -2.36 | $5.5 \times 10^{-3}$ | lncRNA | RIKEN cDNA 1700082M22 gene | 3 |
| Mir100hg | -2.31 | $1.8 \times 10^{-4}$ | lncRNA | Mir100 Mirlet7a-2 Mir125b-1 cluster host gene | 3 |

Hierarchical clustering of the top 50 interaction-associated genes identified three distinct expression clusters. **Cluster 1** comprises genes that show higher expression in *BEMF* spheroids when exposed to DOX, while showing relatively low or baseline expression in the other clusters. **Cluster 2** includes genes that exhibit lower expression levels in *BEMF* spheroids receiving DOX. **Cluster 3** consists of genes showing higher expression levels in *BEM* spheroids upon DOX administration, with lower expression levels across both control and *DOX-BEMF* spheroids (BEMF spheroids exposed to DOX) (Fig. 5).

To characterize the main function of the expression clusters, genes within each cluster were annotated based on their membership in Hallmark Gene Sets from MSigDB (Fig. 6). Genes upregulated in *DOX-BEMF* spheroids in comparison with *DOX-BEM* spheroids (BEM spheroids exposed to DOX) within **Cluster 1** were predominantly assigned to immune- and inflammation-related Hallmark categories, including *Allograft Rejection*, *Complement*, *Interferon-α Response, Interferon-γ Response, Inflammatory Response*, and *Coagulation*, as well as stress-associated categories such as *Apoptosis* and *Epithelial–Mesenchymal Transition*. Genes downregulated in *DOX-BEMF* spheroids within **Cluster 2** were frequently assigned to cell-cycle–related Hallmark categories, including *E2F Targets, G2M Checkpoint*, and *Mitotic Spindle*, along with hormone-responsive categories (*Estrogen Response Early* and *Late*) and developmental signalling pathways such as *Hedgehog Signalling*. Genes in **Cluster 3**, which were induced in *DOX-BEM* spheroids and reduced in *DOX-BEMF* spheroids, were primarily assigned to *KRAS Signalling Down* and *Myogenesis*.

**Figure 6.**
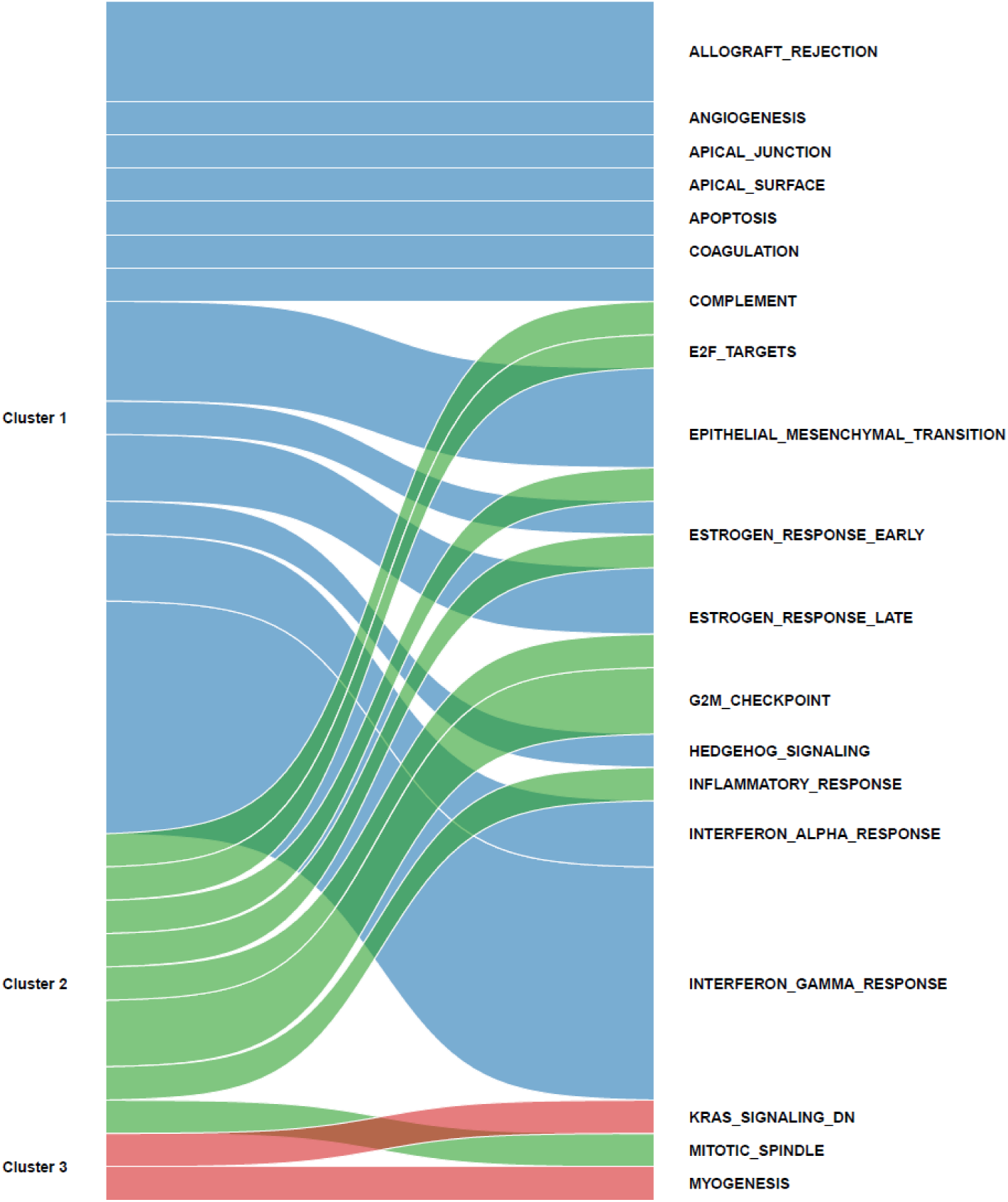
Functional programs linked to distinct clusters of fibroblast-dependent DOX responses. Sankey diagram illustrates the correspondence between interaction-derived gene clusters and their membership in MSigDB Hallmark gene sets. Each cluster (left) represents genes grouped by hierarchical clustering of expression profiles associated with the cell composition and treatment interaction. Cluster 1 comprises genes upregulated by doxorubicin in *BEMF* spheroids, Cluster 2 includes genes more strongly downregulated in doxorubicin-treated *BEMF* spheroids, and Cluster 3 contains genes preferentially induced in *BEM* spheroids. Ribbons connect each cluster to Hallmark pathways (right) to which genes within the cluster belong. Ribbon thickness reflects the number of genes in each cluster annotated to a given Hallmark pathway, and ribbon colour denotes cluster identity. The diagram summarizes the distribution of Hallmark pathway memberships across clusters for genes exhibiting fibroblast-dependent DOX responses.

While cluster-based analysis highlighted distinct gene expression patterns associated with fibroblast inclusion in melanoma spheroids under DOX treatment, these findings suggested coordinated shifts in underlying biological programs, thus prompting a global pathway-level assessment using Gene Set Enrichment Analysis (GSEA).

#### 2.3.1. Hallmark GSEA defines the primary transcriptional programs underlying fibroblast-modulated response to DOX in melanoma spheroids

GSEA of the MSigDB Hallmark collection identified distinct patterns of pathway regulation associated with fibroblast effect on DOX response of melanoma spheroids (Fig. 7; Table S2). The most significantly suppressed pathways in *DOX-BEMF* spheroids (NES < 0) included core proliferative programs, such as *G2/M Checkpoint* (NES = −2.38, q = 1.3 × 10⁻²¹), *E2F Targets* (NES = −2.25, q = 9.9 × 10⁻¹⁶), *Mitotic Spindle* (NES = −2.13, q = 5.8 × 10⁻¹²), and *MYC Targets V2* (NES = −1.88, q = 6.2 × 10⁻⁴), as well as *Unfolded Protein Response* and *MYC Targets V1* (NES ≈ −1.77 and −1.60, respectively; both q ≈ 1.1 × 10⁻³). Together, these negatively enriched signatures correspond to pathways involved in DNA replication, mitotic progression, and cell-cycle regulation.

**Figure 7.**
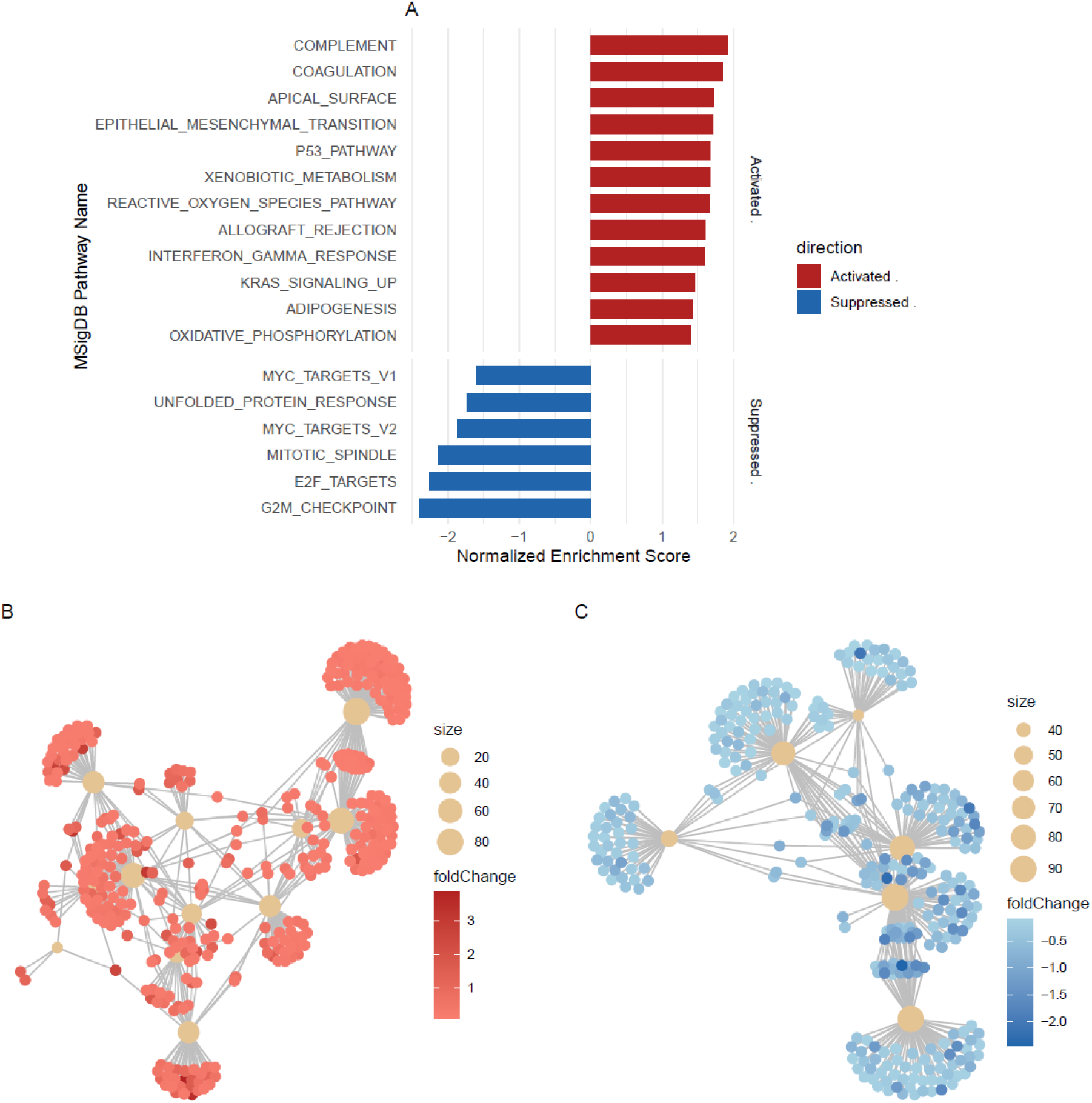
Gene set enrichment and network representation of DOX-responsive pathways in *BEMF* versus *BEM* spheroids. **(A)** Gene Set Enrichment Analysis (GSEA) results for MSigDB Hallmark pathways following DOX treatment in *BEMF* versus *BEM* spheroids Bar plots display the top positively and negatively enriched pathways ranked by normalized enrichment score (NES). Red bars indicate positively enriched pathways, and blue bars indicate negatively enriched pathways. **(B)** CNET plot of positively enriched Hallmark pathways. Nodes represent genes, coloured by log₂ fold change, while beige nodes represent Hallmark pathways and are sized according to the number of genes in the pathway. Edges connect pathways to their leading-edge genes. **(C)** CNET plot of negatively enriched Hallmark pathways. Genes and pathways are displayed as in panel B, highlighting network structure among negatively enriched pathways.

Conversely, several immune, inflammatory, and stress-adaptation pathways were strongly activated in *DOX-BEMF* (NES > 0), including *Complement* (NES = 1.87, q = 6.2 × 10⁻⁴), *Coagulation*, *Interferon Gamma Response*, *Allograft Rejection*, and *Xenobiotic Metabolism* (all NES ≈ 1.57–1.81, q ≤ 8.3 × 10⁻³), alongside epithelial and metabolic programs such as *Epithelial–Mesenchymal Transition*, *Apical Surface*, *Adipogenesis*, and *Oxidative Phosphorylation* (NES ≈ 1.39–1.70, q ≈ 2.6 × 10⁻²). These positively enriched signatures correspond to immune- and inflammation-associated pathways, epithelial/mesenchymal-related programs, and metabolic processes in fibroblast-containing spheroids under DOX exposure.

#### 2.3.2. Reactome analysis revealed several recurring functional themes associated with fibroblast effects on the response of melanoma spheroids to DOX

To examine the higher-order organisation of pathway-level responses to DOX in BEMF compared to BEM spheroids, and to reduce the redundancy inherent in the large set of enriched Reactome pathways (Table S3), semantically similar pathways were grouped into representative clusters using a semantic similarity–based approach (Table 5). Clustering of significantly enriched pathways (FDR < 0.05) identified eight distinct pathway modules showing significant differences in enrichment between the two types of spheroids, when treated with DOX. Visualization of these pathways in a low-dimensional embedding space revealed clear separation between clusters and consistent grouping of related pathways (Fig. 8A). Across clusters, pathways related to cell-cycle progression and mitotic regulation consistently exhibited negative normalized enrichment scores in *DOX-BEMF* compared to *DOX-BEM* spheroids. These included pathways involved in G2/M progression, mitotic spindle organization, chromosome segregation, and cell-cycle checkpoint regulation. In the *BEMF* spheroids receiving DOX, two clusters were enriched for such pathways, encompassing spindle-assembly checkpoint components (e.g. MAD2 signalling, PLK1 regulation), APC/C-mediated anaphase control, ATR/ATM-mediated DNA damage response modules, and core mitotic transition pathways (Fig. 8B). Negative enrichment of these pathways was more pronounced in the *DOX-BEMF* spheroids compared with *DOX-BEM* spheroids.

**Figure 8.**
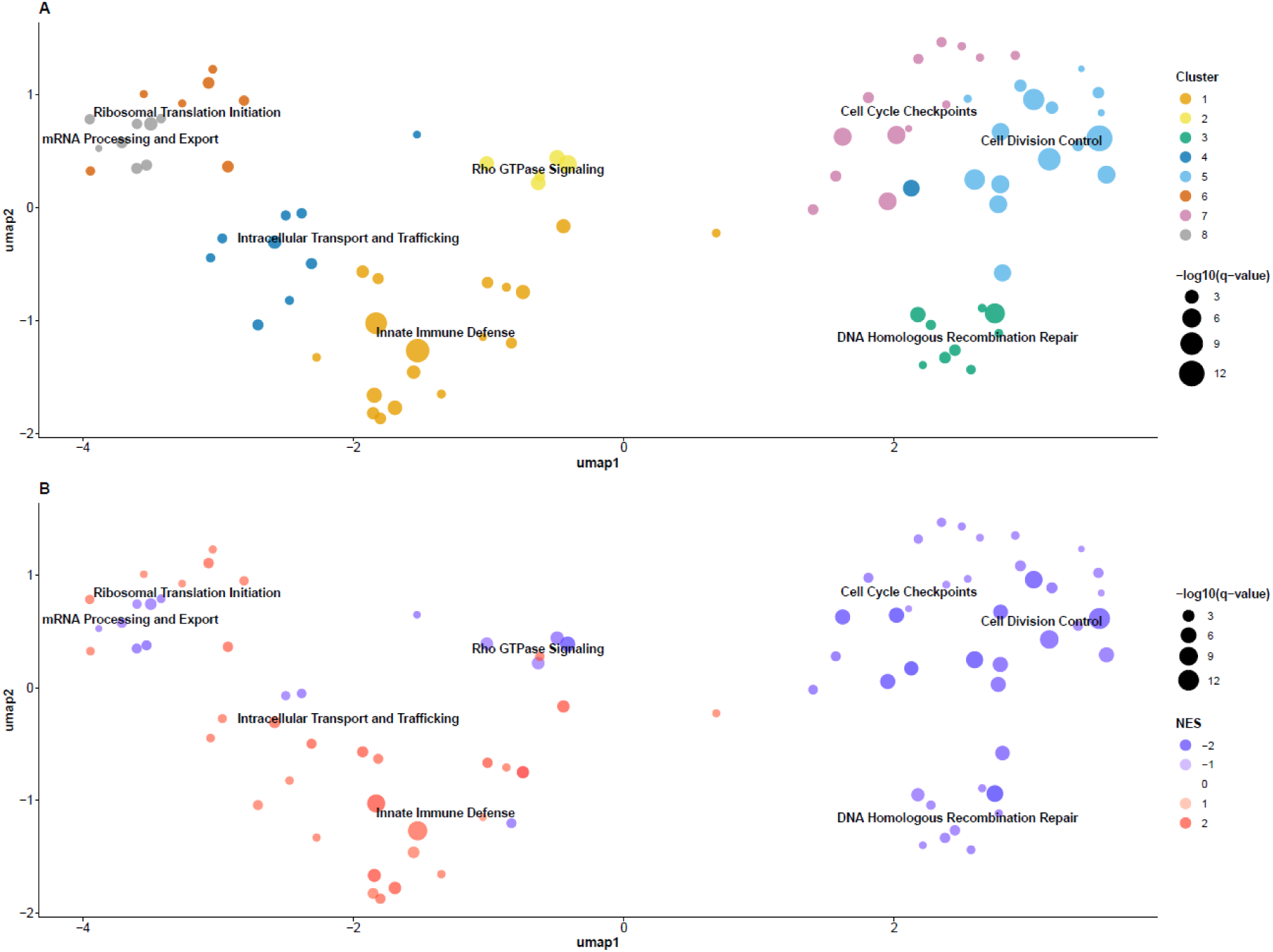
UMAP-based clustering of enriched Reactome pathways with LLM-derived biological labels. **(A)** UMAP projection of enriched Reactome pathways based on semantic similarity of pathway descriptions and k-means clustering (k = 8). Each point represents a pathway and is coloured according to its assigned cluster. Point size reflects pathway significance (−log₁₀FDR). Cluster labels were generated by LLM–based semantic summarization of pathway names within each cluster and are displayed at cluster centroids. **(B)** The same UMAP embedding as in panel A, with pathways coloured according to normalized enrichment score (NES) to indicate the direction and magnitude of enrichment. Warmer colours represent positive NES values, whereas cooler colours represent negative NES values. Point size corresponds to −log₁₀FDR.

**Table 5.**
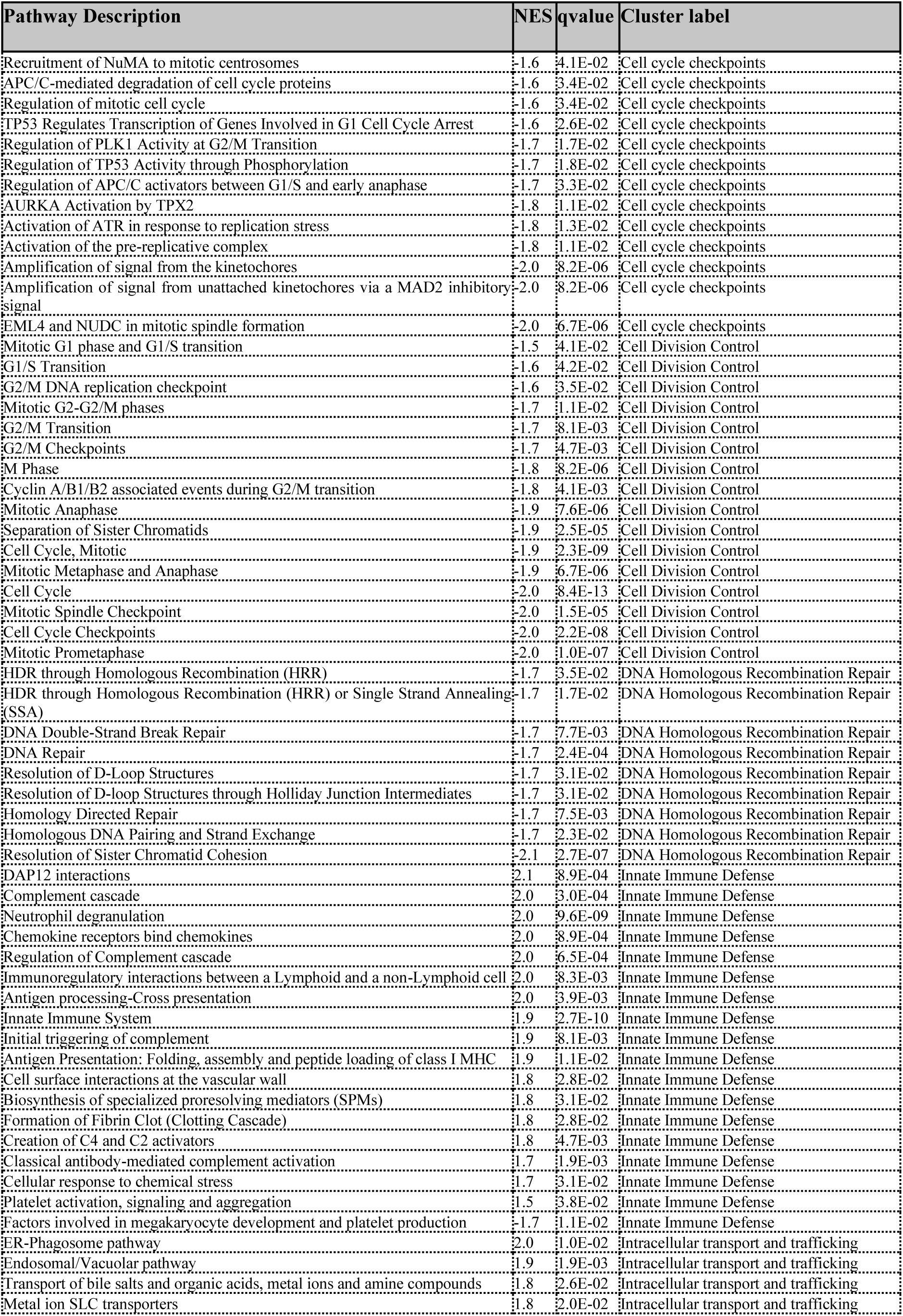
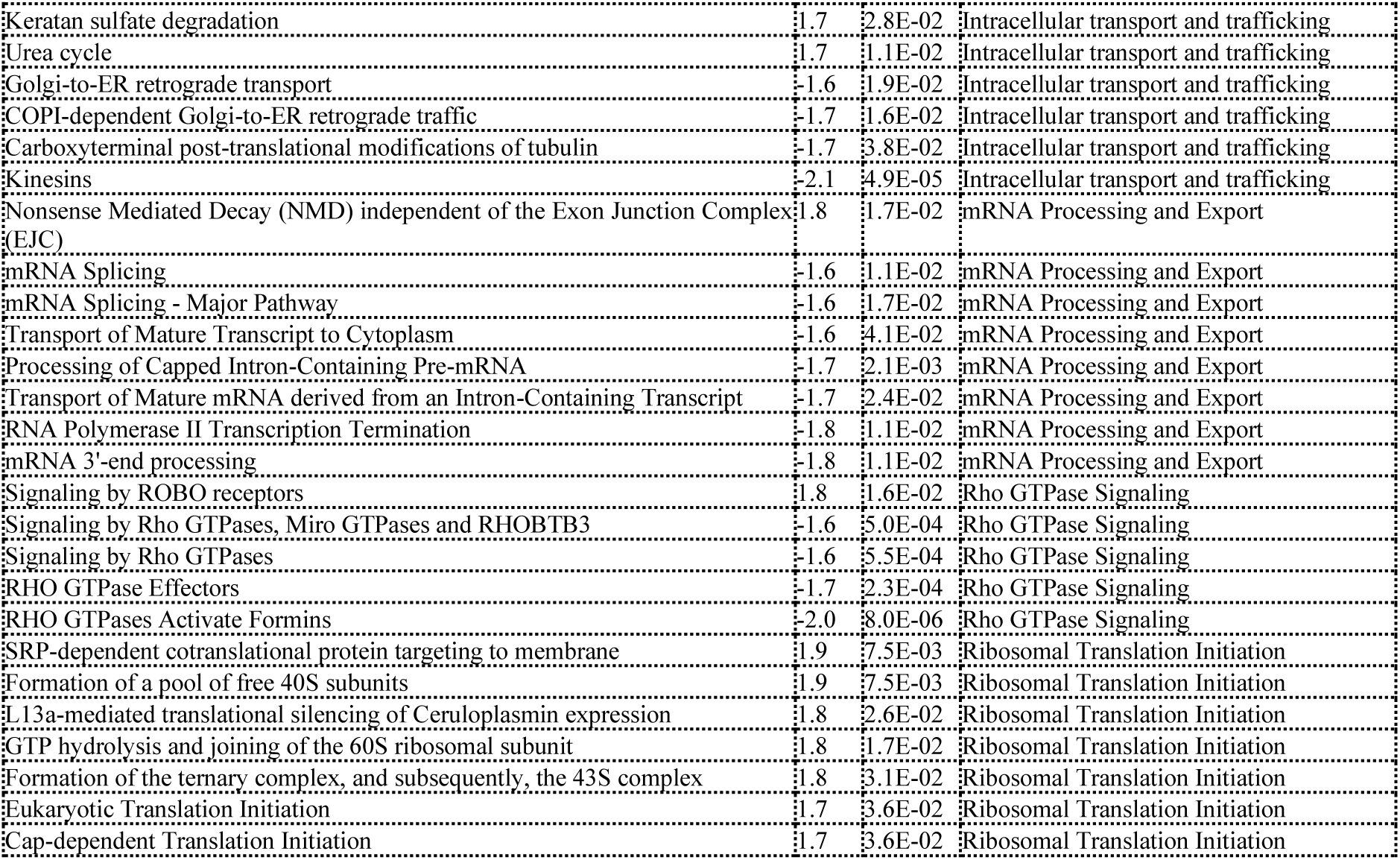
Top Reactome pathways activated or suppressed by DOX treatment.

Pathways associated with homologous recombination repair exhibited suppression in both models, with stronger negative effect observed in treated BEMF spheroids. This pattern was consistent with the downregulation of cell-cycle– and checkpoint-related pathways identified in the same analysis.

A distinct semantic cluster comprising multiple Rho GTPase signalling pathways exhibited negative normalized enrichment scores, indicating coordinated downregulation of these pathways in response to DOX in *BEMF* spheroids. Pathways within this cluster included several modules involved in Rho GTPase–mediated signalling. In contrast, ROBO signalling pathways showed modest transcriptomic positive enrichment in *BEMF* compared to *BEM* spheroids (Fig. 8B).

A second set of semantic clusters was enriched for pathways related to RNA processing and protein synthesis. Pathways involved in mRNA splicing, 3′-end formation, transcription termination, and nuclear export exhibited negative normalized enrichment scores under DOX exposure in *BEMF* compared to *BEM* spheroids. In contrast, in the same group, pathways associated with ribosomal translation initiation showed positive enrichment and the exon junction–independent branch of nonsense-mediated decay was the only RNA-processing–related pathway exhibiting positive enrichment within this module.

One cluster comprised pathways related to immune and vascular signalling. This cluster included DAP12 adaptor signalling, neutrophil degranulation, chemokine receptor signalling, multiple complement cascade pathways, and antigen processing and cross-presentation, all exhibiting positive normalized enrichment scores in *DOX-BEMF* spheroids versus *DOX-BEM* spheroids. Pathways associated with the clotting cascade, platelet activation, and cell–vessel wall interactions were also positively enriched, along with pathways involved in the biosynthesis of specialized pro-resolving mediators (SPMs), suggesting a microenvironment associated not only with inflammatory activation but also with induction of pro-resolving programs linked to tissue remodelling [27].

Another cluster comprised pathways related to intracellular transport and trafficking, which are positively enriched in the *DOX-BEMF* spheroids. Withing the same cluster, pathways associated with kinesin motor activity, carboxyterminal post-translational modifications of tubulin, and COPI-dependent Golgi-to-ER retrograde transport exhibited negative normalized enrichment scores in response to DOX treatment in *BEMF*, compared with *BEM* spheroids.

### 2.4. Transcription factor regulatory network analysis

RNA-Seq based analysis of transcription factor (TF) activity inferred widespread DOX induced changes across both spheroid types. For a substantial subset of TFs, however, the magnitude of the DOX response differed between spheroids. Using a linear modelling framework, we quantified a microenvironment-dependent DOX effect, defined as the difference in the DOX-induced change in TF activity between *BEMF* and *BEM* spheroids (ΔDOX_BEMF−BEM_) (Table 6).

**Table 6.** Difference in DOX-induced change in inferred TF activity between *BEMF* and *BEM* spheroids.

| Transcription factor | TF activity difference between DOX-treated *BEMF* and *BEM* spheroids | FDR-adjusted p-value |
| --- | --- | --- |
| Mybl2 | -0.94 | 3.88E-03 |
| Gfi1b | -0.74 | 9.50E-03 |
| Atf4 | -0.72 | 7.27E-03 |
| Zfp384 | -0.67 | 1.87E-02 |
| E2f4 | -0.65 | 1.03E-02 |
| Arid2 | -0.64 | 2.36E-02 |
| Mafb | -0.62 | 3.74E-02 |
| E2f1 | -0.61 | 7.27E-03 |
| Nrf1 | -0.60 | 9.03E-03 |
| Atf6 | -0.59 | 2.45E-03 |
| Pou5f1 | -0.56 | 2.89E-02 |
| Gabpa | -0.55 | 7.27E-03 |
| Ncoa3 | -0.55 | 2.66E-02 |
| Irf9 | -0.54 | 9.08E-03 |
| Nanog | -0.54 | 1.11E-02 |
| Zfx | -0.52 | 3.25E-02 |
| Gli2 | -0.52 | 2.26E-03 |
| Atf7 | -0.51 | 4.83E-02 |
| Trp53 | 0.55 | 7.49E-03 |
| Fos | 0.55 | 2.45E-03 |
| Lyl1 | 0.58 | 9.03E-03 |
| Usf2 | 0.64 | 2.26E-03 |
| Jun | 0.67 | 2.26E-03 |
| Nfkb1 | 0.67 | 6.62E-04 |
| Ets1 | 0.73 | 4.17E-03 |
| Ets2 | 0.86 | 2.45E-03 |
| Spi1 | 0.89 | 2.45E-03 |
| Erg | 1.16 | 4.74E-03 |

Several TFs exhibited significantly larger DOX-associated activity changes in *DOX-BEMF* compared with *DOX-BEM* spheroids (FDR < 0.05), including *Erg* (ΔDOX_BEMF−BEM_ = 1.16), *Spi1* (0.89), *Ets2* (0.86), *Ets1* (0.73), *Nfkb1* (0.67), *Jun* (0.67), and *Usf2* (0.64). Additional TFs showing significant but more moderate *BEMF*-biased DOX effects included *Trp53, Fos, Rela, Smad4, Foxo3,* and *Stat3*. Collectively, these TFs are consistent with enhanced regulation of inflammatory and stress-response signalling (e.g., *Nfkb1/Rela, Jun/Fos, Stat3, Spi1*) [28–31], DNA-damage and apoptosis-related programs (e.g., *Trp53, Foxo3*) [32,33], and microenvironment- and vascular/remodelling-associated processes (e.g., *Erg, Ets1/Ets2, Smad4*), including cytokine signalling and extracellular matrix regulation [29,31,34–36].

Conversely, a distinct set of TFs showed DOX-associated activity changes that were attenuated in *DOX-BEMF* relative to *DOX-BEM* spheroids. These included *Mybl2* (−0.94), *Atf4* (−0.72), *E2f4* (−0.65), *E2f1* (−0.61), *Atf6* (−0.59), *Gabpa* (−0.55), *Irf9* (−0.54), and *Gli2* (−0.52) (FDR < 0.05). Collectively, these TFs are linked to cell-cycle and proliferation-associated programs (e.g., *Mybl2, E2f1,* and *E2f4*) [37], as well as intrinsic stress-response pathways, including the unfolded protein response (e.g., *Atf4* and *Atf6*) [38], interferon signalling (e.g., *Irf9*) [39], and developmental signalling processes (e.g., *Gli2*) [40].

In summary, these analyses indicate that fibroblast presence selectively amplifies DOX-induced activation of inflammatory and stress-response TF networks while attenuating proliferation-linked regulatory programs.

### 2.5. Literature-derived resistance gene behaviour and cross-study validation

#### 2.5.1 Resistance signature construction and overlap

To construct a melanoma-relevant DOX resistance gene set, we combined literature-derived annotations with pathway-based information from MSigDB. A total of 48 mouse genes were identified from NCBI as referencing DOX resistance in melanoma and were mapped successfully to *Mus musculus* orthologs. Integration with DOX-associated MSigDB gene sets expanded this list to 78 genes. A parallel analysis of multidrug resistance in melanoma yielded 129 mouse genes. Comparison of the extended and multidrug resistance sets revealed 16 shared genes between them (Fig. 9A).

**Figure 9.**
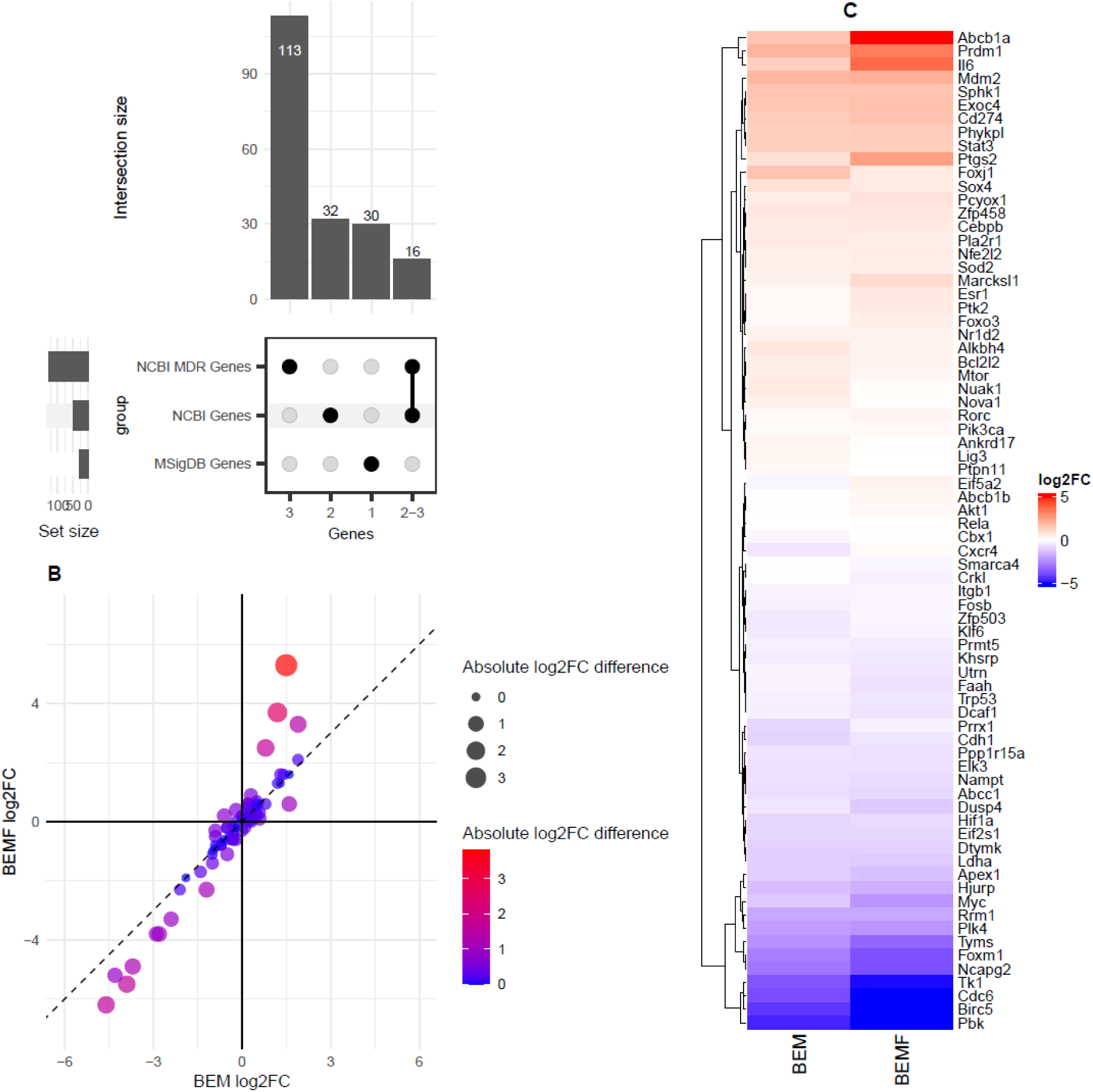
Comparative analysis of curated DOX-resistance genes. **(A)** UpSet plot summarizing the overlap between three resistance gene sets: NCBI-annotated doxorubicin resistance genes (NCBI Genes), MSigDB-derived doxorubicin-related genes (MSigDB Genes), and NCBI multidrug resistance (MDR) genes (NCBI MDR Genes). Bars indicate the size of each exclusive intersection, with filled circles denoting membership in the corresponding gene sets. **(B)** Scatter plot comparing log₂ fold changes (DOX vs control) for genes in the extended DOX-resistance gene list between the *BEM* model (x-axis) and the *BEMF* spheroids (y-axis). Point colour and size represent the absolute difference in log₂ fold change between models. **(C)** Heatmap of log₂ fold changes for the extended DOX-resistance gene set in *BEM* and *BEMF* spheroids. Genes were hierarchically clustered based on their response profiles. Colours represent log₂ fold change values.

#### 2.5.2 Resistance-signature response differences between BEM and BEMF

To assess whether fibroblast inclusion within spheroids altered the transcriptional response of curated DOX resistance–associated genes, DOX-induced log₂ fold changes were compared between the *BEM* and *BEMF* models (Fig. 9A-C). Scatter-based comparison of the extended resistance gene set showed a systematic shift toward larger absolute responses in the *BEMF* condition (Fig. 9B). While many genes exhibited concordant directionality between DOX-treated spheroids, a subset (*Abcb1a, Cdc6, Foxm1, Il6, Myc, Pbk, Prdm1, Ptgs2, Tk1)* displayed pronounced differences in response magnitude. Focusing on the genes in the constructed resistance signature, particularly on those which displayed an absolute difference in treatment response of at least one log₂ fold change between the two spheroid models, nine resistance-associated genes stand out as exhibiting a substantially stronger effect in *DOX-BEMF* compared with *DOX-BEM* spheroids (Fig. 9C). These included *Abcb1a, Il6, Ptgs2* and *Prdm1*, all of which showed increased upregulation in the fibroblast-containing spheroids compared to fibroblast-free spheroids. In contrast, several genes associated with cell-cycle and proliferation processes, including *Pbk, Cdc6, Tk1, Myc*, and *Foxm1*, displayed stronger downregulation in *DOX*-*BEMF* relative to *DOX*-*BEM* spheroids.

#### 2.5.3. Directionality meta-analysis across six conditions

From the studies selected for comparison, the following datasets were included: *GSE33624* (in this dataset, MelJuSo human cells grown as monolayer, were exposed to doxorubicin for 2 h, after which the drug was removed and samples were collected 1 and 2 days later.), *GSE212112* (in this dataset, B16.F10 mouse cells grown as a monolayer, were exposed to 0.1 µM DOX for 7 days, followed by sampling), and *GSE246690* (in this dataset, SK-MEL-147 human cells grown as a monolayer, were exposed to0.1 µM DOX for 3 days, after which the medium was replaced with drug-free medium and samples were collected 7 days after treatment initiation). These datasets encompass both human and murine melanoma models and represent distinct DOX exposure paradigms, ranging from acute high-dose treatment to prolonged low-dose exposure and recovery-based adaptive response models. Although these models differ in species, cell line, type of culture, DOX dose, and the duration of DOX exposure, a directionality-based meta-analysis was used to test whether a common transcriptional response to DOX could be detected across heterogeneous experimental contexts. This approach prioritizes consistency in the sign of regulation rather than exact agreement in effect size, making it well suited for identifying conserved response programs across biologically and technically distinct systems.

Directionality-based meta-analysis across six conditions (*DOX*-*BEM* vs *BEM* control, *DOX-BEMF* vs *BEMF* control, *GSE33624* day 1 treated vs control, *GSE33624* day 2 treated vs control, *GSE212112*, and *GSE246690*) identified a subset of genes exhibiting statistically significant and highly consistent regulation (Table 7). Among the top-ranked genes, *Mki67, Birc5, Cenpf, Pbk, Cdc6, Tk1,* and *Tyms* showed strong and concordant down-regulation across datasets, with negative aggregated signed scores ranging from −3.5 to −5.0 and FDR-adjusted q-values below 0.001. These genes displayed near-complete directional concordance, with negative effect directions observed in five or six of the six conditions and mean absolute log₂ fold-change values exceeding 2 in most cases. Together, these findings indicate the presence of a conserved DOX-associated suppression program enriched for genes involved in proliferation, cell-cycle progression, and DNA replication.

**Table 7.** Cross-model directionality meta-analysis identifies conserved transcriptional responses to DOX.

| **Gene** | **Number of studies where the gene is identified** | **Number of studies where the gene is upregulated** | **Number of studies where the gene is downregulated** | **Directionality classification** |
| --- | --- | --- | --- | --- |
| Mki67 | 6 | 0 | 6 | Consistently Down |
| Birc5 | 6 | 0 | 6 | Consistently Down |
| Cenpf | 6 | 0 | 6 | Consistently Down |
| Pbk | 6 | 0 | 6 | Consistently Down |
| Cdc6 | 6 | 1 | 5 | Consistently Down |
| Ncapg2 | 4 | 0 | 4 | Consistently Down |
| Tfap4 | 6 | 0 | 6 | Consistently Down |
| Tk1 | 6 | 1 | 5 | Consistently Down |
| Tyms | 6 | 0 | 6 | Consistently Down |
| Il6 | 6 | 6 | 0 | Consistently Up |
| Myc | 6 | 0 | 6 | Consistently Down |
| Foxm1 | 6 | 2 | 4 | Consistently Down |
| Pdss1 | 6 | 0 | 6 | Consistently Down |
| Medag | 2 | 0 | 2 | Consistently Down |
| Etv4 | 6 | 2 | 4 | Consistently Down |
| Lrr1 | 4 | 1 | 3 | Mixed / Inconsistent |
| Lum | 5 | 4 | 1 | Mixed / Inconsistent |
| Ptgs2 | 4 | 3 | 1 | Mixed / Inconsistent |
| Topbp1 | 6 | 1 | 5 | Mixed / Inconsistent |
| Siva1 | 4 | 1 | 3 | Mixed / Inconsistent |
| Prdm1 | 5 | 4 | 1 | Mixed / Inconsistent |
| Abcb1a | 6 | 3 | 3 | Mixed / Inconsistent |
| Cxcl12 | 4 | 3 | 1 | Mixed / Inconsistent |
| Slc29a2 | 4 | 2 | 2 | Mixed / Inconsistent |

Additional genes, including *Ncapg2, Tfap4, Myc, Foxm1, Pdss1, Medag*, and *Etv4*, also exhibited significant negative aggregated signed scores (q < 0.05), although with reduced directional concordance and greater variability in effect magnitude across conditions. In contrast, *Il6* was the only gene demonstrating statistically significant and consistent up-regulation, with a positive aggregated signed score (E = 3.40, q = 0.0016) and complete directional concordance across all six conditions. The remaining genes did not reach statistical significance after multiple-testing correction and were classified as directionally inconsistent, despite exhibiting directional biases in individual datasets.

#### 2.5.4 Cross-study overlap intersections and *BEMF* spheroid specific shared genes

To identify conserved DOX response programmes associated with fibroblast-containing spheroids, we extracted lists of under-expressed genes (FDR < 0.05 and log₂ fold change < −1) and over-expressed genes (FDR < 0.05 and log₂ fold change > 1) from the differential expression analyses of *DOX BEM* and *DOX-BEMF* spheroids versus their respective controls, as well as from two of the public RNA-Seq datasets described above (*GSE246690* and *GSE212112*; DOX-treated versus control). Across all four conditions, 392 genes were significantly and consistently differentially expressed, of which 293 were downregulated and 99 were upregulated. Intersecting this shared gene set with the curated resistance gene list identified 10 downregulated resistance-associated genes: *Birc5, Cdc6, Foxm1, Myc, Plk4, Ncapg2, Rrm1, Tk1, Pbk,* and *Tyms*. Five of these genes (*Cdc6, Foxm1, Myc, Tk1,* and *Pbk*) were also among those showing the largest DOX-induced response differences between *BEM* and *BEMF* spheroids.

Focusing on genes that were significantly differentially expressed in *DOX-BEMF* spheroids but not in *DOX-BEM* spheroids, and that were shared with the other studies, we identified a set of 20 genes. Among these, only a single gene (*Siva1*) was upregulated, whereas 19 genes were downregulated: *Etv1, Ddx11, Dazap1, Tfap4, Hnrnpm, Cep128, Srsf7, Hnrnpa2b1, Cse1l, Exosc2, Dkc1, Mdc1, Ncapd3, Srsf2, Topbp1, Tm4sf1, Etv4, Ifrd2*, and *Rbm14*. None of these genes overlapped with the curated DOX resistance gene signature derived from NCBI annotations.

## 3. Discussion

Non-mutational drug resistance plays a critical role in protecting cancer cells from lethal drug exposure [41] and is increasingly recognized as being shaped by the TME [42]. TME-derived factors can mediate drug resistance, including stromal growth factors such as HGF and TGF-β, inflammatory cytokines and chemokines such as IL-6, IL-8, and CXCL12, extracellular matrix- and integrin-mediated signals, exosome-mediated transfer of resistance-promoting cargo, and hypoxia-associated metabolic adaptation [43,44]. Accordingly, anticancer agents effective against tumour cells cultured alone often lose efficacy in the presence of stromal cells [45–47]. Cancer associated fibroblasts (CAFs) are known to be involved in remodelling the extracellular matrix to enhance tumour stiffness and restrict drug penetration, secreting paracrine factors such as IL-6, and TGF-β that activate pro-survival and stemness-associated signalling pathways in cancer cells, activation of drug efflux pathways, and modulating immunosuppressive stromal crosstalk [10,15,48]. Collectively, by coordinating these interconnected processes, CAFs favour a tumour milieu that supports resistance towards various anticancer strategies including chemo-/immunotherapy, radiotherapy, targeted therapy, anti-angiogenic therapy, as well as endocrine therapy [49].

In line with these findings, incorporation of fibroblasts into melanoma spheroids containing macrophages and endothelial cells was associated with an increased DOX concentration required to achieve growth inhibition. *BEMF* spheroids exhibited higher IC₃₀ and IC₅₀ values and maintained greater viability across a range of DOX concentrations (0.156–10 μM) compared with *BEM* spheroids (Fig. 1). Comparison of the fitted dose–response curves did not reveal a statistically significant difference between conditions, though area under the curve (AUC) analysis across the full dose range showed higher overall viability in *DOX-BEMF* compared to *DOX-BEM.* In untreated spheroids, fibroblast inclusion was associated with modest transcriptional changes (Fig. 2), suggesting the involvement of these cells in the establishment of a primed state rather than large-scale transcriptional rewiring, that could be favour drug tolerance by shaping subsequent stress responses [41,50].

Upon DOX exposure, fibroblast-containing spheroids exhibited a broader and more pronounced transcriptional response than fibroblast-free spheroids, as reflected in both the number and magnitude of differentially expressed genes. Notably, nearly 300 genes displayed a statistically significant increase or decrease in effect size of at least twofold in *DOX*-*BEMF* spheroids compared with *DOX-BEM* (Fig. 4), consistent with coordinated biological regulation.

Fibroblasts reshaped drug-induced responses by driving a shift away from proliferative programs toward inflammatory, immune-associated, and stress-adaptive transcriptional states. This shift was evident at both the gene (Fig. 5) and pathway levels, manifested as coordinated suppression of cell-cycle and cell-division programmes, including attenuation of E2F- and MYC-associated transcriptional networks and core mitotic processes such as G2/M checkpoint regulation, mitotic spindle assembly, chromosome segregation, and DNA replication, which were more strongly suppressed in *DOX-BEMF* compared with *DOX-BEM* (Fig. 6–7). These observations are consistent with the ability of cancer-associated fibroblasts to modulate tumour cell proliferation and therapeutic response through paracrine signalling, extracellular matrix remodelling, and induction of stress-adaptive transcriptional programmes [15,45,51].

Higher-resolution pathway analyses reinforced the pattern above, revealing suppression of spindle-assembly checkpoint control, APC/C-mediated mitotic progression, PLK1 signalling, and ATR/ATM-linked checkpoint modules (Table 5), consistent with fibroblast-mediated enforcement of cell-cycle arrest under stress [52]. Homologous recombination repair pathways were uniformly down-regulated, consistent with reduced S/G2-phase engagement (Fig. 7). Because template-dependent repair is tightly coupled to proliferation, this pattern supports a shift toward checkpoint-mediated stabilization rather than enhanced DNA repair capacity, a state characteristic of tolerance-based responses to genotoxic stress [53].

Additional pathway changes (Fig. 7) reinforced this DOX-induced adaptive state of cells within *DOX-BEMF* spheroids, including suppression of Rho GTPase signalling, suggestive of reduced cytoskeletal tension and altered mechanotransduction, potentially shaped by fibroblast–matrix interactions [54], alongside broad down-regulation of mRNA processing pathways with preserved translation initiation and nonsense-mediated decay, a quality-control mechanism that degrades aberrant transcripts and contributes to post-transcriptional regulation of gene expression [55]. Together, these patterns are consistent with selective translation of stress-responsive transcripts and controlled transcriptome remodelling under DOX-induced cytotoxic stress [56,57].

By contrast, genes and pathways preferentially induced in *DOX-BEMF* spheroids were enriched for immune and inflammatory programmes, including interferon signalling, complement activation, cytokine responses, and innate immune–like pathways, together with EMT-associated, metabolic, and xenobiotic stress signatures. Complement pathway enrichment likely reflects activation of innate immune–like signalling within the tumour microenvironment, as both tumour and stromal cells, including fibroblasts, can produce complement components that contribute to paracrine communication and tissue remodelling [58]. This transcriptional profile is characteristic of a stress-adapted state, in which inflammatory signalling and metabolic rewiring support phenotypic plasticity rather than proliferation, consistent with therapy-tolerant cell states described in melanoma and other cancers [59].

Consistent with these transcriptional changes, transcriptome-derived TF activity analysis revealed increased activity of ETS family members, NF-κB, and AP-1 programmes, alongside repression of E2F- and MYBL2-driven networks (Table 6). This is also supported by the pathway-level analysis (Fig. 7) in the assumption that these coordinated shifts integrate extracellular matrix, cytokine, and DNA damage–associated signals into a transcriptional state characterised by proliferative arrest, reduced DNA repair capacity, and enhanced inflammatory signalling [60–63]. Collectively, these features define a stress-adapted, microenvironment-driven phenotype that prioritises survival and phenotypic plasticity over proliferation, consistent with persister-like states described in therapy-exposed cancers [15,59].

Integration with curated DOX resistance gene sets and external transcriptomic datasets revealed strong conservation across heterogenous experimental contexts, including human and mouse data, independent melanoma cell line models and treatment conditions, with consistent downregulation of core proliferation-associated genes (*Mki67, Birc5, Cenpf, Pbk, Cdc6, Tk1,* and *Tyms*) and concordant up-regulation of *IL6* (Table 7). These patterns align with prior studies linking suppression of mitotic regulators and induction of inflammatory signalling to chemotherapy tolerance [59,64], suggesting that fibroblast inclusion is associated with a shift in the cellular stress landscape that couples proliferative suppression with transcriptional stress activation.

By integrating bulk transcriptomics, pathway-level analyses, transcription factor inference, cross-dataset meta-analysis, and RT-qPCR validation, this study establishes a framework for dissecting fibroblast-mediated drug tolerance in melanoma while highlighting the limitations of tumour cell–only models. Interpretation is constrained by the use of bulk transcriptomics, which lacks cell-type–specific resolution, highlighting the need for single-cell or spatially resolved approaches.

In conclusion, our data indicated that fibroblast incorporation into melanoma spheroids co-cultured with macrophages and endothelial cells promotes a shift in the transcriptional response of melanoma spheroids to DOX toward a stress-adaptive response, characterized by enhanced inflammatory and survival-related gene expression together with suppression of proliferative and cell-cycle-associated pathways. Rather than simply attenuating drug effects, fibroblasts reshape both the magnitude and regulatory structure of DOX-induced responses at the levels of gene expression and transcription factor activity. The conservation of these responses across analytical layers and external datasets supports the robustness and biological relevance of this fibroblast-driven program. Together, these findings establish the *BEMF* spheroids model as a robust and physiologically relevant platform for studying TME–mediated chemotherapy tolerance and for evaluating therapeutic strategies targeting stromal–tumour interactions.

## 4. Methods

### 4.1. Development of spheroids mimicking melanoma microenvironment

To model TME–mediated regulation of chemotherapy response, spheroids comprising melanoma, endothelial cells, macrophages, with or without fibroblasts were established and exposed to DOX, as shown below.

#### 4.1.1. Obtaining bone marrow-derived macrophages (BMDMs)

All animal experiments were conducted in strict accordance with the European Directive 2010/63/EU and national legislation (Law 43/2014). The study is reported in accordance with ARRIVE guidelines (under UEFISCDI grant PN-III-P2-2_1-PED-2021-0411, Contract no.659PED/23.06.2022), and were approved by the Babes-Bolyai University Ethics Committee (no.15924/15.11.2022) and National Sanitary Veterinary and Food Safety Authority (Cluj-Napoca, Romania, authorization no. 349/01.02.202). Mice, C57BL/6 (Cantacuzino Institute, Bucharest, Romania), were euthanized by CO₂ anoxia as previously described by our group [65], and all efforts were made to minimize animal suffering. Bone marrow-derived progenitors were isolated from the femurs of 8-week-old male mice [65]. Femurs were harvested in 70% ethanol, and epiphyses were removed to expose the bone marrow cavity. Bone marrow was flushed using standard DMEM culture medium, collected, and centrifuged at 350 × g for 10 min [65]. The resulting cell pellet was resuspended in DMEM supplemented with 10 ng/mL macrophage colony-stimulating factor (M-CSF; Cell Signaling Technology, MA, USA) and maintained in culture for differentiation. Cells were cultured at 37 °C in a humidified atmosphere with 5% CO₂, with medium changes every 2–3 days. After 7 days of differentiation, adherent mature macrophages were obtained and used for spheroids formation in the following experiments (Fig. 10A).

**Figure 10.**
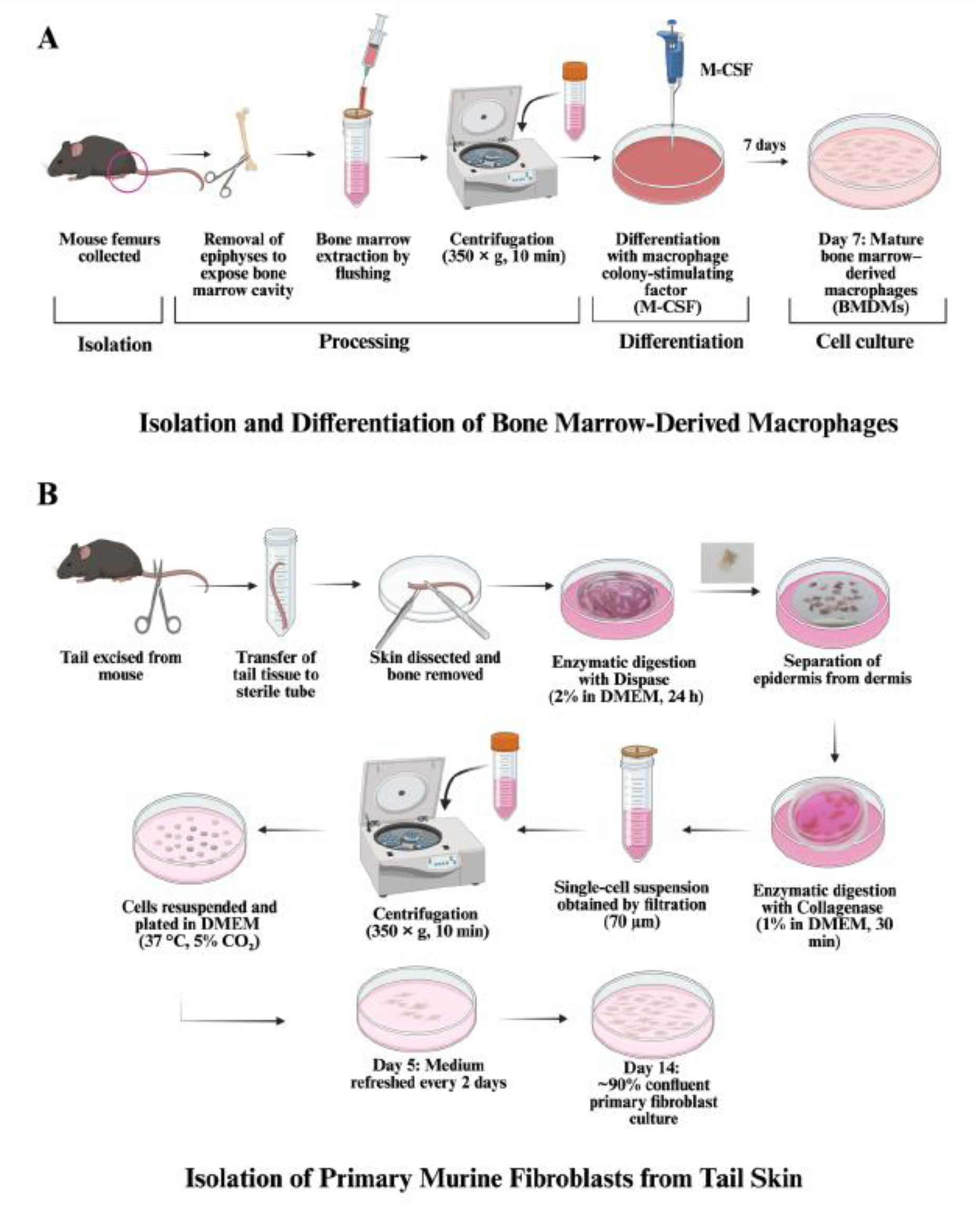
Procedure for obtaining primary cells from mice. **(A)** Isolation and differentiation of bone marrow-derived macrophages; **(B)** Isolation of primary murine fibroblasts from tail skin. Created in BioRender. Gabriela, G. (2026)

#### 4.1.2. Obtaining primary fibroblast culture from murine tail skin

Tails were collected from C57BL/6 mice and immersed in 70% ethanol for surface sterilization and then washed twice in PBS. The skin was carefully peeled, followed by bone removal [66,67]. Remaining connective tissue was removed using sterile curved scissors, and the skin was cut into small pieces (approximately 5 × 10 mm). Skin fragments were transferred into Dispase II solution (2 mg/mL in DMEM) and incubated overnight at 37 °C to facilitate separation of the epidermis. After incubation, the tissue was washed with PBS, and the epidermis was gently separated from the dermis using sterile forceps. After removal of the epidermis, the dermis was washed with PBS and further placed in collagenase solution (1 mg/mL in DMEM) for 30 min at 37 °C, with the dermal side that had been in contact with the epidermis oriented downward. The resulting cell suspension was passed through a 70 µm cell strainer and centrifuged at 350 × g for 10 min. The cell pellet was resuspended in DMEM and seeded under standard culture conditions, in a Petri dish. The culture medium was changed every 2 days. After 14 days, primary fibroblasts reached approximately 90% confluence and were used for subsequent experiments (Fig. 10B).

#### 4.1.3. Endothelial and melanoma cell culture

B16.F10 melanoma (ATCC CRL-6457) and 2H11 endothelial (ATCC CRL-2163) cell lines were cultured in DMEM complete medium with supplements [68] until spheroids formation.

#### 4.1.4. Spheroids formation

The two types of spheroids were formed by liquid overlay technique using ultra-low attachment plates [69]. To obtain the three-cell-type spheroids (*BEM*), B16.F10 melanoma cells (B, 5 × 10³/well) were embedded in 1.5% commercial extracellular matrix (ECM; Sigma-Aldrich, Germany) together with 2H11 endothelial cells (E) and primary macrophages (M), in a ratio of 1:1:4 as previously described [20,70–72]. Four-cell-type spheroids (*BEMF*) were generated by additionally incorporating primary fibroblasts (F) in a final ratio between melanoma cells, 2H11 endothelial cells, primary macrophages, and primary fibroblasts (F) of 1:1:4:1 [22,73]. The selected cell ratios were chosen to approximate the cellular composition of the *in vivo* melanoma TME, enabling physiologically relevant tumour–vascular, immune, and stromal crosstalk that collectively shapes cancer cell behaviour and therapy response [42]. Plates were centrifuged for 15 min, at 350*×g* and incubated at 37° C with 5% CO₂, for 4 days to allow spheroids formation.

### 4.2. Spheroids viability upon DOX exposure

4-day old melanoma spheroids (500-600 µm in diameter) were treated with DOX (Sigma-Aldrich, Germany) at concentrations ranging from 0.156 to 10 µM, while untreated spheroids were maintained as controls (CTR). After 48 h of DOX exposure, cell viability was measured to generate dose–response curves and to determine IC₅₀ and IC₃₀ values of DOX. The corresponding IC₃₀ concentration of DOX was selected and used in all subsequent experiments, to impose sublethal drug stress, and allowing the analysis of early resistance-associated transcriptional responses while maintaining spheroids viability. Spheroid morphology and diameter was assessed by light microscopy using an Axio Vert.A1 FL microscope (Carl Zeiss, Germany).

Cell viability was measured using a modified acid phosphatase (APH) assay, as previously described [74]. Briefly, 100 µL of APH reaction buffer (0.2 M acetate buffer, pH 5.8, 0.1% Triton X-100, and 2 mg/mL p-nitrophenyl phosphate) was added to each well and incubated for 30 min at 37 °C. The reaction was terminated by addition of 10 µL of 1 M NaOH, and absorbance was measured at 405 nm within 10 min using a FLUOstar Omega microplate reader (BMG Labtech, Offenburg, Germany). Cell viability within spheroids exposed to DOX was calculated as a percentage of the viability measured for cells within control spheroids. All experiments were performed in triplicate. Data are presented as mean ± SD.

### 4.3. RNA extraction, library prep and sequencing

After 48 h of exposure to DOX, a total of eight samples were collected and washed twice with phosphate-buffered saline (PBS), namely two biological replicates from each treated condition (*BEM*/*BEMF* exposed to DOX and two from each corresponding control (*BEM*/*BEMF*) [75]. Total RNA was extracted using the RNeasy Mini Kit (Qiagen, 74104) according to the manufacturer’s instructions [76]. Briefly, spheroids were lysed in RLT buffer, and the lysates were homogenized and loaded onto RNeasy spin columns. After on-column washing steps, RNA was eluted in 40 µl of RNase-free water. RNA concentration and purity were assessed using a NanoDrop 1000 spectrophotometer (Thermo Scientific, MA, USA). Strand-specific cDNA libraries were prepared using the TruSeq Stranded mRNA Library Prep Kit (Illumina, USA), followed by quality control performed by Macrogen Europe (Netherlands). Paired-end sequencing was carried out at Macrogen facilities using the Illumina NovaSeq 6000 platform.

### 4.4. RNA-Seq preprocessing and quantification

Raw paired-end sequencing reads were quality controlled using *fastp* [77] (v0.23.4) with default parameters. Quality-control metrics from RnaSeq reads preprocessing and transcript quantification were aggregated and inspected using *MultiQC* [78] (v1.28). Transcript abundances were quantified using Salmon [79] (v1.10.3) in mapping-based mode with automatic library-type detection. A decoy-aware Salmon index (k = 31) was constructed using GENCODE vM36 transcript annotations [80] and the GRCm39 primary genome assembly by concatenating transcriptome and genome FASTA files to generate a decoy-aware reference transcriptome (*gentrome*).

### 4.5. Differential expression modelling

Transcript-level abundance estimates generated by Salmon were imported into *R* [81] (v4.5.1) using *tximport* [82] (v1.36.1) and summarized to gene-level counts based on GENCODE vM36 transcript–gene annotations. Gene-level counts were normalized using *DESeq2* [83] (v1.48.2) and genes with low expression were filtered by requiring a minimum normalized count of 10 in at least two samples.

Variance-stabilizing transformation (VST) was applied for exploratory analyses and data visualization. Sample relationships and data quality were assessed using principal component analysis (PCA), hierarchical clustering, and sample-to-sample distance metrics. Differential expression analysis was performed using *DESeq2* with a generalized linear model incorporating cell composition, treatment, and their interaction (model formula: *∼cell composition + treatment + cell composition:treatment*). Log_2_ fold changes were shrunken using the *apeglm* method [84] (v1.30.0). Differential expression results were annotated using Ensembl, MGI, and Entrez Gene identifiers via the package *biomaRt* [85] (v2.64.0) and *org.Mm.eg.db* [86] (v3.21.0), and multiple-testing correction was applied using the Benjamini–Hochberg procedure [87].

### 4.6. Pathway analysis

Gene Set Enrichment Analysis (GSEA) was performed to assess pathway-level changes associated with fibroblast inclusion in the spheroid models under DOX exposure. Gene-level statistics were derived from the differential expression model using DESeq2, and results were ranked by signed log₂ fold change to generate an ordered gene vector used as input for enrichment analysis [88].

#### 4.6.1. MSigDB Hallmark GSEA

High-level functional analysis was performed by comparing the ranked gene list to the Molecular Signatures Database (MSigDB) Mouse Hallmark gene sets to identify biologically coherent major transcriptional programs associated with fibroblast inclusion under DOX exposure. Pathway annotations were retrieved using the *msigdbr* R package [89] (v25.1.1) by filtering just to the “Hallmark” collection for *Mus musculus*. GSEA was performed using the *GSEA* function from the *clusterProfiler* R package [90] (v4.16) with the *fgsea* algorithm [91].

Genes contributing to enriched Hallmark pathways were converted to human-readable gene symbols using the *setReadable* function with the *org.Mm.eg.db* R package. Gene set descriptions and associated metadata were merged with the enrichment results. Pathways were classified as activated or suppressed based on the sign of the normalized enrichment score (NES).

#### 4.6.2. Reactome GSEA

To dissect changes at a finer resolution and identify gene sets highlithing specific molecular interactions and reactions, pathway analysis on Reactome curated gene sets was applied. Pathway enrichment was performed by running GSEA on curated pathways from the Reactome database using the *gsePathway* function from the *ReactomePA* R package [92] (v1.52.0). The same ranked gene list and analysis parameters used for the MSigDB analysis were applied.

#### 4.6.3. Reactome pathway embedding, clustering, and semantic cluster labelling

Curated pathway annotations encode structured biological knowledge that can be quantitatively compared within a semantic embedding space. Clustering pathway representations in such a space therefore provides an approach to reduce overlap and redundance in results and identify higher-order functional programs among enriched pathways [93,94]. Pathway descriptions were embedded into a shared semantic space using pretrained language models, followed by unsupervised clustering to group functionally related pathways. For each Reactome pathway significantly enriched in the analysis (FDR < 0.05), the pathway description was converted into a numerical embedding using the OpenAI *text-embedding-3-large* model (OpenAI API, v2024-01), which produces 3,072-dimensional dense vector representations of text. Embeddings were generated in mini-batches to ensure consistent formatting and to minimize rate-limit–related artifacts.

The resulting embedding matrix was standardized and clustered using *k-means clustering* [95]. The optimal number of clusters was determined by inspection of the silhouette index and the total within-cluster sum of squares [96], yielding an optimal solution of eight clusters.

To visualize pathway similarity in two dimensions, Uniform Manifold Approximation and Projection (UMAP) was applied using the *umap* R package [97] (v0.2.10.0). Cluster-level biological themes were generated using a large language model (LLM). For each cluster, the corresponding Reactome pathway names were provided to an Anthropic Claude model (*claude-3.5-sonnet*, Anthropic API, v2024-10) with a structured prompt requesting a concise (2–5 words), mechanistic, and biologically meaningful theme summarizing the shared functional content of the cluster. The model returned a single short label per cluster, which was used for annotation of UMAP visualizations and downstream summary tables. The temperature parameter was fixed at 0 to ensure deterministic and reproducible labelling.

This exploratory workflow was used to reduce redundancy among significantly enriched Reactome pathways and facilitate visual interpretation. Clusters were derived from semantic similarity of pathway descriptions rather than pathway topology, gene overlap, or enrichment statistics, and therefore should be interpreted as descriptive groupings rather than formally inferred biological modules.

### 4.7. Transcription factor activity analysis

Variance-stabilized gene expression values were obtained using *DESeq2* and extracted as an assay matrix formatted as *samples × genes* for downstream analyses [98]. Ensembl gene identifiers were harmonized by removing version suffixes and mapped to gene symbols using the *bitr* function from *clusterProfiler* in conjunction with the *org.Mm.eg.db* annotation database. Only genes with successful mappings were retained.

Transcription factor (TF) activities were inferred based on the mouse DoRothEA TF–target regulon resource (*dorothea_mm*) retrieved via the *dorothea* R package [99] (v1.20.0) and restricting the analysis to high- and medium-confidence regulatory interactions (confidence levels A–C). Regulons were further filtered to include only target genes present in the expression matrix and to TFs with a minimum of four targets, to ensure robust activity estimation.

TF activities were computed using the *decoupleR* package [100] (v2.14.0) employing the univariate linear model (ULM) method, which estimates TF activity scores per sample based on the expression of target genes and signed mode-of-regulation annotations.

Differential TF activity was assessed using a factorial design with two experimental factors: cell composition (*BEMF* vs. *BEM*) and treatment (DOX vs. control), yielding four conditions. Linear models were fit as *activity* ∼ *cell composition* × *treatment*, and the interaction term was used to quantify whether DOX-induced TF activity changes differed between fibroblast-free and fibroblast-containing spheroids.

### 4.8. Resistance gene curation and comparative analysis

#### 4.8.1. Resistance gene set construction

Genes associated with DOX resistance in melanoma were identified by querying annotations from the NCBI Gene database using the *rentrez* R package [101] (v1.2.4). Searches were performed using the combined terms “doxorubicin resistance” and “melanoma.” The search returned a set of human gene identifiers, which were subsequently mapped to mouse orthologs using Ensembl BioMart via the *biomaRt* R package. Genes encoding microRNAs were excluded from downstream analyses.

Additional candidate genes were obtained from curated gene set resources by querying the Molecular Signatures Database (MSigDB) for DOX-related mouse gene sets using the *msigdbr* R package. All mouse gene sets were retrieved, and those containing the term “doxorubicin” or “DOX” in their gene set name were identified. To increase the likelihood of involvement in resistance mechanisms, only genes present in at least three independent gene sets were retained.

To assess whether the retrieved genes were specifically associated with DOX resistance or reflected more general resistance mechanisms, a second NCBI Gene search was performed to identify genes annotated as multidrug resistance–related in melanoma. As before, human gene identifiers were subsequently mapped to mouse orthologs.

#### 4.8.2. Comparative analysis of resistance-associated transcriptional responses

Treatment-induced transcriptional responses between models were compared by extracting log₂ fold change estimates for genes in the extended resistance gene set from both *BEM* and *BEMF* differential expression results. For each gene, the absolute difference in treatment response magnitude between models was calculated as:

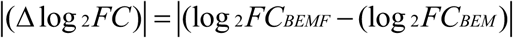

Genes showing |Δlog₂FC| ≥ 1 were classified as exhibiting model-dependent differences in response magnitude. To avoid inclusion of genes lacking robust evidence of treatment responsiveness, genes that failed to reach statistical significance (FDR > 0.05) in both models were excluded from this subset.

Normalized gene expression values were log₂-transformed and Z-score scaled across samples prior to visualization. Expression heatmaps were generated using the *ComplexHeatmap* R package [102] (v2.24.1), with hierarchical clustering applied to genes and treatment status displayed as a column annotation. Fold-change heatmaps were constructed using unscaled log₂ fold changes, with genes ordered by response magnitude differences between models. Scatter plots comparing *BEM* and *BEMF* treatment responses were generated using *ggplot2*, with point colour and size encoding the absolute difference in log₂ fold change between models. Directional concordance of treatment responses was assessed by evaluating the sign consistency of log₂ fold changes across models.

Where applicable, Ensembl gene identifiers were mapped to MGI gene symbols using *biomaRt* R package. In cases where gene symbols were unavailable, Ensembl identifiers were retained to ensure unambiguous reporting.

### 4.9. Public dataset selection, processing and analysis

Public transcriptomic datasets relevant to DOX response in melanoma were identified through systematic searches of the NCBI Gene Expression Omnibus (GEO) using keywords related to melanoma, DOX treatment, and chemotherapy resistance. Twelve candidate datasets were initially identified, of which three were retained following manual curation based on experimental relevance, availability of treated and control conditions, and suitability for comparative analysis (GSE212112, GSE246690, and GSE33624).

Expression data were retrieved using the *GEOquery* R package [103] (v2.76.0). Expression or counts matrices provided by the original authors were used without reprocessing raw sequencing or microarray data. Selected datasets comprised both RNA-seq and microarray platforms and included human and mouse melanoma cell lines exposed to DOX under diverse experimental conditions. Human gene identifiers were mapped to *Mus musculus* orthologs to enable cross-species comparison.

Differential expression analysis was performed for datasets with appropriate experimental designs using *DESeq2* [83] for RNA-Seq studies and *limma* [104] (v3.64.3) for microarray studies.

### 4.10. Cross-study integration and directionality meta-analysis

#### 4.10.1 Cross-study overlap analysis

Top differentially expressed genes from public RNA-Seq datasets were integrated with statistically significant differentially expressed genes (DEGs) identified in the *BEM* and *BEMF* experimental models to define resistance-associated genes exhibiting consistent DOX-responsive regulation across systems. Given the heterogeneity of public datasets with respect to species, platforms, treatment conditions, and replication, cross-dataset comparisons emphasized concordance in the direction of transcriptional response rather than absolute effect size.

Significantly upregulated and downregulated genes (FDR < 0.05 and absolute log₂ fold change > 1) from the two RNA-Seq datasets (GSE212112 and GSE246690) were intersected with the significant differential expression results from the DOX-treated *BEM* and *BEMF* spheroid models to assess overlap in resistance-associated transcriptional programs. We also intersected the common identified genes with the curated list of resistance genes.

#### 4.10.2 Directionality meta-analysis

To assess whether selected genes exhibited consistent transcriptional regulation across multiple independent experimental systems and public melanoma datasets, log_2_ fold-change estimate (log2FC) values were harmonized into a gene-by-dataset matrix. We employed a directionality-based meta-analysis that aggregates signed effects rather than relying on classical effect-size meta-analysis. For each gene, study-specific log₂ fold-changes were transformed into signed effect-size scores:

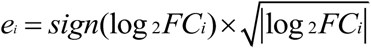

to preserve direction while down-weighting extreme magnitudes. Gene-level directional consistency across studies was summarized by an aggregated signed score:

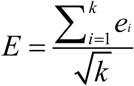

where *k* is the number of available studies. Statistical significance was assessed heuristically using a normal approximation to derive p-values, followed by Benjamini–Hochberg correction for multiple testing.

This approach is well suited for integrative analyses in which only effect direction and relative magnitude are consistently available and is robust to heterogeneity across study designs, a setting in which classical p-value or variance-based meta-analysis methods may be suboptimal [105]. Genes were classified as consistently up or downregulated if they exhibited a significant aggregated signed score (adjusted q<0.05) with concordant directionality across datasets. All analyses were performed in R (v4.5.1 or later) using *readODS* for data import, *dplyr*, *tidyr*, and *tibble* for data manipulation, as well as *base* functions for statistical testing.

### 4.11. RT-qPCR validation

Gene expression changes were validated by reverse transcription quantitative PCR (RT-qPCR). Briefly, 1 µg of total RNA was reverse transcribed into cDNA using iScript™ Reverse Transcription Supermix (#1708841, Bio-Rad Laboratories, Gladesville, Australia) in a final reaction volume of 20 µL [106]. The reverse transcription reaction was carried out with an initial priming step at 25 °C for 5 min, followed by cDNA synthesis at 46 °C for 20 min and enzyme inactivation at 95 °C for 1 min. Quantitative PCR was then performed with SsoFast™ EvaGreen® Supermix (#172-5200, Bio-Rad Laboratories, Gladesville, Australia) [106]. Each reaction contained 1 µL of cDNA and 500 nM of gene-specific primers (sequences listed in Table S4; Eurogentec, Seraing, Belgium). All reactions were run on a CFX96™ Real-Time PCR System (C1000 Touch Thermal Cycler, Bio-Rad Laboratories, Gladesville, Australia) according to the manufacturer’s instructions.

Quantification cycle (Ct) values were obtained for target genes and normalized to β-actin as the internal reference gene [107]. RT-qPCR measurements were performed on biological triplicates for each condition. Samples were stratified by cell composition (*BEM* or *BEMF*) and treatment condition (control or DOX), following recommended best practices for quantitative PCR experiments [108].

For each sample, ΔCt values were calculated as the difference between Ct of the target gene and that of the reference gene Ct (ΔCt = Ct_target_ − Ct_reference_). Within each cell composition, the mean ΔCt of control samples was used as the calibrator. ΔΔCt values were computed as the difference between sample-specific ΔCt values and the corresponding control mean ΔCt. Relative gene expression was determined using the comparative Ct (ΔΔCt) method and expressed as log2 fold change, calculated as −ΔΔCt, using the comparative Ct (ΔΔCt) method [107].

### 4.12. Statistical analysis

For the viability assays one-way Anova was performed using GraphPad Prism 7, followed by Dunnett’s multiple comparisons test. DOX IC_50_ values were calculated via nonlinear regression of dose-response curves. To assess differences between conditions, dose–response curves were compared using an extra sum-of-squares F-test. IC_30_ was calculated in Excel using: ICx=[(x/100-x)^1/Hillslope]*IC50. In addition, area under the curve (AUC) values were calculated for each biological replicate using trapezoidal integration of viability values across log₁₀-transformed DOX concentrations in R, and AUC distributions between conditions were compared using an unpaired two-sided t-test.

For statistical assessment of expression differences between RT-qPCR results from treatment and control conditions, two-sided t-tests were performed on ΔCt values, which represent the appropriate scale for statistical inference in this type of analysis and more closely satisfy assumptions of normality than fold-change values [109]. Summary statistics are reported as mean ± standard deviation across biological replicates. Significance was set at p < 0.05 (*), with **p < 0.01, ***p < 0.001, ****p < 0.0001; ns = not significant.

To compare the proportion of genes exceeding predefined absolute log₂ fold-change thresholds between conditions, McNemar’s test was applied to paired binary classifications (above vs. below threshold), with each gene treated as a matched observation across models.

All transcriptomic data normalization, statistical analysis, and visualization were performed in R v4.5.1.

## Author Contributions

IOP: Methodology, Investigation, Data Curation, Formal Analysis, Validation, Writing Original Draft; G.G.N.: Methodology, Validation; B.D. Methodology; E.L Methodology; V.F.R Methodology; S.M Methodology; S.D. Methodology; L.P. Methodology; M.B.: Conceptualization, Supervision, Review & Editing Supervision; A.S.: Conceptualization, Review & Editing, Funding Acquisition, Project Administration.

## Acknowledgements

This work was supported by L’Oréal - UNESCO “For Women in Science” [Fellowship Programme no. 914, 2020]; the UEFISCDI grant [PN-III-P2-2_1-PED-2021-0411, No. 659PED, 2022] granted to dr. A.S.

## Data availability

The RNA sequencing data generated in this study have been deposited in the NCBI Gene Expression Omnibus (GEO) and are accessible through GEO Series accession number GSE336658.

## Consent

The authors have nothing to report.

## Ethics Statement

All animal experiments were conducted in strict accordance with the European Directive 2010/63/EU and national legislation (Law 43/2014). The study is reported in accordance with ARRIVE guidelines (under UEFISCDI grant PN-III-P2-2_1-PED-2021-0411, Contract no.659PED/23.06.2022), and were approved by the Babes-Bolyai University Ethics Committee (no. 15924/15.11.2022) and National Sanitary Veterinary and Food Safety Authority (Cluj-Napoca, Romania, authorization no. 349/01.02.202).

## Declaration of generative AI in scientific writing

The authors acknowledge the use of generative AI (ChatGPT 5.1) in enhancing the readability and language of the manuscript. After using the tool, the authors thoroughly reviewed and edited the content, taking full responsibility for the final manuscript.

## Supporting Information

**Table S1.** Raw quantitative real-time PCR cycle threshold (Ct) values for doxorubicin-responsive target genes in BEM and BEMF spheroids.

**Table S2.** Gene set enrichment analysis of MSigDB Hallmark pathways for the fibroblast-dependent (BEMF versus BEM) doxorubicin-interaction response.

**Table S3.** Gene set enrichment analysis of Reactome pathways for the fibroblast-dependent (BEMF versus BEM) doxorubicin-interaction response.

**Table S4.** qRT-PCR sequences for used primers

## Abbreviations

APH: Acid phosphatase assay
ATCC: American Type Culture Collection
BEM: Spheroids with Melanoma cells (B), endothelial cells (E), and macrophages (M)
BEMF: Spheroids with Melanoma cells (B), endothelial cells (E), macrophages (M), and Fibroblasts (F)
BMDMs: Bone marrow–derived macrophages
CAFs: Cancer-associated fibroblasts
cDNA: Complementary DNA
DEG: Differentially expressed gene
DMEM: Dulbecco’s Modified Eagle Medium
DOX: Doxorubicin
ECM: Extracellular matrix
EMT: Epithelial–mesenchymal transition
FDR: False discovery rate
FGSEA: Fast Gene Set Enrichment Analysis
GEO: Gene Expression Omnibus
GSEA: Gene Set Enrichment Analysis
IC₃₀: the concentration of a compound that results in 30% inhibition of cell growth
IC₅₀: the concentration of a compound that results in 50% inhibition of cell growth
IL: Interleukin
LLM: Large language model
MAPK: Mitogen-activated protein kinase
M-CSF: Macrophage colony-stimulating factor
MDR: Multidrug resistance
MIQE: Minimum Information for Publication of Quantitative Real-Time PCR Experiments
MSigDB: Molecular Signatures Database
NES: Normalized enrichment score
NMD: Nonsense-mediated decay
PBS: Phosphate-buffered saline
PCA: Principal component analysis
RT-qPCR: Reverse transcription quantitative polymerase chain reaction
STAT: Signal transducer and activator of transcription
TF: Transcription factor
TME: Tumour microenvironment
ULM: Univariate linear model
UMAP: Uniform Manifold Approximation and Projection
VST: Variance-stabilizing transformation

