## Supporting Information for "Fibroblast-Enhanced Tumour Microenvironment Signalling Promotes Adaptive Doxorubicin Tolerance in Heterotypic Melanoma Spheroids"

**Table S1.** Raw quantitative real-time PCR cycle threshold (Ct) values for doxorubicin-responsive target genes in BEM and BEMF spheroids.

*Ct values for the reference gene (Actin) and 25 target genes across all samples (n = 3 biological replicates per cell composition and treatment), used to calculate the relative expression changes*

| Sample | BEM_CTR_R1 | BEM_CTR_R2 | BEM_CTR_R3 | BEM_DOX_R1 | BEM_DOX_R2 | BEM_DOX_R3 | BEMF_CTR_R1 | BEMF_CTR_R2 | BEMF_CTR_R3 | BEMF_DOX_R1 | BEMF_DOX_R2 | BEMF_DOX_R3 |
|---|---|---|---|---|---|---|---|---|---|---|---|---|
| Replicate | R1 | R2 | R3 | R1 | R2 | R3 | R1 | R2 | R3 | R1 | R2 | R3 |
| Ct_Actin | 22.12 | 22.2 | 22.45 | 22.47 | 22.58 | 22.52 | 21.38 | 21.35 | 21.46 | 22.01 | 21.84 | 22.31 |
| Ct_Tk1 | 30.24 | 30.46 | 30.3 | 31.43 | 31.68 | 31.19 | 26.62 | 26.52 | 27.52 | 29.64 | 29.88 | 30.03 |
| Ct_Cenpf | 31.11 | 31.18 | 31.5 | 33.52 | 33.59 | 34.79 | 34.16 | 34.14 | 34.35 | 35.65 | 35.53 | 35.73 |
| Ct_Slc29a2 | 31.57 | 31.76 | 31.51 | 30.81 | 30.99 | 30.66 | 30.93 | 30.63 | 30.65 | 31.98 | 31.57 | 31.78 |
| Ct_Mki67 | 31.73 | 31.82 | 30.47 | 34.42 | 34.04 | 33.86 | 31.1 | 31.08 | 30.53 | 31.62 | 31.8 | 30.95 |
| Ct_Lrr1 | 34.07 | 34.59 | 34.03 | 35.52 | 35.49 | 35.72 | 31.88 | 32.06 | 31.6 | 34.8 | 35.28 | 34.55 |
| Ct_Foxf2 | 37.59 | 37.91 | 37.27 | 36.67 | 36.29 | 35.73 | 35.44 | 35.06 | 34.43 | 35.99 | 35.46 | 35.1 |
| Ct_Etv4 | 34.27 | 34.18 | 34.41 | 34.58 | 34.31 | 34.49 | 32.34 | 32.08 | 32.21 | 32.09 | 32.05 | 32.93 |
| Ct_Tfap4 | 30.01 | 30.23 | 30.54 | 30.9 | 31.07 | 31.31 | 28.46 | 28.57 | 28.6 | 29.46 | 29.41 | 29.56 |
| Ct_Topbp1 | 28.99 | 29.1 | 29.14 | 29.07 | 29.12 | 29.27 | 27.39 | 27.36 | 28.04 | 28.42 | 28.53 | 28.8 |
| Ct_Cxcl12 | 33.54 | 33.77 | 33.34 | 34.57 | 35.11 | 34.71 | 34.53 | 34.63 | 35.1 | 34.59 | 34.71 | 35.63 |
| Ct_Lum | 31.37 | 31.48 | 31.79 | 32.13 | 32.19 | 32.16 | 30.16 | 30.38 | 30.36 | 30.09 | 30.06 | 30.18 |
| Ct_Medag | 32.23 | 32.22 | 31.93 | 34.53 | 33.91 | 34.35 | 33.51 | 33.32 | 33.73 | 37.33 | 35.93 | 35.05 |
| Ct_Ncapg2 | 29.87 | 29.79 | 29.8 | 31.22 | 31.05 | 31.22 | 27.77 | 27.86 | 28.67 | 31.56 | 30.86 | 32.38 |
| Ct_Tyms | 30.27 | 30.46 | 30.44 | 31.47 | 31.65 | 31.33 | 28.88 | 28.6 | 29.11 | 29.83 | 30.27 | 30.25 |
| Ct_Siva1 | 27.45 | 27.39 | 27.42 | 26.69 | 26.55 | 26.46 | 26.99 | 26.97 | 26.79 | 26.21 | 26.24 | 26.13 |
| Ct_Cgas | 29.92 | 30.43 | 30.23 | 31.03 | 31.11 | 30.96 | 30.25 | 30.77 | 30.52 | 31.38 | 32.15 | 31.61 |
| Ct_Pdss | 29.42 | 30.05 | 30.41 | 30.97 | 30.23 | 30.53 | 30.7 | 30.78 | 30.56 | 31.71 | 31.88 | 31.86 |
| Ct_Abcb1a | 35.39 | 35.18 | 35.93 | 35.9 | 36.87 | 36.04 | 36.62 | 36.89 | 36.8 | 35.11 | 34.65 | 34.55 |
| Ct_Foxm1 | 31.79 | 31.67 | 31.64 | 32.75 | 33.15 | 32.77 | 28.37 | 27.99 | 27.9 | 29.47 | 29.16 | 29.21 |
| Ct_Myc | 27.82 | 27.83 | 27.84 | 28.31 | 28.23 | 28.26 | 27.17 | 27.12 | 27.03 | 28.09 | 27.9 | 27.71 |
| Ct_Prdm | 36.99 | 37.16 | 36.69 | 35.75 | 35.69 | 36.18 | 36 | 36.37 | 35.9 | 36.09 | 35.29 | 35.35 |
| Ct_Pbk | 27.42 | 27.53 | 27.44 | 29.07 | 29.02 | 29.15 | 29.68 | 29.7 | 29.47 | 31.18 | 31.35 | 31.49 |

| | | | | | | | | | | | | |
|---|---|---|---|---|---|---|---|---|---|---|---|---|
| Ct_Birc5 | 28.91 | 28.91 | 28.73 | 29.52 | 29.63 | 29.57 | 26.01 | 26.12 | 25.97 | 27.78 | 27.69 | 28.06 |
| Ct_Ptgs2 | 29.85 | 29.76 | 30 | 29.5 | 29.59 | 29.56 | 30.45 | 30.37 | 30.5 | 29.32 | 29.42 | 29.39 |
| Ct_Il6 | 29.41 | 29.36 | 29.39 | 29.28 | 29.36 | 29.19 | 30.81 | 30.79 | 30.85 | 29.77 | 29.75 | 29.78 |

**Table S2.** Gene set enrichment analysis of MSigDB Hallmark pathways for the fibroblast-dependent (BEMF versus BEM) doxorubicin-interaction response.

| ID | Description | Set Size | Enrichment Score | NES | Pvalue | P.adjusted | Qvalue | Rank | leading_edge |
|---|---|---|---|---|---|---|---|---|---|
| HALLMARK_G2M_CHECKPOINT | HALLMARK_G2M_CHECKPOINT | 198 | -0.761639878 | -2.386806918 | 2.07e-22 | 1.04e-20 | 6.56e-21 | 1737 | tags=47% |
| HALLMARK_E2F_TARGETS | HALLMARK_E2F_TARGETS | 200 | -0.716918206 | -2.256408161 | 4.28e-17 | 1.07e-15 | 6.77e-16 | 1737 | tags=40% |
| HALLMARK_MITOTIC_SPINDLE | HALLMARK_MITOTIC_SPINDLE | 199 | -0.679520444 | -2.133908345 | 4.69e-13 | 7.83e-12 | 4.94e-12 | 2474 | tags=45% |
| HALLMARK_COMPLEMENT | HALLMARK_COMPLEMENT | 152 | 0.583315553 | 1.908028535 | 2.56e-05 | 0.000320248 | 0.000202262 | 1150 | tags=21% |
| HALLMARK_MYC_TARGETS_V2 | HALLMARK_MYC_TARGETS_V2 | 58 | -0.699104218 | -1.870772749 | 5.72e-05 | 0.000572474 | 0.000361563 | 3322 | tags=66% |
| HALLMARK_EPITHELIAL_MESENCHYMAL_TRANSITION | HALLMARK_EPITHELIAL_MESENCHYMAL_TRANSITION | 181 | 0.518927682 | 1.708362749 | 0.000134181 | 0.001118176 | 0.000706216 | 1230 | tags=23% |
| HALLMARK_UNFOLDED_PROTEIN_RESPONSE | HALLMARK_UNFOLDED_PROTEIN_RESPONSE | 111 | -0.593596598 | -1.736210483 | 0.000161676 | 0.001154826 | 0.000729364 | 2230 | tags=41% |
| HALLMARK_P53_PATHWAY | HALLMARK_P53_PATHWAY | 189 | 0.506738328 | 1.674549381 | 0.000405841 | 0.002536506 | 0.001602004 | 1758 | tags=35% |
| HALLMARK_MYC_TARGETS_V1 | HALLMARK_MYC_TARGETS_V1 | 201 | -0.508395033 | -1.601756929 | 0.000459317 | 0.002551762 | 0.001611639 | 3135 | tags=37% |
| HALLMARK_COAGULATION | HALLMARK_COAGULATION | 93 | 0.595886957 | 1.843680891 | 0.001319059 | 0.005995725 | 0.003786774 | 1249 | tags=27% |
| HALLMARK_XENOBIOTIC_METABOLISM | HALLMARK_XENOBIOTIC_METABOLISM | 152 | 0.51054407 | 1.669992592 | 0.001207452 | 0.005995725 | 0.003786774 | 1438 | tags=27% |
| HALLMARK_ALLOGRAFT_REJECTION | HALLMARK_ALLOGRAFT_REJECTION | 154 | 0.493414017 | 1.60588586 | 0.003080907 | 0.011849642 | 0.007483984 | 544 | tags=12% |
| HALLMARK_INTERFERON_GAMMA_RESPONSE | HALLMARK_INTERFERON_GAMMA_RESPONSE | 183 | 0.481327782 | 1.589277841 | 0.002986791 | 0.011849642 | 0.007483984 | 1809 | tags=23% |
| HALLMARK_APICAL_SURFACE | HALLMARK_APICAL_SURFACE | 35 | 0.649397662 | 1.726554348 | 0.004427163 | 0.015811295 | 0.009986081 | 96 | tags=11% |
| HALLMARK_REACTIVE_OXYGEN_SPECIES_PATHWAY | HALLMARK_REACTIVE_OXYGEN_SPECIES_PATHWAY | 46 | 0.601533423 | 1.65436458 | 0.011444128 | 0.031789245 | 0.020077418 | 1992 | tags=46% |
| HALLMARK_KRAS_SIGNALING_UP | HALLMARK_KRAS_SIGNALING_UP | 159 | 0.448279845 | 1.459205984 | 0.010314351 | 0.031789245 | 0.020077418 | 386 | tags=12% |
| HALLMARK_ADIPOGENESIS | HALLMARK_ADIPOGENESIS | 187 | 0.433155422 | 1.429950614 | 0.009837477 | 0.031789245 | 0.020077418 | 2895 | tags=41% |
| HALLMARK_OXIDATIVE_PHOSPHORYLATION | HALLMARK_OXIDATIVE_PHOSPHORYLATION | 197 | 0.418733917 | 1.402533851 | 0.010828619 | 0.031789245 | 0.020077418 | 3363 | tags=44% |

**Table S3.** Gene set enrichment analysis of Reactome pathways for the fibroblast-dependent (BEMF versus BEM) doxorubicin-interaction response.

| ID | Description | Set Size | Enrichment Score | NES | Pvalue | P.adjusted | Qvalue | Rank | Leading Edge | Core Enrichment |
|---|---|---|---|---|---|---|---|---|---|---|
| R-MMU-1640170 | Cell Cycle | 541 | -0.576237173 | -1.982816962 | 6.77E-16 | 9.21E-13 | 8.38E-13 | 1688 | tags=25%, list=11%, signal=23% | Blm/Pold1/Pola2/Skp2/Pif1/Nhp2/Nipbl/Pola1/Zw10/Pot1a/Espl1/Orc1/Ckap5/Fignl1/Rbm39/Npm1/Ncapd3/Hsp90aa1/Cdk1/Ppp1r12a/Clasp1/Topbp1/Pcna/A730008H23Rik/Nup98/Ncapd2/Kpnb1/Nup133/Anapc1/Akt3/Esco1/Rsf1/Pds5b/Smarca5/Mcm8/Terf2/Tubb5/Dscc1/Cdca5/Dynll1/Nup155/Sfi1/Tubgcp4/Lig1/Taok1/Cdc25b/Ranbp2/Mcm6/Ccnb1/Akap9/Trp53bp1/Gar1/Mcm4/Cdt1/Atrx/Pcnt/Mdc1/Rfc5/Tpr/Cnep1r1/Spdl1/Ncapg/Tuba1a/Bard1/Mis18a/Nsd2/Pcm1/Lmnb1/Cks1b/Ccnd1/Ankrd28/Brca1/Cenpa/Tpx2/Cep152/Cpap/Nup107/Rcc1/Herc2/Chek1/Mcm3/Anapc4/Cep192/Tubb2a/Dync1h1/Dkc1/Mcm10/Cdk2/Brip1/Tubb2b/Cep135/Cenph/Plk1/Spc24/Tubb3/Ncapg2/Smc2/Sgo2a/Pole2/Aurka/Esco2/Bub1/Kif18a/Fbxo5/Bub1b/Pmf1/Cep290/Cenpe/Mdm4/Kif23/Ube2c/Aurkb/Kif2c/Ccna2/Chtf18/Nuf2/Nek2/Mcm5/Ercc6l/Cdc45/Pole/Cdc20/Pkmyt1/Cdc6/Rad51/Ndc80/Sgo1/Hmmr/Clspn/E2f2/Exo1/Ccnb2/Kntc1/Kif20a/Cenpu/Cenpf |
| R-MMU-168249 | Innate Immune System | 700 | 0.504270187 | 1.90741899 | 4.39E-13 | 2.99E-10 | 2.72E-10 | 2583 | tags=31%, list=17%, signal=27% | H2-Q7/Serping1/Pld4/Crispld2/Pirb/Nfam1/C1qc/C1qa/Mndal/C1qb/Cybb/Cxcl1/Ifi211/Itgb2/Cfp/Fcer1g/Lyz2/Ctss/B2m/Ncf1/C3ar1/Arg1/Cd93/H2-K1/Nckap1l/Tnfrsf1b/Usp18/Tyrobp/Fth1/Cd53/Clec4d/Pycard/Qsox1/Txnip/Mmp8/Adam8/Elmo2/Cyba/Fos/H2-T22/Ctsh/Cst3/P2rx7/H2-T23/Fuca1/Gyg1/Ctsd/C2/Man2b1/Aga/Prcp/Aprt/Gstp2/Nlrx1/Arl8a/Padi2/Cdc34/Cat/Npc2/Cd14/Fabp5/Pld3/Cd81/Hk3/Psap/Ostf1/Hras/Slpi/Creg1/Stom/Ptk2/Gsn/Grn/Calm3/Gm2a/H2-Q4/Pgrmc1/Atox1/Baiap2/Cd36/Map2k4/Map2k3/Glb1/Nkiras2/Psma1/Serpinb6a/Trem2/Prkaca/Rnaset2a/Kcmf1/Atp8a1/Lgmn/Galns/Rps6ka1/Ctsb/Slc44a2/B4galt1/Vav2/Lamp2/Cpped1/Mapkapk2/Actr10/Gns/Hgsnat/Gaa/Peli2/Psmc1/Arhgap9/Prkacb/Mapk3/Adgre5/Lamp1/Jun/Casp9/Hexb/Ptprc/C5ar1/Arsa/Alad/Bst2/Golga7/Manba/Dnase1l1/Mapkapk3/Rab3d/Txn |

| | | | | | | | | | | |
|---|---|---|---|---|---|---|---|---|---|---|
| | | | | | | | | | | 1/App/Neu1/Atp6v0d1/Pygb/Nckap1/Cstb/Jup/Prkce/Rnaset2b/Mlec/Ube2m/Rap1a/Arpc4/Casp8/Magt1/Atp6v1d/Ube2v1/Ikbkg/Atp6ap2/Fuca2/Irf5/Ubb/Tspan14/Trappc1/Ghdc/Dynlt1b/Fcgr1/Tmbim1/Psmb3/Slc11a1/Mif/Srp14/Tmem30a/Psmb2/Ckap4/Ctsl/Agl/Arpc2/Ecsit/Slc15a4/Psma6/Lyn/Panx1/Skp1/Clec4n/Rps6ka5/Psmb1/Rab6a/Rab5c/Uba52/Cd68/Elmo1/Mapk13/Ncstn/Cnpy3/Cyfip2/Atp6v1f/Nit2/Unc93b1/Pygl/Cyb5r3/Psmb6/Ddost/Vapa/Lamtor2/Pdxk/Cant1/Nme2/Wasf2/Ubc/Rab27a/Cap1/Atg7/Psmd3/Pik3r2/Mapk14/Mapk9/Hck/Cd300lb/Fbxw11/Psmb4/Gdi2/Psmb7/Limk1/Cpne3/Clec4e/Tax1bp1/Lpcat1/Ube2d2a/Ndufc2/Asah1/Vav3 |
| R-MMU-69278 | Cell Cycle, Mitotic | 457 | -0.560782633 | -1.910119009 | 5.59E-12 | 2.53E-09 | 2.31E-09 | 1940 | tags=26%, list=13%, signal=23% | Ppp2r1b/Psmd12/Ahctf1/Cdca8/Mcph1/Numa1/Mis12/Orc2/Cdk5rap2/Smc1a/Pold1/Pola2/Skp2/Nipbl/Pola1/Zw10/Espl1/Orc1/Ckap5/Rbm39/Ncapd3/Hsp90aa1/Cdk1/Ppp1r12a/Clasp1/Pcna/A730008H23Rik/Nup98/Ncapd2/Kpnb1/Nup133/Anapc1/Akt3/Esco1/Pds5b/Mcm8/Tubb5/Cdca5/Dynll1/Nup155/Sfi1/Tubgcp4/Lig1/Taok1/Cdc25b/Ranbp2/Mcm6/Ccnb1/Akap9/Mcm4/Cdt1/Pcnt/Rfc5/Tpr/Cnep1r1/Spdl1/Ncapg/Tuba1a/Pcm1/Lmnb1/Cks1b/Ccnd1/Cenpa/Tpx2/Cep152/Cpap/Nup107/Rcc1/Mcm3/Anapc4/Cep192/Tubb2a/Dync1h1/Mcm10/Cdk2/Tubb2b/Cep135/Cenph/Plk1/Spc24/Tubb3/Ncapg2/Smc2/Sgo2a/Pole2/Aurka/Esco2/Bub1/Kif18a/Fbxo5/Bub1b/Pmf1/Cep290/Cenpe/Kif23/Ube2c/Aurkb/Kif2c/Ccna2/Nuf2/Nek2/Mcm5/Ercc6l/Cdc45/Pole/Cdc20/Pkmyt1/Cdc6/Ndc80/Sgo1/Hmmr/E2f2/Ccnb2/Kntc1/Kif20a/Cenpu/Cenpf |
| R-MMU-6798695 | Neutrophil degranulation | 397 | 0.564943585 | 2.036978265 | 3.09E-11 | 1.05E-08 | 9.57E-09 | 1839 | tags=28%, list=12%, signal=25% | H2-Q7/Crispld2/Pirb/Nfam1/Mndal/Cybb/Cxcl1/Ifi211/Itgb2/Cfp/Fcer1g/Lyz2/Ctss/B2m/C3ar1/Arg1/Cd93/H2-K1/Nckap1l/Tnfrsf1b/Tyrobp/Fth1/Cd53/Clec4d/Pycard/Qsox1/Mmp8/Adam8/Cyba/H2-T22/Ctsh/Cst3/H2-T23/Fuca1/Gyg1/Ctsd/Man2b1/Aga/Prcp/Aprt/Gstp2/Arl8a/Padi2/Cat/Npc2/Cd14/Fabp5/Hk3/Psap/Ostf1/Slpi/Creg1/Stom/Gsn/Grn/Gm2a/H2-Q4/Pgrmc1/Cd36/Glb1/Serpinb6a/Rnaset2a/Kcmf1/Atp8a1/Galns/Ctsb/Slc44a2/B4 |

| | | | | | | | | | | |
|---|---|---|---|---|---|---|---|---|---|---|
| | | | | | | | | | | galt1/Lamp2/Cpped1/Actr10/Gns/Hgsnat/Gaa/Arhgap9/Adgre5/Lamp1/Hexb/Ptprc/C5ar1/Arsa/Alad/Bst2/Golga7/Manba/Dnase1l1/Rab3d/Neu1/Pygb/Cstb/Jup/Rnaset2b/Mlec/Rap1a/Magt1/Atp6v1d/Atp6ap2/Fuca2/Tspan14/Trappc1/Ghdc/Dynlt1b/Tmbim1/Slc11a1/Mif/Srp14/Tmem30a/Ckap4/Agl/Slc15a4 |
| R-MMU-69620 | Cell Cycle Checkpoints | 245 | -0.622979669 | -2.005523962 | 9.06E-11 | 2.47E-08 | 2.24E-08 | 1454 | tags=25%, list=10%, signal=23% | Orc1/Ckap5/Cdk1/Clasp1/Topbp1/Nup98/Nup133/Anapc1/Mcm8/Dynll1/Taok1/Ranbp2/Mcm6/Ccnb1/Trp53bp1/Mcm4/Mdc1/Rfc5/Spdl1/Bard1/Nsd2/Brca1/Cenpa/Nup107/Herc2/Chek1/Mcm3/Anapc4/Dync1h1/Mcm10/Cdk2/Brip1/Cenph/Plk1/Spc24/Sgo2a/Bub1/Kif18a/Bub1b/Pmf1/Cenpe/Mdm4/Ube2c/Aurkb/Kif2c/Ccna2/Nuf2/Mcm5/Ercc6l/Cdc45/Cdc20/Pkmyt1/Cdc6/Ndc80/Sgo1/Clspn/Exo1/Ccnb2/Kntc1/Cenpu/Cenpf |
| R-MMU-68877 | Mitotic Prometaphase | 182 | -0.647924048 | -2.035219919 | 4.83E-10 | 1.09E-07 | 9.96E-08 | 2150 | tags=38%, list=14%, signal=33% | Dync1li2/Cenpn/Tubg2/Dync1li1/Alms1/Ska1/Ppp2r1b/Ahctf1/Cdca8/Numa1/Mis12/Cdk5rap2/Smc1a/Zw10/Ckap5/Hsp90aa1/Cdk1/Clasp1/Nup98/Ncapd2/Nup133/Pds5b/Tubb5/Cdca5/Dynll1/Sfi1/Tubgcp4/Taok1/Ranbp2/Ccnb1/Akap9/Pcnt/Spdl1/Ncapg/Tuba1a/Pcm1/Cenpa/Cep152/Cpap/Nup107/Cep192/Tubb2a/Dync1h1/Tubb2b/Cep135/Cenph/Plk1/Spc24/Tubb3/Smc2/Sgo2a/Bub1/Kif18a/Bub1b/Pmf1/Cep290/Cenpe/Aurkb/Kif2c/Nuf2/Nek2/Ercc6l/Cdc20/Ndc80/Sgo1/Ccnb2/Kntc1/Cenpu/Cenpf |
| R-MMU-2500257 | Resolution of Sister Chromatid Cohesion | 113 | -0.701176545 | -2.077298703 | 1.54E-09 | 2.98E-07 | 2.71E-07 | 2150 | tags=43%, list=14%, signal=37% | Dync1li2/Cenpn/Dync1li1/Ska1/Ppp2r1b/Ahctf1/Cdca8/Mis12/Smc1a/Zw10/Ckap5/Cdk1/Clasp1/Nup98/Nup133/Pds5b/Cdca5/Dynll1/Taok1/Ranbp2/Ccnb1/Spdl1/Tuba1a/Cenpa/Nup107/Tubb2a/Dync1h1/Tubb2b/Cenph/Plk1/Spc24/Tubb3/Sgo2a/Bub1/Kif18a/Bub1b/Pmf1/Cenpe/Aurkb/Kif2c/Nuf2/Ercc6l/Cdc20/Ndc80/Sgo1/Ccnb2/Kntc1/Cenpu/Cenpf |
| R-MMU-9648025 | EML4 and NUDC in mitotic spindle formation | 101 | -0.700822323 | -2.04921139 | 4.85E-08 | 7.33E-06 | 6.67E-06 | 1320 | tags=43%, list=9%, signal=39% | Dync1li2/Cenpn/Dync1li1/Ska1/Ppp2r1b/Ahctf1/Cdca8/Mis12/Zw10/Ckap5/Clasp1/Nup98/Nup133/Dynll1/Taok1/Ranbp2/Spdl1/Tuba1a/Cenpa/Nup107/Tubb2a/Dync1h1/Tubb2b/Cenph/Plk1/Spc24/Tubb3/Sgo2a/Bub1/Kif18a/Bub1b/Pmf1/Cenpe/Aurkb/Kif2c/Nuf2/Ercc6l/Cdc20/Ndc80/Sgo1/Kntc1/Cenpu/Cenpf |

| | | | | | | | | | | |
|---|---|---|---|---|---|---|---|---|---|---|
| R-MMU-2555396 | Mitotic Metaphase and Anaphase | 200 | -0.605306445 | -1.912014381 | 4.43E-08 | 7.33E-06 | 6.67E-06 | 1543 | tags=25%, list=10%, signal=23% | Zw10/Espl1/Ckap5/Rbm39/Cdk1/Clasp1/Nup98/Kpnb1/Nup133/Anapc1/Pds5b/Cdca5/Dynll1/Nup155/Taok1/Ranbp2/Ccnb1/Spdl1/Tuba1a/Lmnb1/Cenpa/Nup107/Rcc1/Anapc4/Tubb2a/Dync1h1/Tubb2b/Cenph/Plk1/Spc24/Tubb3/Sgo2a/Bub1/Kif18a/Fbxo5/Bub1b/Pmf1/Cenpe/Ube2c/Aurkb/Kif2c/Nuf2/Ercc6l/Cdc20/Ndc80/Sgo1/Ccnb2/Kntc1/Cenpu/Cenpf |
| R-MMU-68882 | Mitotic Anaphase | 199 | -0.600154113 | -1.895285373 | 6.14E-08 | 8.35E-06 | 7.60E-06 | 1543 | tags=25%, list=10%, signal=22% | Zw10/Espl1/Ckap5/Rbm39/Cdk1/Clasp1/Nup98/Kpnb1/Nup133/Anapc1/Pds5b/Cdca5/Dynll1/Nup155/Taok1/Ranbp2/Ccnb1/Spdl1/Tuba1a/Lmnb1/Cenpa/Nup107/Rcc1/Anapc4/Tubb2a/Dync1h1/Tubb2b/Cenph/Plk1/Spc24/Tubb3/Sgo2a/Bub1/Kif18a/Bub1b/Pmf1/Cenpe/Ube2c/Aurkb/Kif2c/Nuf2/Ercc6l/Cdc20/Ndc80/Sgo1/Ccnb2/Kntc1/Cenpu/Cenpf |
| R-MMU-5663220 | RHO GTPases Activate Formins | 126 | -0.662364339 | -2.000311454 | 7.10E-08 | 8.78E-06 | 7.99E-06 | 2235 | tags=38%, list=15%, signal=33% | Dvl2/Actb/Dync1li2/Cenpn/Dync1li1/Daam1/Ska1/Ppp2r1b/Ahctf1/Cdca8/Mis12/Dvl1/Zw10/Scai/Ckap5/Clasp1/Nup98/Nup133/Dynll1/Taok1/Ranbp2/Spdl1/Tuba1a/Cenpa/Nup107/Tubb2a/Dync1h1/Tubb2b/Cenph/Plk1/Spc24/Tubb3/Sgo2a/Bub1/Kif18a/Bub1b/Pmf1/Cenpe/Aurkb/Kif2c/Nuf2/Ercc6l/Cdc20/Ndc80/Sgo1/Kntc1/Cenpu/Cenpf |
| R-MMU-141424 | Amplification of signal from the kinetochores | 89 | -0.704450091 | -2.010624506 | 9.27E-08 | 9.01E-06 | 8.20E-06 | 2150 | tags=44%, list=14%, signal=38% | Dync1li2/Cenpn/Dync1li1/Ska1/Ppp2r1b/Ahctf1/Cdca8/Mis12/Zw10/Ckap5/Clasp1/Nup98/Nup133/Dynll1/Taok1/Ranbp2/Spdl1/Cenpa/Nup107/Dync1h1/Cenph/Plk1/Spc24/Sgo2a/Bub1/Kif18a/Bub1b/Pmf1/Cenpe/Aurkb/Kif2c/Nuf2/Ercc6l/Cdc20/Ndc80/Sgo1/Kntc1/Cenpu/Cenpf |
| R-MMU-141444 | Amplification of signal from unattached kinetochores via a MAD2 inhibitory signal | 89 | -0.704450091 | -2.010624506 | 9.27E-08 | 9.01E-06 | 8.20E-06 | 2150 | tags=44%, list=14%, signal=38% | Dync1li2/Cenpn/Dync1li1/Ska1/Ppp2r1b/Ahctf1/Cdca8/Mis12/Zw10/Ckap5/Clasp1/Nup98/Nup133/Dynll1/Taok1/Ranbp2/Spdl1/Cenpa/Nup107/Dync1h1/Cenph/Plk1/Spc24/Sgo2a/Bub1/Kif18a/Bub1b/Pmf1/Cenpe/Aurkb/Kif2c/Nuf2/Ercc6l/Cdc20/Ndc80/Sgo1/Kntc1/Cenpu/Cenpf |
| R-MMU-68886 | M Phase | 337 | -0.541656341 | -1.789123374 | 8.99E-08 | 9.01E-06 | 8.20E-06 | 1580 | tags=22%, list=10%, signal=20% | Nipbl/Zw10/Espl1/Ckap5/Rbm39/Ncapd3/Hsp90aa1/Cdk1/Clasp1/Nup98/Ncapd2/Kpnb1/Nup133/Anapc1/Pds5b/Tubb5/Cdca5/Dynll1/Nup155/Sfi1/Tubgcp4/Taok1/Ranbp2/Ccnb1/Akap9/Pcnt/Tpr/Cnep1r1/Spdl1/Ncapg/Tuba1a/Pcm1/Lmnb1/Cenpa/Cep152/Cpap/Nup107/Rcc1/Anapc4/Cep192/Tubb2a/Dync1h1/Tubb2b/Cep135/Cenph/Plk1/Spc24/Tubb3/Ncapg2/Smc2/S |

| | | | | | | | | | | |
|---|---|---|---|---|---|---|---|---|---|---|
| | | | | | | | | | | go2a/Bub1/Kif18a/Fbxo5/Bub1b/Pmf1/Cep290/Cenpe/Kif23/Ube2c/Aurkb/Kif2c/Nuf2/Nek2/Ercc6l/Cdc20/Ndc80/Sgo1/Ccnb2/Kntc1/Kif20a/Cenpu/Cenpf |
| R-MMU-69618 | Mitotic Spindle Checkpoint | 106 | -0.680308334 | -1.999081592 | 1.85E-07 | 1.68E-05 | 1.53E-05 | 1320 | tags=41%, list=9%, signal=37% | Cdc27/Dync1li2/Cenpn/Dync1li1/Ska1/Ppp2r1b/Ahctf1/Cdca8/Mis12/Zw10/Ckap5/Clasp1/Nup98/Nup133/Anapc1/Dynll1/Taok1/Ranbp2/Spdl1/Cenpa/Nup107/Anapc4/Dync1h1/Cenph/Plk1/Spc24/Sgo2a/Bub1/Kif18a/Bub1b/Pmf1/Cenpe/Ube2c/Aurkb/Kif2c/Nuf2/Ercc6l/Cdc20/Ndc80/Sgo1/Kntc1/Cenpu/Cenpf |
| R-MMU-2467813 | Separation of Sister Chromatids | 163 | -0.615714204 | -1.906583641 | 3.18E-07 | 2.70E-05 | 2.46E-05 | 2926 | tags=42%, list=19%, signal=34% | Nup160/Xpo1/Wapl/Rcc2/Stag1/Cenpo/Anapc10/Ppp1cc/Nde1/Nup85/Cenpq/Tuba1b/Anapc2/Psmc6/Stag2/Incenp/Kif2a/Cdc27/Dync1li2/Cenpn/Dync1li1/Ska1/Ppp2r1b/Psmd12/Ahctf1/Cdca8/Mis12/Smc1a/Zw10/Espl1/Ckap5/Clasp1/Nup98/Nup133/Anapc1/Pds5b/Cdca5/Dynll1/Taok1/Ranbp2/Spdl1/Tuba1a/Cenpa/Nup107/Anapc4/Tubb2a/Dync1h1/Tubb2b/Cenph/Plk1/Spc24/Tubb3/Sgo2a/Bub1/Kif18a/Bub1b/Pmf1/Cenpe/Ube2c/Aurkb/Kif2c/Nuf2/Ercc6l/Cdc20/Ndc80/Sgo1/Kntc1/Cenpu/Cenpf |
| R-MMU-983189 | Kinesins | 45 | -0.787500142 | -2.065635194 | 6.73E-07 | 5.39E-05 | 4.90E-05 | 967 | tags=38%, list=6%, signal=35% | Kif1b/Kif18b/Tuba1a/Kif22/Tubb2a/Kif21b/Tubb2b/Tubb3/Kif18a/Cenpe/Kif23/Racgap1/Kif2c/Kif4/Kif11/Kif20b/Kif20a |
| R-MMU-195258 | RHO GTPase Effectors | 226 | -0.54887414 | -1.748658673 | 3.28E-06 | 0.000247655 | 0.000225419 | 1545 | tags=23%, list=10%, signal=21% | Dvl1/Zw10/Scai/Rock1/Rtkn/Ckap5/Calm2/Ppp1r12a/Calm1/Clasp1/Nup98/Nup133/Myh9/Myh14/Abi1/Actr3/Dynll1/Taok1/Ranbp2/Prkcz/Spdl1/Tuba1a/Mapk11/Prc1/Cenpa/Nup107/Tubb2a/Dync1h1/Tubb2b/Cenph/Plk1/Spc24/Tubb3/Sgo2a/Bub1/Kif18a/Iqgap3/Bub1b/Pmf1/Cenpe/Aurkb/Kif2c/Nuf2/Myh10/Ercc6l/Cdc20/Ndc80/Sgo1/Kntc1/Kif14/Cenpu/Cit/Cenpf |
| R-MMU-73894 | DNA Repair | 276 | -0.525795367 | -1.709474108 | 3.62E-06 | 0.000259319 | 0.000236036 | 2988 | tags=34%, list=20%, signal=28% | Poln/Tfpt/Rpa2/Mre11a/Abl1/Actr8/Cops5/Slx4/Ddb1/Rif1/Ruvbl1/Firrm/Gtf2h3/Rev1/Dclre1c/Chd1l/Gtf2h1/Fancl/Pias4/Polm/Ufd1/Ercc4/Polr2d/Mbd4/Rad18/Rmi2/Pole4/Actb/Usp10/Usp1/Kdm4a/Rnf111/Trp53/Usp7/Fancg/Baz1b/Sirt6/Apbb1/Msh3/Blm/Pold1/Apex1/Eme1/Polr2a/Pot1a/Aqr/Ercc2/Fignl1/Cops8/Dtl/Tdg/Topbp1/Pcna/Polr2b/Smarca5/Terf2/Isg15/Mapk8/Rev3l/Lig4/Lig1/Parp2/Xrcc5/Trp53bp1/Mdc1/Rfc5/Nfrkb/Timeless/Rad23a/Bard1/Nsd2/Ino80/Brca1/Herc2/Chek1/Uvssa/Prkdc/Ino80d/Fancm/Cdk2/Brip1/Pole2/Fan |

| | | | | | | | | | | |
|---|---|---|---|---|---|---|---|---|---|---|
| | | | | | | | | | | ca/Polq/Neil3/Ccna2/Rad51ap1/Fanci/Pole/Fancd2/Rad51/Clspn/Exo1/Gen1/Pclaf |
| R-MMU-166658 | Complement cascade | 20 | 0.887042092 | 2.043856282 | 4.80E-06 | 0.000326228 | 0.000296938 | 542 | tags=40%, list=4%, signal=39% | Serping1/C1qc/C1qa/C1qb/Cfp/C3ar1/C2/Cd81 |
| R-MMU-9716542 | Signaling by Rho GTPases, Miro GTPases and RHOBTB3 | 557 | -0.4519696 | -1.558749941 | 8.46E-06 | 0.0005481 | 0.00049889 | 2317 | tags=25%, list=15%, signal=22% | Actn1/Arhgef1/Ptpn13/Sptbn1/Frs3/Dvl2/Actb/Plxna1/Bcr/Cdc25c/Map3k11/Arhgap33/Dync1li2/Cenpn/Vangl2/Dync1li1/Tor1aip1/Daam1/Ska1/Usp9x/Senp1/Ppp2r1b/Ahctf1/Cdca8/Fgd4/Mis12/Arhgap29/Net1/Itsn1/Ankfy1/Rac2/Tjp2/Rhot2/Cct6a/Erbin/Frs2/Epha2/Dvl1/Zw10/Sema4f/Scai/Tra2b/Rock1/Rtkn/Ckap5/Myo9a/Rbm39/Hsp90aa1/Calm2/Rbbp6/Rapgef1/Ppp1r12a/Calm1/Ddx39b/Clasp1/Fgd1/Nup98/Nup133/Pcdh7/Garre1/Myh9/Myh14/Abi1/Cdc42bpb/Golga3/Fam13b/Arhgap30/Rnf20/Actr3/Arhgef25/Arhgef10/Dynll1/Arhgef18/Myo6/Rhobtb2/Peak1/Arhgap32/Arhgap31/Mfn1/Depdc1b/Taok1/Tfrc/Ranbp2/Uaca/Prkcz/Rhobtb1/Prex2/Ckb/Trak2/Cpsf7/Spdl1/Tuba1a/Mapk11/Trio/Farp2/Prc1/Lmnb1/Nisch/Plxnd1/Cenpa/Nup107/Shmt2/Rbmx/Tubb2a/Akap13/Dync1h1/Dst/Tubb2b/Cenph/Plk1/Spc24/Tubb3/Sgo2a/Anln/Bub1/Kif18a/Dock5/Iqgap3/Bub1b/Pmf1/Cenpe/Stard13/Aurkb/Racgap1/Kif2c/Nuf2/Myh10/Ercc6l/Cdc20/Ndc80/Arhgef7/Sgo1/Kntc1/Arhgef39/Kif14/Cenpu/Cit/Cenpf |
| R-MMU-194315 | Signaling by Rho GTPases | 543 | -0.455418193 | -1.568398382 | 9.80E-06 | 0.000605544 | 0.000551177 | 2317 | tags=25%, list=15%, signal=22% | Actn1/Arhgef1/Ptpn13/Sptbn1/Frs3/Dvl2/Actb/Plxna1/Bcr/Cdc25c/Map3k11/Arhgap33/Dync1li2/Cenpn/Vangl2/Dync1li1/Tor1aip1/Daam1/Ska1/Usp9x/Senp1/Ppp2r1b/Ahctf1/Cdca8/Fgd4/Mis12/Arhgap29/Net1/Itsn1/Ankfy1/Rac2/Tjp2/Cct6a/Erbin/Frs2/Epha2/Dvl1/Zw10/Sema4f/Scai/Tra2b/Rock1/Rtkn/Ckap5/Myo9a/Rbm39/Hsp90aa1/Calm2/Rbbp6/Rapgef1/Ppp1r12a/Calm1/Ddx39b/Clasp1/Fgd1/Nup98/Nup133/Pcdh7/Garre1/Myh9/Myh14/Abi1/Cdc42bpb/Golga3/Fam13b/Arhgap30/Rnf20/Actr3/Arhgef25/Arhgef10/Dynll1/Arhgef18/Myo6/Rhobtb2/Peak1/Arhgap32/Arhgap31/Depdc1b/Taok1/Tfrc/Ranbp2/Uaca/Prkcz/Rhobtb1/Prex2/Ckb/Cpsf7/Spdl1/Tuba1a/Mapk11/Trio/Farp2/Prc1/Lmnb1/Nisch/Plxnd1/Cenpa/Nup107/Shmt2/Rbmx/Tubb2a/Akap13/Dync1h1/Dst/Tubb2b/Cenph/Plk1/Spc24/Tubb3/Sgo2a/Anln/Bub1/Kif |

| | | | | | | | | | | |
|---|---|---|---|---|---|---|---|---|---|---|
| | | | | | | | | | | 18a/Dock5/Iqgap3/Bub1b/Pmf1/Cenpe/Stard13/Aurkb/Racgap1/Kif2c/Nuf2/Myh10/Ercc6l/Cdc20/Ndc80/Arhgef7/Sgo1/Kntc1/Arhgef39/Kif14/Cenpu/Cit/Cenpf |
| R-MMU-977606 | Regulation of Complement cascade | 18 | 0.893034667 | 2.006429219 | 1.21E-05 | 0.000715097 | 0.000650894 | 542 | tags=44%, list=4%, signal=43% | Serping1/C1qc/C1qa/C1qb/Cfp/C3ar1/C2/Cd81 |
| R-MMU-2172127 | DAP12 interactions | 38 | 0.804134497 | 2.112629264 | 1.74E-05 | 0.000977723 | 0.00088994 | 978 | tags=26%, list=6%, signal=25% | H2-Q7/B2m/H2-K1/Tyrobp/H2-T22/H2-T23/Hras/H2-Q4/Trem2/Vav2 |
| R-MMU-380108 | Chemokine receptors bind chemokines | 23 | 0.863814836 | 2.01443434 | 1.80E-05 | 0.000977723 | 0.00088994 | 92 | tags=17%, list=1%, signal=17% | Cxcl12/Cxcl1/Pf4/Ackr3 |
| R-MMU-1236977 | Endosomal/Vacuolar pathway | 13 | 0.906518534 | 1.898181193 | 4.17E-05 | 0.002102669 | 0.001913884 | 688 | tags=46%, list=5%, signal=44% | H2-Q7/B2m/H2-K1/H2-T22/H2-T23/H2-Q4 |
| R-MMU-173623 | Classical antibody-mediated complement activation | 5 | 0.981575323 | 1.707395552 | 4.05E-05 | 0.002102669 | 0.001913884 | 44 | tags=60%, list=0%, signal=60% | C1qc/C1qa/C1qb |
| R-MMU-72203 | Processing of Capped Intron-Containing Pre-mRNA | 255 | -0.514058149 | -1.65716572 | 4.66E-05 | 0.002264538 | 0.00206122 | 3050 | tags=36%, list=20%, signal=29% | Sympk/Nup88/Nup54/Prpf31/Lsm8/Nup160/Rbm5/Pqbp1/Ppwd1/Sart1/Rbm10/Lsm5/Cpsf1/Nup85/Hnrnph2/Htatsf1/Ddx23/Eftud2/Ddx42/Dhx16/Tcerg1/Wbp4/Nup42/Cstf3/Nup214/Polr2d/Snrpf/Nup35/Papola/Zc3h11a/Slbp/Cstf1/Dhx15/Mettl3/Cdc40/Wbp11/Snrnp70/Prpf4b/Pabpn1/Rbm8a/Cwc22/Snrnp48/Polr2a/Ncbp1/Prpf40a/Aqr/Cpsf6/Tra2b/Rbm39/Sarnp/Lsm6/Ddx39b/Hnrnpu/Ppil1/Nup98/Nup133/Pcf11/Polr2b/Nxf1/Nup155/Acin1/Luc7l3/U2af2/Srsf10/Ddx46/Cstf2/Mtrex/Ranbp2/Tpr/Cpsf7/Thoc2/Srsf1/Srsf2/Pnn/Prpf3/Hnrnpa2b1/Fus/Srsf11/Ddx39a/Srsf7/Nup107/Hnrnpa1/Hnrnph1/Dhx9/Fyttd1/Rbmx/Rbm25/Srsf3/Sf3b3/Ppil4/Srrm2/Srrt |
| R-MMU-1236975 | Antigen processing-Cross presentation | 70 | 0.680385071 | 1.986595171 | 9.20E-05 | 0.004313248 | 0.003925991 | 2486 | tags=39%, list=16%, signal=32% | H2-Q7/Cybb/Psmb9/B2m/Ncf1/H2-K1/Tap1/Cyba/H2-T22/H2-T23/Tap2/H2-Q4/Cd36/Psma1/Psmc1/Itgb5/Psmb8/Fcgr1/Psmb3/Psmb2/Psma6/Psmb1/Psmb6/Psme1/Psmd3/Psmb4/Psmb7 |
| R-MMU-69273 | Cyclin A/B1/B2 associated events during G2/M transition | 29 | -0.767794407 | -1.840438622 | 9.92E-05 | 0.004498825 | 0.004094906 | 1354 | tags=34%, list=9%, signal=31% | Cdk1/A730008H23Rik/Cdc25b/Ccnb1/Cdk2/Plk1/Ccna2/Pkmyt1/Sgo1/Ccnb2 |
| R-MMU-166786 | Creation of C4 and C2 activators | 6 | 0.969124352 | 1.754914079 | 0.000117982 | 0.005176 | 0.004711282 | 44 | tags=50%, list=0%, signal=50% | C1qc/C1qa/C1qb |

| | | | | | | | | | | |
|---|---|---|---|---|---|---|---|---|---|---|
| R-MMU-69481 | G2/M Checkpoints | 129 | -0.572002215 | -1.725985412 | 0.000122098 | 0.005189185 | 0.004723283 | 895 | tags=21%, list=6%, signal=20% | Orc1/Cdk1/Topbp1/Mcm8/Mcm6/Ccnb1/Trp53bp1/Mcm4/Mdc1/Rfc5/Bard1/Nsd2/Brca1/Herc2/Chek1/Mcm3/Mcm10/Cdk2/Brip1/Ccna2/Mcm5/Cdc45/Pkmyt1/Cdc6/Clspn/Exo1/Ccnb2 |
| R-MMU-1799339 | SRP-dependent cotranslational protein targeting to membrane | 89 | 0.629359165 | 1.922907166 | 0.00021264 | 0.00826298 | 0.00752075 5 | 2850 | tags=64%, list=19%, signal=52% | Rps19/Rps27rt/Rps15/Rps5/Rps27l/Rpl15/Rps14/Rpl19/Rpl28/Rpl18a/Rpl9/Rpl37/Rps11/Rpl32/Rps17/Rps21/Rpl29/Rps13/Rps28/Rpl24/Rpl38/Rps10/Rpl7a/Rps7/Rpl26/Rpl36/Rpl23a/Rplp2/Srp14/Rps2/Rpl13a/Rps16/Rplp0/Rpl39/Rpl18/Rpl13/Rpl14/Uba52/Rpl6/Rpl35a/Rps18/Rps24/Rps20/Rps29/Rpl22/Rps12/Rps25/Rpl34/Fau/Rps4x/Rpl17/Rpl3/Rps6/Rps27/Rpl27/Rpl37a/Rpl7 |
| R-MMU-72689 | Formation of a pool of free 40S subunits | 98 | 0.616837016 | 1.913284545 | 0.000209277 | 0.008262598 | 0.007520755 | 2850 | tags=62%, list=19%, signal=51% | Rps19/Rps27rt/Rps15/Rps5/Rps27l/Eif3f/Rpl15/Rps14/Rpl19/Rpl28/Rpl18a/Rpl9/Rpl37/Rps11/Rpl32/Rps17/Rps21/Eif3k/Rpl29/Rps13/Eif3h/Rps28/Rpl24/Rpl38/Rps10/Rpl7a/Rps7/Rpl26/Rpl36/Rpl23a/Rplp2/Eif3d/Rps2/Rpl13a/Rps16/Rplp0/Rpl39/Rpl18/Rpl13/Rpl14/Uba52/Rpl6/Rpl35a/Rps18/Rps24/Rps20/Rps29/Rpl22/Rps12/Rps25/Rpl34/Fau/Rps4x/Rpl17/Eif1ax/Rpl3/Rps6/Rps27/Rpl27/Rpl37a/Rpl7 |
| R-MMU-5693538 | Homology Directed Repair | 108 | -0.591487912 | -1.739645057 | 0.000203217 | 0.008262598 | 0.007520755 | 1688 | tags=25%, list=11%, signal=22% | Blm/Pold1/Eme1/Fignl1/Topbp1/Pcna/Parp2/Trp53bp1/Mdc1/Rfc5/Timeless/Bard1/Nsd2/Brca1/Herc2/Chek1/Cdk2/Brip1/Pole2/Polq/Ccna2/Rad51ap1/Pole/Rad51/Clspn/Exo1/Gen1 |
| R-MMU-5693532 | DNA Double-Strand Break Repair | 136 | -0.557276222 | -1.699689642 | 0.000225261 | 0.008509846 | 0.007745804 | 2428 | tags=31%, list=16%, signal=26% | Pias4/Polm/Ercc4/Rmi2/Pole4/Kdm4a/Trp53/Baz1b/Sirt6/Apbb1/Blm/Pold1/Eme1/Fignl1/Topbp1/Pcna/Smarca5/Mapk8/Lig4/Parp2/Xrcc5/Trp53bp1/Mdc1/Rfc5/Timeless/Bard1/Nsd2/Brca1/Herc2/Chek1/Prkdc/Cdk2/Brip1/Pole2/Polq/Ccna2/Rad51ap1/Pole/Rad51/Clspn/Exo1/Gen1 |
| R-MMU-69275 | G2/M Transition | 162 | -0.540354865 | -1.674002288 | 0.000241118 | 0.008862701 | 0.008066978 | 1448 | tags=22%, list=10%, signal=20% | Ckap5/Hsp90aa1/Cdk1/Ppp1r12a/Clasp1/A730008H23Rik/Tubb5/Dynll1/Sfi1/Tubgcp4/Cdc25b/Ccnb1/Akap9/Pcnt/Tuba1a/Pcm1/Tpx2/Cep152/Cpap/Cep192/Tubb2a/Dync1h1/Cdk2/Tubb2b/Cep135/Plk1/Tubb3/Aurka/Cep290/Ccna2/Nek2/Pkmyt1/Sgo1/Hmmr/Ccnb2 |
| R-MMU-166663 | Initial triggering of complement | 11 | 0.903152603 | 1.881836354 | 0.000249701 | 0.008936665 | 0.008134302 | 394 | tags=45%, list=3%, signal=44% | C1qc/C1qa/C1qb/Cfp/C2 |
| R-MMU-198933 | Immunoregulatory interactions between a | 54 | 0.713932962 | 2.005692863 | 0.000260048 | 0.009068323 | 0.00825414 | 861 | tags=28%, list=6%, signal=26% | H2-Q7/Siglec1/Itgb2/B2m/H2-K1/Col3a1/Tyrobp/H2-T22/H2- |

| | | | | | | | | | | |
|---|---|---|---|---|---|---|---|---|---|---|
| | Lymphoid and a non-Lymphoid cell | | | | | | | | | T23/Col1a2/Cd81/Nectin2/H2-Q4/Icam5/Trem2 |
| R-MMU-1236974 | ER-Phagosome pathway | 38 | 0.747062809 | 1.962690019 | 0.000336519 | 0.011441635 | 0.010414368 | 838 | tags=26%, list=6%, signal=25% | H2-Q7/Psmb9/B2m/H2-K1/Tap1/H2-T22/H2-T23/Tap2/H2-Q4/Psma1 |
| R-MMU-983170 | Antigen Presentation: Folding, assembly and peptide loading of class I MHC | 29 | 0.76780962 | 1.867510939 | 0.000379166 | 0.011565978 | 0.010527547 | 880 | tags=31%, list=6%, signal=29% | H2-Q7/B2m/H2-K1/Tap1/H2-T22/H2-T23/Tap2/H2-Q4/Bcap31 |
| R-MMU-72187 | mRNA 3'-end processing | 54 | -0.676451837 | -1.819694377 | 0.000382698 | 0.011565978 | 0.010527547 | 1975 | tags=41%, list=13%, signal=36% | Papola/Zc3h11a/Cstf1/Cdc40/Pabpn1/Rbm8a/Ncbp1/Cpsf6/Sarnp/Ddx39b/Pcf11/U2af2/Cstf2/Cpsf7/Thoc2/Srsf1/Srsf2/Srsf11/Ddx39a/Srsf7/Fyttd1/Srsf3 |
| R-MMU-8854518 | AURKA Activation by TPX2 | 68 | -0.63928237 | -1.772415886 | 0.000351851 | 0.011565978 | 0.010527547 | 1448 | tags=32%, list=10%, signal=29% | Ckap5/Hsp90aa1/Cdk1/Clasp1/Tubb5/Dynll1/Sfi1/Akap9/Pcnt/Tuba1a/Pcm1/Tpx2/Cep152/Cpap/Cep192/Dync1h1/Cep135/Plk1/Aurka/Cep290/Nek2/Hmmr |
| R-MMU-983231 | Factors involved in megakaryocyte development and platelet production | 104 | -0.587231218 | -1.722741028 | 0.000371707 | 0.011565978 | 0.010527547 | 996 | tags=20%, list=7%, signal=19% | Mfn1/Kif1b/Kif18b/Jmjd1c/Tuba1a/Cbx5/Kif22/Tubb2a/Kif21b/Tubb2b/Tubb3/Kif18a/Dock5/Cenpe/Kif23/Racgap1/Kif2c/Kif4/Kif11/Kif20b/Kif20a |
| R-MMU-453274 | Mitotic G2-G2/M phases | 164 | -0.535192308 | -1.656350269 | 0.000376173 | 0.011565978 | 0.010527547 | 1448 | tags=21%, list=10%, signal=20% | Ckap5/Hsp90aa1/Cdk1/Ppp1r12a/Clasp1/A730008H23Rik/Tubb5/Dynll1/Sfi1/Tubgcp4/Cdc25b/Ccnb1/Akap9/Pcnt/Tuba1a/Pcm1/Tpx2/Cep152/Cpap/Cep192/Tubb2a/Dync1h1/Cdk2/Tubb2b/Cep135/Plk1/Tubb3/Aurka/Cep290/Ccna2/Nek2/Pkmyt1/Sgo1/Hmmr/Ccnb2 |
| R-MMU-68962 | Activation of the pre-replicative complex | 31 | -0.750381035 | -1.821498445 | 0.000424962 | 0.011843054 | 0.010779746 | 1813 | tags=52%, list=12%, signal=45% | Orc2/Pola2/Pola1/Orc1/Mcm8/Mcm6/Mcm4/Cdt1/Mcm3/Mcm10/Cdk2/Pole2/Mcm5/Cdc45/Pole/Cdc6 |
| R-MMU-73856 | RNA Polymerase II Transcription Termination | 63 | -0.646072824 | -1.769724184 | 0.000426698 | 0.011843054 | 0.010779746 | 2322 | tags=41%, list=15%, signal=35% | Cstf3/Snrpf/Lsm11/Papola/Zc3h11a/Slbp/Cstf1/Cdc40/Pabpn1/Rbm8a/Ncbp1/Cpsf6/Sarnp/Ddx39b/Pcf11/U2af2/Cstf2/Cpsf7/Thoc2/Srsf1/Srsf2/Srsf11/Ddx39a/Srsf7/Fyttd1/Srsf3 |
| R-MMU-70635 | Urea cycle | 6 | 0.958274236 | 1.735266424 | 0.000405995 | 0.011843054 | 0.010779746 | 253 | tags=33%, list=2%, signal=33% | Arg1/Ass1 |
| R-MMU-72172 | mRNA Splicing | 189 | -0.499794063 | -1.573741416 | 0.000409697 | 0.011843054 | 0.010779746 | 2937 | tags=32%, list=19%, signal=26% | Prpf31/Lsm8/Rbm5/Pqbp1/Ppwd1/Sart1/Rbm10/Lsm5/Hnrnph2/Htatsf1/Ddx23/Eftud2/Ddx42/Dhx16/Tcerg1/Wbp4/Polr2d/Snrpf/Dhx15/Cdc40/Wbp11/Snrnp70/Prpf4b/Rbm8a/Cwc22/Snrnp48/Polr2a/Ncbp1/Prpf40a/Aqr/Tra2b/Rbm39/Lsm6/Ddx39b/Hnrnpu/Ppil1/Polr2b/Acin1/Luc7l3/U2af2/Srsf10/Ddx46/Mtrex/Srsf1/Srsf2/Pnn/Prpf3/Hnrnpa2b1/Fus/Srsf11/Srsf7/Hnrnpa1/ |

| | | | | | | | | | | |
|---|---|---|---|---|---|---|---|---|---|---|
| | | | | | | | | | | Hnrnph1/Dhx9/Rbmx/Rbm25/Srsf3/Sf3b3/Ppil4/Srrm2/Srrt |
| R-MMU-176187 | Activation of ATR in response to replication stress | 35 | -0.721986403 | -1.795550931 | 0.000533624 | 0.014514582 | 0.013211415 | 1454 | tags=37%, list=10%, signal=34% | Orc1/Mcm8/Mcm6/Mcm4/Rfc5/Chek1/Mcm3/Mcm10/Cdk2/Mcm5/Cdc45/Cdc6/Clspn |
| R-MMU-6811434 | COPI-dependent Golgi-to-ER retrograde traffic | 84 | -0.609214818 | -1.725253175 | 0.000641798 | 0.017114624 | 0.015578017 | 1012 | tags=29%, list=7%, signal=27% | Kif26b/Copz2/Copg1/Zw10/Kif13b/Kif15/Bnip1/Kif1b/Kif18b/Tuba1a/Kif22/Tubb2a/Kif21b/Tubb2b/Tubb3/Kif18a/Cenpe/Kif23/Racgap1/Kif2c/Kif4/Kif11/Kif20b/Kif20a |
| R-MMU-376176 | Signaling by ROBO receptors | 19 | 0.808530249 | 1.834963923 | 0.00066477 | 0.017386282 | 0.015825284 | 1077 | tags=37%, list=7%, signal=34% | Cxcl12/Slit3/Pfn2/Gpc1/Robo1/Slit2/Enah |
| R-MMU-975956 | Nonsense Mediated Decay (NMD) independent of the Exon Junction Complex (EJC) | 91 | 0.587985509 | 1.797209601 | 0.000770361 | 0.018380534 | 0.016730269 | 3308 | tags=67%, list=22%, signal=53% | Rps19/Rps27rt/Rps15/Rps5/Rps27l/Rpl15/Rps14/Rpl19/Rpl28/Rpl18a/Rpl9/Rpl37/Rps11/Rpl32/Rps17/Rps21/Rpl29/Rps13/Rps28/Rpl24/Rpl38/Rps10/Rpl7a/Rps7/Rpl26/Rpl36/Rpl23a/Rplp2/Rps2/Rpl13a/Rps16/Rplp0/Rpl39/Rpl18/Rpl13/Rpl14/Uba52/Rpl6/Rpl35a/Rps18/Rps24/Rps20/Rps29/Rpl22/Rps12/Rps25/Rpl34/Fau/Rps4x/Rpl17/Rpl3/Rps6/Rps27/Rpl27/Rpl37a/Rpl7/Rpl8/Rpl21/Ncbp2/Rplp1/Rpl4 |
| R-MMU-72706 | GTP hydrolysis and joining of the 60S ribosomal subunit | 109 | 0.565328514 | 1.782683016 | 0.000764328 | 0.018380534 | 0.016730269 | 3057 | tags=59%, list=20%, signal=47% | Rps19/Rps27rt/Rps15/Rps5/Rps27l/Eif3f/Rpl15/Rps14/Rpl19/Rpl28/Rpl18a/Rpl9/Rpl37/Rps11/Rpl32/Rps17/Rps21/Eif3k/Rpl29/Rps13/Eif3h/Rps28/Rpl24/Rpl38/Rps10/Rpl7a/Rps7/Rpl26/Rpl36/Rpl23a/Rplp2/Eif3d/Rps2/Rpl13a/Rps16/Rplp0/Rpl39/Rpl18/Rpl13/Rpl14/Uba52/Rpl6/Rpl35a/Rps18/Rps24/Rps20/Rps29/Rpl22/Rps12/Rps25/Rpl34/Fau/Rps4x/Rpl17/Eif1ax/Rpl3/Rps6/Rps27/Rpl27/Rpl37a/Rpl7/Eif4a2/Rpl8/Rpl21 |
| R-MMU-5693567 | HDR through Homologous Recombination (HRR) or Single Strand Annealing (SSA) | 105 | -0.57704106 | -1.697948798 | 0.000742386 | 0.018380534 | 0.016730269 | 1688 | tags=24%, list=11%, signal=21% | Blm/Pold1/Eme1/Fignl1/Topbp1/Pcna/Trp53bp1/Mdc1/Rfc5/Timeless/Bard1/Nsd2/Brca1/Herc2/Chek1/Cdk2/Brip1/Pole2/Ccna2/Rad51ap1/Pole/Rad51/Clspn/Exo1/Gen1 |
| R-MMU-2565942 | Regulation of PLK1 Activity at G2/M Transition | 81 | -0.595080545 | -1.675308467 | 0.000764009 | 0.018380534 | 0.016730269 | 1448 | tags=28%, list=10%, signal=26% | Ckap5/Hsp90aa1/Cdk1/Ppp1r12a/Clasp1/Tubb5/Dynll1/Sfi1/Ccnb1/Akap9/Pcnt/Tuba1a/Pcm1/Cep152/Cpap/Cep192/Dync1h1/Cep135/Plk1/Aurka/Cep290/Nek2/Ccnb2 |
| R-MMU-72163 | mRNA Splicing - Major Pathway | 183 | -0.509747729 | -1.600655019 | 0.000736838 | 0.018380534 | 0.016730269 | 2937 | tags=33%, list=19%, signal=27% | Prpf31/Lsm8/Rbm5/Pqbp1/Ppwd1/Sart1/Rbm10/Lsm5/Hnrnph2/Htatsf1/Ddx23/Eftud2/Ddx42/Dhx16/Tcerg1/Wbp4/Polr2d/Snrpf/Dhx15/Cdc40/Wbp11/Snrnp70/Prpf4b/Rbm8a/Cwc22/Polr2a/Ncbp1/Prpf40a/Aqr/Tra2b/Rbm39/Lsm6/Ddx39b/Hnrnpu/P |

| | | | | | | | | | | |
|---|---|---|---|---|---|---|---|---|---|---|
| | | | | | | | | | | pil1/Polr2b/Acin1/Luc7l3/U2af2/Srsf10/Ddx46/Mtrex/Srsf1/Srsf2/Pnn/Prpf3/Hnrnpa2b1/Fus/Srsf11/Srsf7/Hnrnpa1/Hnrnph1/Dhx9/Rbmx/Rbm25/Srsf3/Sf3b3/Ppil4/Srrm2/Srrt |
| R-MMU-6804756 | Regulation of TP53 Activity through Phosphorylation | 82 | -0.596477816 | -1.681161554 | 0.000832792 | 0.019527546 | 0.017774299 | 1306 | tags=28%, list=9%, signal=26% | Blm/Prkab1/Topbp1/Taf15/Prkaa1/Cdk5/Prkab2/Dyrk2/Nuak1/Rfc5/Bard1/Mapk11/Brca1/Tpx2/Taf1/Chek1/Cdk2/Brip1/Aurka/Mdm4/Aurkb/Ccna2/Exo1 |
| R-MMU-8856688 | Golgi-to-ER retrograde transport | 116 | -0.544397828 | -1.621979221 | 0.000891142 | 0.020541568 | 0.018697279 | 2150 | tags=27%, list=14%, signal=23% | Dync1li2/Dync1li1/Kif3b/Kif26b/Copz2/Copg1/Rab3gap2/Zw10/Kif13b/Kif15/Dynll1/Bnip1/Kif1b/Kif18b/Tuba1a/Kif22/Tubb2a/Dync1h1/Kif21b/Tubb2b/Tubb3/Bicd1/Kif18a/Cenpe/Kif23/Racgap1/Kif2c/Kif4/Kif11/Kif20b/Kif20a |
| R-MMU-425410 | Metal ion SLC transporters | 18 | 0.798090153 | 1.793112251 | 0.000966147 | 0.021899332 | 0.019933138 | 1128 | tags=22%, list=7%, signal=21% | Heph/Slc39a7/Slc41a1/Slc40a1 |
| R-MMU-5693579 | Homologous DNA Pairing and Strand Exchange | 25 | -0.743154231 | -1.742667755 | 0.001138261 | 0.025377631 | 0.023099144 | 725 | tags=28%, list=5%, signal=27% | Bard1/Brca1/Chek1/Brip1/Rad51ap1/Rad51/Exo1 |
| R-MMU-159236 | Transport of Mature mRNA derived from an Intron-Containing Transcript | 67 | -0.610655406 | -1.685662996 | 0.001183211 | 0.025954312 | 0.023624049 | 2355 | tags=43%, list=16%, signal=37% | Nup88/Nup54/Nup160/Nup85/Nup42/Nup214/Nup35/Zc3h11a/Cdc40/Rbm8a/Ncbp1/Sarnp/Ddx39b/Nup98/Nup133/Nxf1/Nup155/U2af2/Ranbp2/Tpr/Thoc2/Srsf1/Srsf2/Srsf11/Ddx39a/Srsf7/Nup107/Fyttd1/Srsf3 |
| R-MMU-425366 | Transport of bile salts and organic acids, metal ions and amine compounds | 43 | 0.682460611 | 1.818001864 | 0.001345671 | 0.028155578 | 0.025627678 | 1128 | tags=19%, list=7%, signal=17% | Heph/Emb/Slc39a7/Slc22a18/Slc41a1/Slc6a15/Slc44a2/Slc40a1 |
| R-MMU-156827 | L13a-mediated translational silencing of Ceruloplasmin expression | 108 | 0.56790153 | 1.784443738 | 0.001322085 | 0.028155578 | 0.025627678 | 3057 | tags=59%, list=20%, signal=48% | Rps19/Rps27rt/Rps15/Rps5/Rps27l/Eif3f/Rpl15/Rps14/Rpl19/Rpl28/Rpl18a/Rpl9/Rpl37/Rps11/Rpl32/Rps17/Rps21/Eif3k/Rpl29/Rps13/Eif3h/Rps28/Rpl24/Rpl38/Rps10/Rpl7a/Rps7/Rpl26/Rpl36/Rpl23a/Rplp2/Eif3d/Rps2/Rpl13a/Rps16/Rplp0/Rpl39/Rpl18/Rpl13/Rpl14/Uba52/Rpl6/Rpl35a/Rps18/Rps24/Rps20/Rps29/Rpl22/Rps12/Rps25/Rpl34/Fau/Rps4x/Rpl17/Eif1ax/Rpl3/Rps6/Rps27/Rpl27/Rpl37a/Rpl7/Eif4a2/Rpl8/Rpl21 |
| R-MMU-6804116 | TP53 Regulates Transcription of Genes Involved in G1 Cell Cycle Arrest | 10 | -0.864318148 | -1.648062906 | 0.001334586 | 0.028155578 | 0.025627678 | 663 | tags=40%, list=4%, signal=38% | E2f7/Cdk2/Ccna2/E2f8 |

| | | | | | | | | | | |
|---|---|---|---|---|---|---|---|---|---|---|
| R-MMU-202733 | Cell surface interactions at the vascular wall | 81 | 0.593095201 | 1.77522849 | 0.001516073 | 0.030832363 | 0.028064132 | 621 | tags=21%, list=4%, signal=20% | Itgb2/Fcer1g/Pf4/Cd74/Slc7a8/Angpt2/Sdc1/Atp1b2/Gas6/F11r/Jam2/Cav1/Col1a2/Procr/Gpc1/Sdc2/Hras |
| R-MMU-140877 | Formation of Fibrin Clot (Clotting Cascade) | 16 | 0.802626234 | 1.760882911 | 0.001531479 | 0.030832363 | 0.028064132 | 577 | tags=31%, list=4%, signal=30% | Serping1/F13a1/Pf4/Prcp/Procr |
| R-MMU-2022857 | Keratan sulfate degradation | 10 | 0.862832212 | 1.742854945 | 0.001541618 | 0.030832363 | 0.028064132 | 1271 | tags=50%, list=8%, signal=46% | Lum/Glb1/Galns/Gns/Hexb |
| R-MMU-9018678 | Biosynthesis of specialized proresolving mediators (SPMs) | 9 | 0.903147063 | 1.7740428 | 0.00172155 | 0.033843213 | 0.030804658 | 56 | tags=33%, list=0%, signal=33% | Alox5ap/Ptgs2/Cyp1a1 |
| R-MMU-9711123 | Cellular response to chemical stress | 129 | 0.528171805 | 1.700118444 | 0.00174193 | 0.033843213 | 0.030804658 | 2549 | tags=40%, list=17%, signal=34% | Cybb/Ncf1/Blvrb/mt-Co1/Hmox1/Cyba/Prdx5/Cox8a/Ncoa1/Gstp2/Cat/Prkaa2/Sod1/Sesn1/Cox6a1/Prdx1/Sod3/Cox4i1/Psma1/Prkci/Rxra/Srxn1/Psmc1/Txn1/Cox6b1/Gsr/Txnrd1/Akt1/Ubb/Gpx3/Med1/Psmb3/Cox6c/Chd9/Psmb2/Cox7a2/Psma6/Skp1/Prdx3/Psmb1/Cox7c/Uba52/Cox7a2l/Sin3b/Psmb6/Ubc/Psmd3/Psmb4/Psmb7/Cox5a/Txn2/Ccs |
| R-MMU-72695 | Formation of the ternary complex, and subsequently, the 43S complex | 52 | 0.637662955 | 1.763335288 | 0.001771097 | 0.03392524 | 0.03087932 | 2717 | tags=58%, list=18%, signal=47% | Rps19/Rps27rt/Rps15/Rps5/Rps27l/Eif3f/Rps14/Rps11/Rps17/Rps21/Eif3k/Rps13/Eif3h/Rps28/Rps10/Rps7/Eif3d/Rps2/Rps16/Rps18/Rps24/Rps20/Rps29/Rps12/Rps25/Fau/Rps4x/Eif1ax/Rps6/Rps27 |
| R-MMU-5693537 | Resolution of D-Loop Structures | 33 | -0.700645469 | -1.713446619 | 0.001855579 | 0.0345697 | 0.031465919 | 1688 | tags=30%, list=11%, signal=27% | Blm/Eme1/Fignl1/Bard1/Brca1/Brip1/Rad51ap1/Rad51/Exo1/Gen1 |
| R-MMU-5693568 | Resolution of D-loop Structures through Holliday Junction Intermediates | 33 | -0.700645469 | -1.713446619 | 0.001855579 | 0.0345697 | 0.031465919 | 1688 | tags=30%, list=11%, signal=27% | Blm/Eme1/Fignl1/Bard1/Brca1/Brip1/Rad51ap1/Rad51/Exo1/Gen1 |
| R-MMU-176408 | Regulation of APC/C activators between G1/S and early anaphase | 35 | -0.683165987 | -1.699006128 | 0.001964022 | 0.036095547 | 0.03285477 | 1354 | tags=31%, list=9%, signal=29% | Cdk1/Anapc1/Ccnb1/Anapc4/Cdk2/Plk1/Fbxo5/Bub1b/Ube2c/Ccna2/Cdc20 |
| R-MMU-174143 | APC/C-mediated degradation of cell cycle proteins | 73 | -0.576498982 | -1.598473602 | 0.002116779 | 0.037879204 | 0.034478285 | 414 | tags=19%, list=3%, signal=19% | Cdk1/Anapc1/Ccnb1/Anapc4/Cdk2/Plk1/Aurka/Fbxo5/Bub1b/Ube2c/Aurkb/Ccna2/Nek2/Cdc20 |
| R-MMU-453276 | Regulation of mitotic cell cycle | 73 | -0.576498982 | -1.598473602 | 0.002116779 | 0.037879204 | 0.034478285 | 414 | tags=19%, list=3%, signal=19% | Cdk1/Anapc1/Ccnb1/Anapc4/Cdk2/Plk1/Aurka/Fbxo5/Bub1b/Ube2c/Aurkb/Ccna2/Nek2/Cdc20 |
| R-MMU-5685942 | HDR through Homologous Recombination (HRR) | 56 | -0.626731174 | -1.67968888 | 0.00221566 | 0.038632016 | 0.035163506 | 1688 | tags=29%, list=11%, signal=25% | Blm/Pold1/Eme1/Fignl1/Pcna/Rfc5/Bard1/Brca1/Chek1/Brip1/Pole2/Rad51ap1/Pole/Rad51/Exo1/Gen1 |

| | | | | | | | | | | |
|---|---|---|---|---|---|---|---|---|---|---|
| R-MMU-69478 | G2/M DNA replication checkpoint | 7 | -0.902783036 | -1.635927761 | 0.002192079 | 0.038632016 | 0.035163506 | 1354 | tags=71%, list=9%, signal=65% | Cdk1/Ccnb1/Ccna2/Pkmyt1/Ccnb2 |
| R-MMU-72613 | Eukaryotic Translation Initiation | 116 | 0.538314724 | 1.695666436 | 0.002311157 | 0.039289661 | 0.035762107 | 3057 | tags=55%, list=20%, signal=44% | Rps19/Rps27rt/Rps15/Rps5/Rps27l/Eif3f/Rpl15/Rps14/Rpl19/Rpl28/Rpl18a/Rpl9/Rpl37/Rps11/Rpl32/Rps17/Rps21/Eif3k/Rpl29/Rps13/Eif3h/Rps28/Rpl24/Rpl38/Rps10/Rpl7a/Rps7/Rpl26/Rpl36/Rpl23a/Rplp2/Eif3d/Rps2/Rpl13a/Rps16/Rplp0/Rpl39/Rpl18/Rpl13/Rpl14/Uba52/Rpl6/Rpl35a/Rps18/Rps24/Rps20/Rps29/Rpl22/Rps12/Rps25/Rpl34/Fau/Rps4x/Rpl17/Eif1ax/Rpl3/Rps6/Rps27/Rpl27/Rpl37a/Rpl7/Eif4a2/Rpl8/Rpl21 |
| R-MMU-72737 | Cap-dependent Translation Initiation | 116 | 0.538314724 | 1.695666436 | 0.002311157 | 0.039289661 | 0.035762107 | 3057 | tags=55%, list=20%, signal=44% | Rps19/Rps27rt/Rps15/Rps5/Rps27l/Eif3f/Rpl15/Rps14/Rpl19/Rpl28/Rpl18a/Rpl9/Rpl37/Rps11/Rpl32/Rps17/Rps21/Eif3k/Rpl29/Rps13/Eif3h/Rps28/Rpl24/Rpl38/Rps10/Rpl7a/Rps7/Rpl26/Rpl36/Rpl23a/Rplp2/Eif3d/Rps2/Rpl13a/Rps16/Rplp0/Rpl39/Rpl18/Rpl13/Rpl14/Uba52/Rpl6/Rpl35a/Rps18/Rps24/Rps20/Rps29/Rpl22/Rps12/Rps25/Rpl34/Fau/Rps4x/Rpl17/Eif1ax/Rpl3/Rps6/Rps27/Rpl27/Rpl37a/Rpl7/Eif4a2/Rpl8/Rpl21 |
| R-MMU-8955332 | Carboxyterminal post-translational modifications of tubulin | 19 | -0.774958411 | -1.735871298 | 0.002462176 | 0.041340237 | 0.037628575 | 738 | tags=37%, list=5%, signal=35% | Ttll5/Tuba1a/Tubb2a/Ttll4/Tubb2b/Tubb3/Ttll3 |
| R-MMU-76002 | Platelet activation, signaling and aggregation | 191 | 0.454094931 | 1.530930341 | 0.002510293 | 0.041634123 | 0.037896075 | 1891 | tags=24%, list=13%, signal=21% | Serping1/Fcer1g/Itga2b/F13a1/Ecm1/Pf4/Plek/Qsox1/Gas6/Timp3/Cd9/Dgki/Pdpn/Dgkd/Col1a2/Psap/Sod1/Ptk2/Calm3/Cd36/Gnb2/Itpr2/Tmsb4x/Vav2/Lamp2/Srgn/Gna11/Gnai2/Vegfb/Pros1/Mapk3/App/Prkce/Rap1a/Prkca/Akt1/Fam3c/Itgb3/Gnb1/Aplp2/Rhob/Gng10/Gna12/Itpr3/Lyn/Actn4 |
| R-MMU-453279 | Mitotic G1 phase and G1/S transition | 116 | -0.51725894 | -1.541121602 | 0.00273676 | 0.044843298 | 0.04081712 | 1940 | tags=21%, list=13%, signal=18% | Ppp2r1b/Psmd12/Orc2/Pola2/Skp2/Pola1/Orc1/Akt3/Mcm8/Mcm6/Mcm4/Cdt1/Cks1b/Ccnd1/Mcm3/Mcm10/Cdk2/Pole2/Ccna2/Mcm5/Cdc45/Pole/Cdc6/E2f2 |
| R-MMU-72202 | Transport of Mature Transcript to Cytoplasm | 76 | -0.577253785 | -1.614463929 | 0.0028244 | 0.04526654 | 0.041202362 | 3050 | tags=42%, list=20%, signal=34% | Sympk/Nup88/Nup54/Nup160/Cpsf1/Nup85/Nup42/Nup214/Nup35/Zc3h11a/Slbp/Cdc40/Rbm8a/Ncbp1/Sarnp/Ddx39b/Nup98/Nup133/Nxf1/Nup155/U2af2/Ranbp2/Tpr/Thoc2/Srsf1/Srsf2/Srsf11/Ddx39a/Srsf7/Nup107/Fyttd1/Srsf3 |
| R-MMU-380320 | Recruitment of NuMA to mitotic centrosomes | 82 | -0.563875469 | -1.589272451 | 0.002829159 | 0.04526654 | 0.041202362 | 1448 | tags=33%, list=10%, signal=30% | Tubg2/Alms1/Numa1/Cdk5rap2/Ckap5/Hsp90aa1/Cdk1/Clasp1/Tubb5/Dynll1/Sfi1/Tubgcp4/Akap9/Pcnt/Tuba1a/Pcm1/Cep1 |

| | | | | | | | | | | |
|---|---|---|---|---|---|---|---|---|---|---|
| | | | | | | | | | | 52/Cpap/Cep192/Tubb2a/Dync1h1/Tubb2b/Cep135/Plk1/Tubb3/Cep290/Nek2 |
| R-MMU-69206 | G1/S Transition | 93 | -0.546775214 | -1.581584458 | 0.002899386 | 0.045850763 | 0.041734131 | 1940 | tags=25%, list=13%, signal=22% | Ppp2r1b/Psmd12/Orc2/Pola2/Skp2/Pola1/Orc1/Akt3/Mcm8/Mcm6/Mcm4/Cdt1/Cks1b/Ccnd1/Mcm3/Mcm10/Cdk2/Pole2/Ccna2/Mcm5/Cdc45/Pole/Cdc6 |

**Table S4.** qRT-PCR sequences for used primers

| Gene | Gene name | Direction | Primers sequence (5' to 3') |
|---|---|---|---|
| **Foxm1** | Forkhead box protein M1 | F | AGCGTTAAGCAGGAACTGGA |
| | | R | GGAAGTGGTCCTCAATCCAA |
| **Birc5** | Baculoviral IAP repeat-containing 5 (Survivin) | F | GAGGCTGGCTTCATCCACTG |
| | | R | CTTTTTGCTTGTTGTTGGTCTCC |
| **Abcb1a** | ATP-binding cassette sub-family B member 1 | F | ACACTTGGCCCCAAACATAGA |
| | | R | GTCAATGCTTGGCTCGTTATCA |
| **Il-6** | Interleukin-6 | F | CTGCAAGAGACTTCCATCCAG |
| | | R | AGTGGTATAGACAGGTCTGTTGG |
| **Ptgs2** | Prostaglandin-endoperoxide synthase 2 (COX-2) | F | TTCCAATCCATGTCAAAACCGT |
| | | R | AGTCCGGGTACAGTCACACTT |
| **Prdm1** | PR/SET domain 1 | F | CTTCTCTTGGAAAAACGTGTGGG |
| | | R | TCATATCAGCGTCCTCCATGT |
| **c-Myc** | MYC proto-oncogene | F | ATGCCCCTCAACGTGAACTTC |
| | | R | GTCGCAGATGAAATAGGGCTG |
| **Pbk** | PDZ-binding kinase | F | TGGGCCGTGAAAAAGATAAGTC |
| | | R | CTGGCTTCAGTAAAAGCACGATA |
| **Tk1** | Thymidine kinase 1 | F | AGTGCCTGGTCATCAAGTATGC |
| | | R | GCTGCCACAATTACTGTCTTGC |
| **Ncapg2** | Non-SMC condensin II complex subunit G2 | F | GATCCTTATCCACGGGTTCGT |
| | | R | TCGGCTGAGCTTATATCAAATGC |
| **Tyms** | Thymidylate synthetase | F | GGAAGGGTGTTTTGGAGGAGT |
| | | R | GCTGTCCAGAAAATCTCGGGA |
| **Siva1** | SIVA1 apoptosis-inducing factor | F | CCGCTCCAACTCAAAGTCCA |
| | | R | CGGAACAATCTTCGCTCGATATG |
| **cGas** | Cyclic GMP-AMP synthase | F | CAGGAAGGAACCGGACAAGC |
| | | R | CCGACTCCCGTTTCTGCATT |
| **Pdss1** | Decaprenyl diphosphate synthase subunit 1 | F | ACACCAGCAATGTGCAGTTG |

| | | | |
|---|---|---|---|
| | | R | ACAGACCTTTCAAGTCTCTCCAG |
| **Etv4** | ETS variant transcription factor 4 | F | CGGAGGATGAAAGGCGGATAC |
| | | R | TCTTGGAAGTGACTGAGGTCC |
| **Tfap4** | Transcription factor AP-4 | F | CTCTGTAGCCTAGCCAACATTC |
| | | R | GAAGCCCGCGTTGATACTCT |
| **Topbp1** | Topoisomerase II-beta binding protein 1 | F | CAGGATTGTTGGTCCTCAAGTG |
| | | R | ACAGGATACAGTTACGTCAGACA |
| **Cxcl12** | C-X-C motif chemokine ligand 12 | F | TGCATCAGTGACGGTAAACCA |
| | | R | CACAGTTTGGAGTGTTGAGGAT |
| **Lum** | Lumican | F | CTCTTGCCTTGGCATTAGTCG |
| | | R | GGTCATCACAGTACATGGCAGT |
| **Medag** | Mesenteric estrogen-dependent adipogenesis protein | F | CTTGTGCGCCTAGAAGGGC |
| | | R | TGCTCAGTATCGTTTCCCTGTA |
| **Mki67** | Marker of proliferation Ki-67 | F | ATCATTGACCGCTCCTTTAGGT |
| | | R | GCTCGCCTTGATGGTTCCT |
| **Foxf2** | Forkhead box F2 transcription factor | F | CGTCCTCTTCTAACTCCGTCA |
| | | R | ATGTACGAGTAAGGAGGCTTCT |
| **Lrr1** | Leucine-rich repeat protein 1 | F | GCCAACCTTAGGGCTCAAGAG |
| | | R | AGGCTCCTTTAGCCGGACA |
| **Slc29a2** | Solute carrier family 29 member 2 | F | ACTTCAACAACTGGGTGACAC |
| | | R | GGGGATGCACTGATACAGGAA |
| **Cenpf** | Centromere protein F | F | AAGACGGAAAGACTGAGGGTG |
| | | R | GATTGTGCAACTTGTTGGCTTC |
